# Validation and analysis of 12,000 AI-driven CAR-T designs in the *Bits to Binders* competition

**DOI:** 10.64898/2026.03.03.709355

**Authors:** Clayton W. Kosonocky, Alex M. Abel, Aaron L. Feller, Amanda E. Cifuentes Rieffer, Phillip R. Woolley, Jakub Lála, Daryl R. Barth, Tynan Gardner, Bits to Binders Competitors, Stephen C. Ekker, Andrew D. Ellington, Wesley A. Wierson, Edward M. Marcotte

**Affiliations:** Department of Molecular Biosciences, University of Texas at Austin, Austin, Texas, USA; The BioML Society, Austin, Texas, USA; LifEngine Animal Health (LEAH) Laboratories Incorporated, Minneapolis, Minnesota, USA; Department of Materials, Imperial College London, London, United Kingdom; Department of Pediatrics, Dell Medical School, University of Texas at Austin, Austin, TX, USA

## Abstract

Artificial intelligence (AI) methods for proteins have advanced rapidly, improving structure prediction and design, particularly for *de novo* binders. However, most evaluations emphasize binding affinity rather than higher-order biological function. We present *Bits to Binders*, a global competition benchmarking *de novo* binder design in the context of chimeric antigen receptor (CAR) T cells. Teams from 42 countries submitted 12,000 designs of 80-amino acid binders targeting human CD20 as CAR binding domains. Designs were screened by pooled CAR-T proliferation, identifying 707 designs exhibiting significant CD20-specific enrichment, with team hit rates from 0.6% to 38.4%. Top-performing candidates were validated as individual constructs, measuring CD20-specific proliferation, expansion, cytokine production, and targeted cell lysis. We identified common design methodologies and factors correlated with DNA synthesis, expression, and target-specific T cell activation which nearly double the success rates when applied as a retrospective filter. We release this dataset as an open resource, with practical recommendations to support more effective AI-driven binder design.

## Introduction

The application of artificial intelligence (AI) methods to proteins has shown remarkable success in recent years^1^. One application area, protein binder design, has garnered significant interest due to its demonstrated tractability and potential to create antibody-like therapeutics^2^. Many AI-driven models have been validated on their ability to design proteins that bind to arbitrary targets, with reported success rates exceeding 10% of tested designs. Despite this progress, the best practices for protein binder design are not yet standardized and it is unclear which computational approaches are most effective across diverse contexts.

Regularly occurring benchmarks like the Critical Assessment of Structure Prediction (CASP) and the Critical Assessment of Predicted Interactions (CAPRI) have been essential in advancing protein structure prediction models to their current level of performance^3,4^. Similar efforts are emerging in AI-driven protein design, such as the Protein Engineering Tournament and Adaptyv Bio’s EGFR binder challenge, which validate crowd-sourced designs against measurable objectives such as binding affinity and activity, providing insight into method performance^5,6^. While these competitions have been instrumental in assessing AI-driven tools for designing proteins that bind, the suitability of designs for other functional endpoints remains largely unknown.

Binding is one of several important features of immune function, and engineered binding by Chimeric Antigen Receptor (CAR) T Cells is of great therapeutic import^7^. CAR-T cells are derived from a patient’s normal T cells through introduction and expression of a chimeric T cell receptor, where the extracellular domain enables the recognition of a targeted tumor surface antigen^8,9^. Upon target engagement by that antigen, receptor clustering occurs, and downstream pathways are activated that drive T cell proliferation, cytokine secretion, and cytotoxic killing.

Prior work has demonstrated the general feasibility of AI-designed CAR binding domains^10–13^, and thus this system provides an excellent opportunity for evaluating different design approaches in a competition setting. Designing binders in the context of a CAR-T cell requires construct expression and membrane localization of the receptor that is compatible with cell viability and engages downstream signaling upon target recognition, leading to proliferation, expansion, cytokine production, and target cell lysis. We created the *Bits to Binders* competition which engaged a total of 28 teams to generate 12,000 protein binders that targeted the lymphoma surface antigen CD20, to be synthesized and assayed for CAR-T cell function. From the resulting dataset, we extracted design principles and predictive metrics that contribute to functional binder performance in the context of a CAR-T cell therapy, with the goal of moving the field closer to standardized and effective methods for AI-driven protein design.

## Results

### Crowd-sourcing *de novo* designed CD20-binding CARs for the *Bits to Binders* competition

Competitors from 42 countries were tasked to use their chosen AI-driven protein design methods to design 500 80-amino-acid binders to the extracellular domain of human CD20 that could be accommodated in the context of a CAR (**Fig. 1**). This task differed from a pure binder design task in several ways. First, while the competitors submitted only binding domains, they were aware that these should eventually be compatible with the greater CAR scaffold, in particular the transmembrane domain. Second, success was dependent on achieving sufficient affinity to effectively recognize the target, but not so much that the CAR-T cell becomes exhausted^14,15^. And finally, success would ultimately depend on CAR-T function, not just binding. These nuances, along with a primer on CAR-T biology, the exact CAR scaffold used, and an in-depth explanation of all assays were conveyed to the participants during the solicitation (**Supplementary Information C**). In addition, several research and review articles on AI-driven protein design, CAR-T therapies, and CD20 biology were provided to the participants (**Supplementary Information C**).

**Figure 1.**
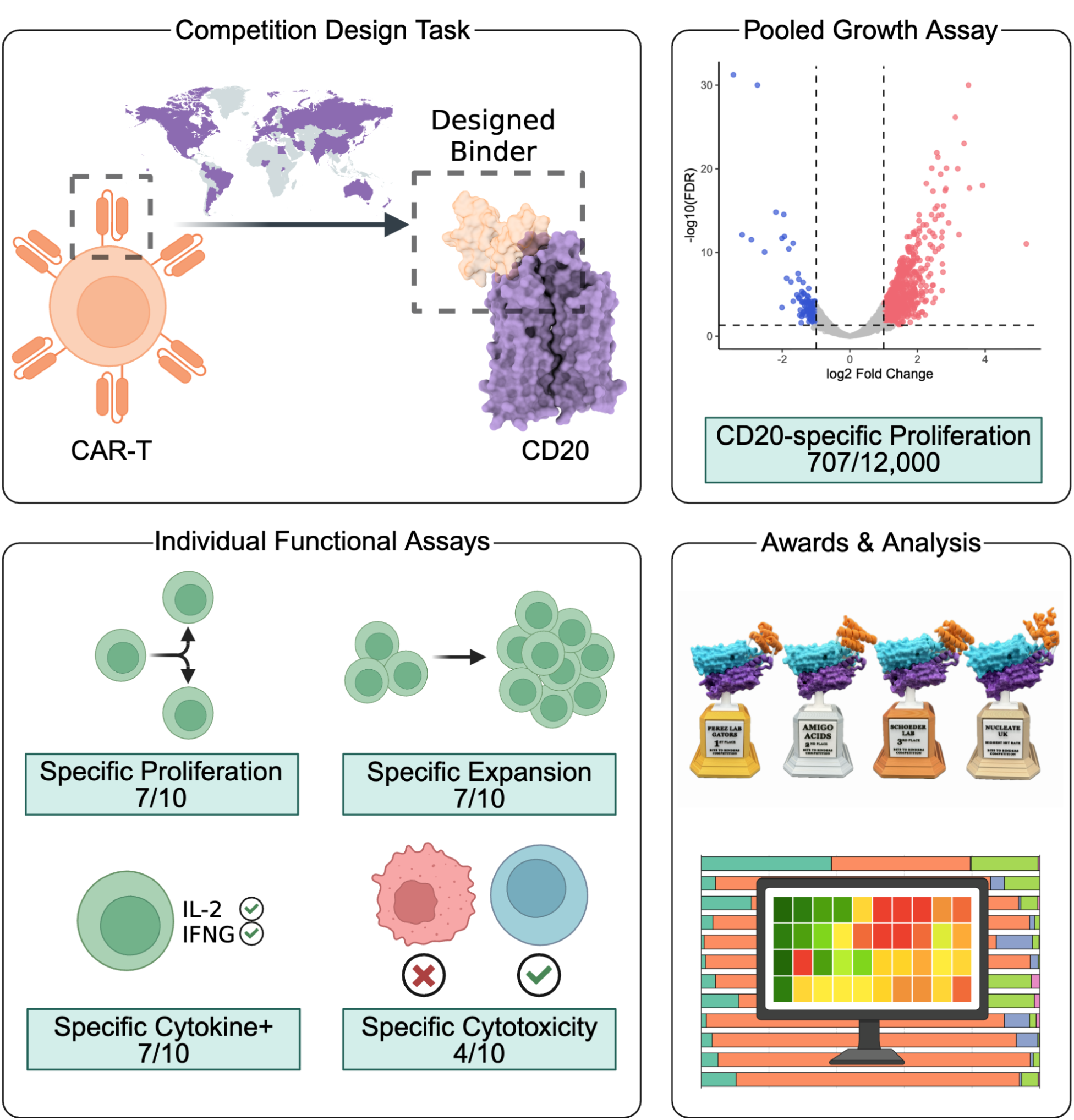
Overview of the *Bits to Binders* competition. Competitors from around the world used AI-driven protein design methods to design 12,000 proteins that bind CD20 and initiate a T cell immune response when used as the binding domain of a CAR-T cell. The 12,000 designs were tested in a competitive growth assay, in which CD20-specific CAR-T cell proliferation was measured. The designs with the greatest CD20-specific fold change in growth were tested individually for CD20-specific proliferation, expansion, cytokine production, and target cell lysis. The results were then analyzed to identify computational features leading to increased success rates for AI-driven CAR design. Figure 1 was created using BioRender (https://biorender.com).

At the end of the six-week design stage, a total of 12,000 designs were collected. The approaches used spanned a wide range of generative design and selection strategies (**Supplementary Information E**). Generative diffusion models were the most common approach, with 18 teams using RFdiffusion^16^ and two teams using sequence-structure co-design models like Chroma^17^. The non-diffusion approaches included constraint- or hallucination-based design such as BindCraft^18^ (3 teams), EvoBind^19^ (2), and ColabDesign (3), as well as protein language models like PepMLM^20^ (1), ESM-2^21^ (2), and RayGun^22^ (1). Fixed backbone design tools were widely used for sequence generation and diversification, notably ProteinMPNN^23^ (19 teams) followed by SolubleMPNN^24^ (2) and CARBonAra^25^ (1). Twelve teams generally followed the default RFdiffusion–ProteinMPNN–AlphaFold2 pipeline, while nine teams opted to perform “iterative diffusion” in which the initial binder was generated, the low-scoring segments were removed, and the process was repeated until the scores were sufficient.

Many teams conditioned their generative models on CD20 in the Rituximab-bound state (PDB: 6VJA) or by starting with fragments of CD20-specific antibodies. Initial structures were frequently refined using FastRelax^26^, GROMACS^27^, or OpenMM^28^. After generation, it was common to select sequences using scores, structure prediction models, or other bioinformatics tools. To confirm that generated sequences were predicted to fold into the desired structures, AlphaFold2^29^ was used by 17 teams, with others using ColabFold^30^ (4), ESMFold^21^ (4), Chai-1^31^ (1), or HelixFold^32^ (1). Interestingly, only one team reported folding their designs in the context of the entire CAR sequence. Other selection criteria included Rosetta energy scores^33,34^, HDOCK scores^35^, and binding affinity predictions from Prodigy^36^. The wide variety of inputs, model types, and scoring strategies resulted in substantial sequence diversity between teams: over 83% of submissions did not produce a significant MMseqs2^37^ alignment to sequences from other teams (**Fig. SA1**). In contrast, within-team sequence redundancy was relatively high, with 20% of sequences aligning nearly perfectly to another design from the same team (≥0.95 MMseqs2 identity).

### Screening 12,000 *de novo* CAR binding domains in a T cell proliferation assay

To functionally evaluate the 12,000 binder designs we employed a pooled CAR-T challenge assay that measures antigen-dependent T cell proliferation (**Fig. 2A**). Pools of codon-optimized DNA fragments encoding the 80-amino-acid binding domain were synthesized and cloned into a plasmid encoding a second-generation CAR backbone by replacing the binder region of a second-gen CAR28z construct (**Methods**). The plasmid library was used as a DNA donor for targeted integration of the CAR at the T cell receptor alpha chain (TRAC) to engineer a pool of TRAC(-), CAR(+) T cells^38,39^.

**Figure 2.**
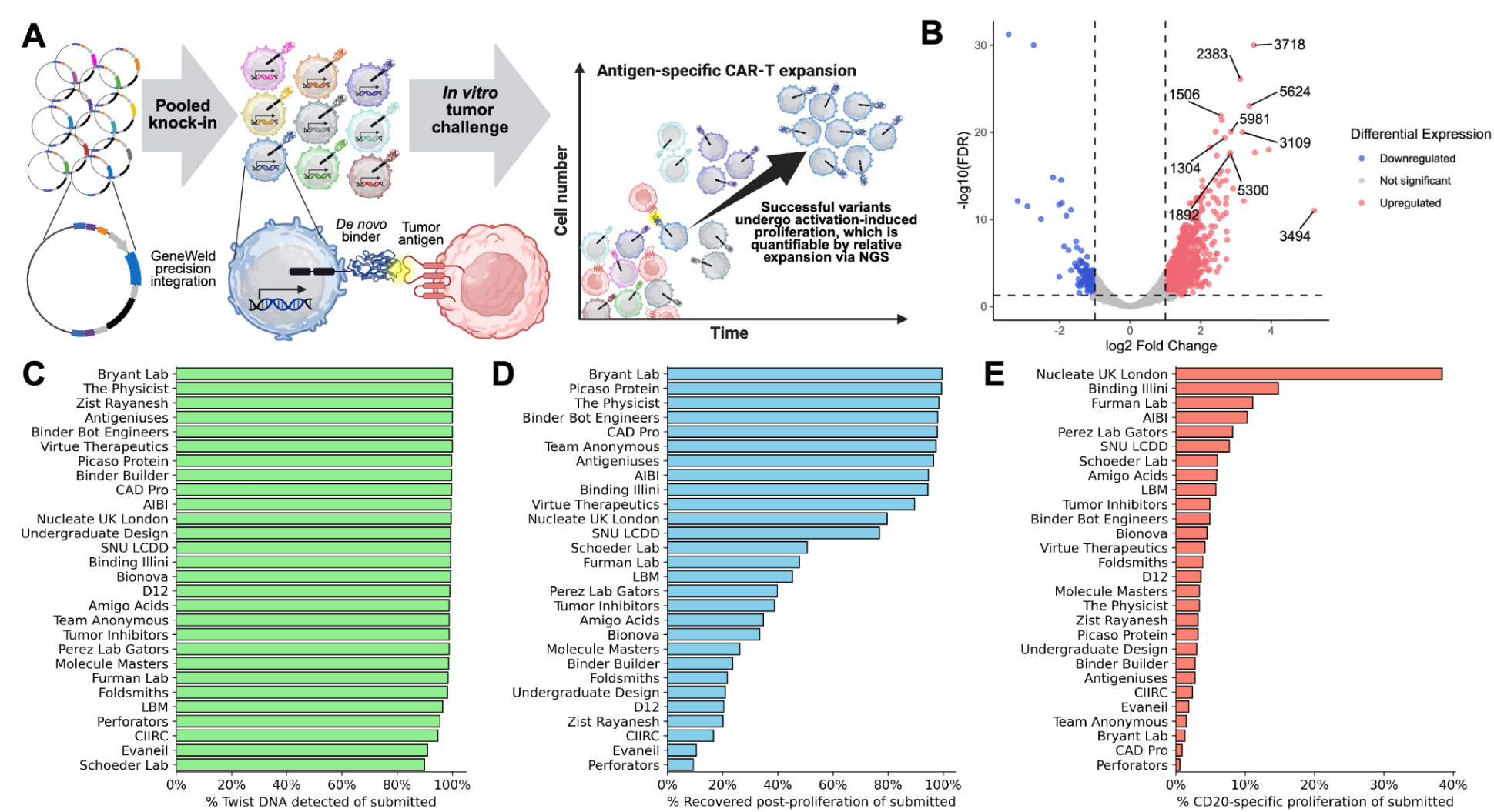
Pooled proliferation assay identifies designs with CD20-specific CAR-T cell response. **(A)** CD20 binder transgenes are knocked into a population of T cells, which is then subjected to either a tumor challenge or a control condition. Target-specific T cell proliferation indicates antigen recognition and is quantified as the relative change in abundance of each transgene sequence in the tumor challenge condition compared with the control. **(B)** Of 12,000 binders, 707 showed a significant CD20-specific ≥2-fold increase in growth, indicating potential recognition of CD20 by the binder. **(C)** Percent of each team’s submitted sequences detected in the plasmid pool during Twist quality control. **(D)** Percent of each team’s submitted sequences with ≥25 reads in each of the six proliferation assay replicates. Post-proliferation recovery suggests that the binder produced a viable CAR-T cell. **(E)** Percent of each team’s submitted sequences with significant ≥2-fold change in CD20-specific proliferation. Panel A was created using BioRender (https://biorender.com).

The sequence of the integrated, designed binder in each CAR construct served as its own unique DNA barcode for each cell. To enrich for T cells carrying integrations of the pooled CAR library and negatively select unedited T cells, we subjected the population to methotrexate selection^40,41^ via the co-expressed DHFR^F/S^ variant. After selection and expansion, the remaining CAR-expressing cells were co-cultured either alone or with the CD20-positive Raji cell line^42^, both in triplicate. Enrichment of binders was measured by the relative change in “barcode” sequences after two weeks in Raji co-culture compared to those that were cultured without CD20+ targets. Enrichment or depletion in the non-target group may be indicative of dysregulation in CAR-T signaling or decreased cellular fitness independent of binding. Thus, comparing the Raji co-culture with this group allows the effect of CD20+ targets on proliferation to be robustly quantified.

NGS reads were quantified across the three technical replicates to identify significant CD20-specific enrichment (**Fig. 2B, Methods**), with few jackpot effects occurring between the replicates (**Fig. SA2**). Across the 12,000 designs, 98.3% passed Twist’s DNA quality check (**Fig. 2C**), but only 56.8% passed the minimum read thresholds after growth. Given that a functional CAR is required to provide the basal signaling required for sustained T cell fitness and survival, this drop-off indicates that many of the designs may have failed to yield a valid receptor (**Fig. 2D**).

Ultimately, 707 clones showed significant CD20-specific enrichment of at least twofold relative to controls. Hit rates varied substantially between teams, ranging from 0.6% to 38.4% of submitted sequences (**Fig. 2E**). The top ten non-redundant designs were chosen to proceed to a series of individual T cell functional assays.

### Confirming function with individual T cell assays

Each of the top ten binders were cloned as individual CAR constructs in a manner identical to how they would have appeared in the pooled library. CAR-T cells expressing each of these binders were characterized in a series of assays probing different aspects of T cell activation and effector function, including assays for expression, proliferation, expansion, cytokine production, and target cell lysis (**Fig. 3A**). CD20-targeting, scFv-based CAR-T and untransfected TCR+ T cells served as positive and negative controls, respectively.

**Figure 3.**
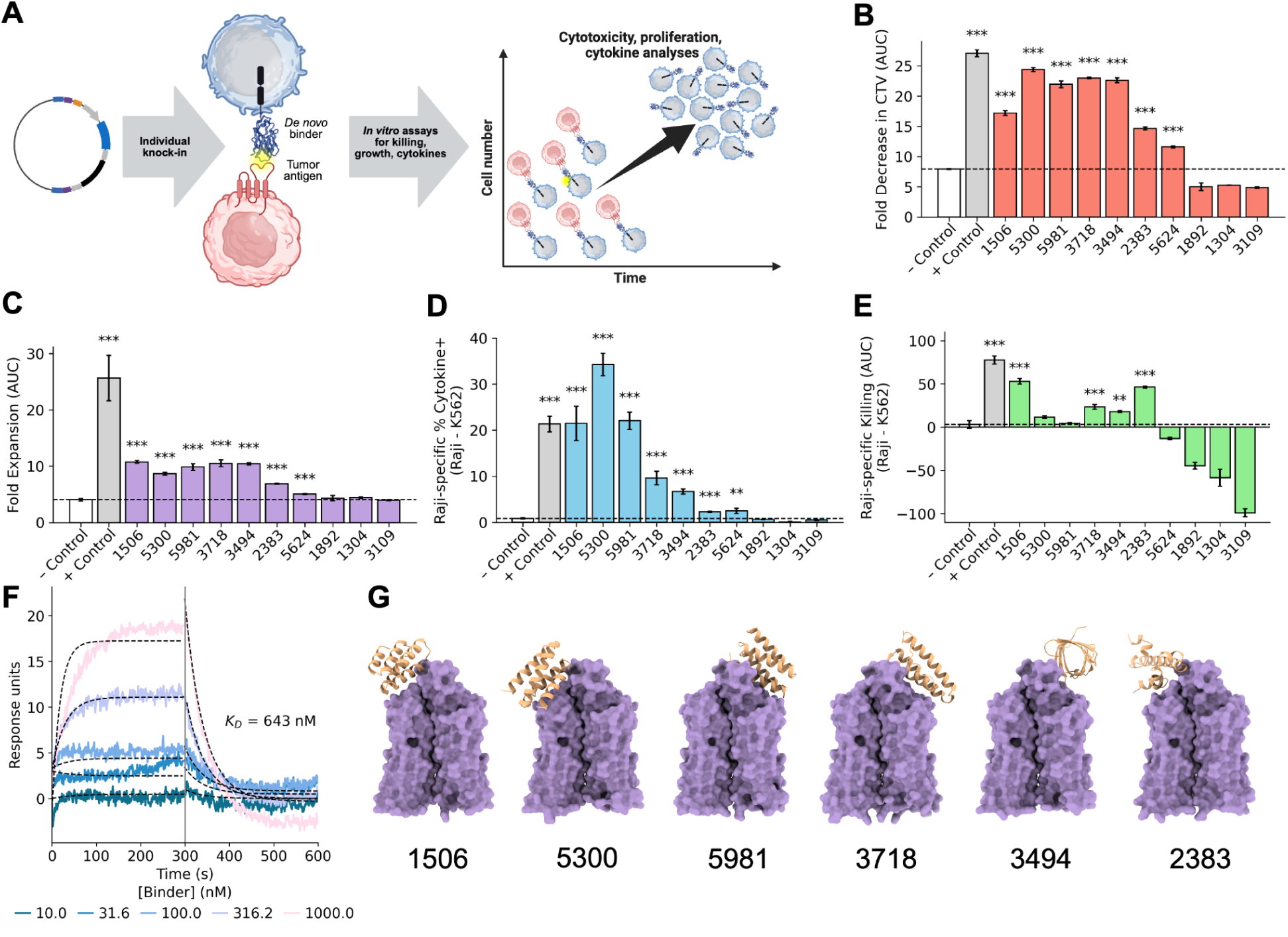
*In vitro* assays confirm designed binders function as CAR-T cells. **(A)** DNA constructs of designed binders are integrated into primary T cells for functional analysis against CD20+ tumor cells. **(B)** CD20-specific proliferation, computed by AUC of CTV dilution (Raji − K562) across four E:T ratios (**Fig. SA4**). Bars show mean CTV MFI ± SE; significance by BH-corrected one-sided Welch’s t-test vs. negative control. **(C)** CD20-specific expansion, computed as AUC of fold-expansion (Raji − K562) across four E:T ratios (**Fig. SA6**). Bars show mean ± SE; BH-corrected one-sided Welch’s t-test vs. negative control. **(D)** CD20-specific production of IL-2 and IFNγ. Bars show mean percent of cytokine+ cells in the vs-Raji condition ± SE; BH-corrected one-sided Welch’s t-test vs. negative control. **(E)** CD20-specific cytotoxicity, computed as AUC (Raji − K562) across three E:T ratios (**Fig. SA8**). Bars show mean ± SE; BH-corrected one-sided z-test vs. negative control. **(F)** SPR measurements for design 2383 indicate moderate binding affinity to detergent-solubilized CD20 when expressed as an isolated 80-mer (*K*_D_ = 643nM). **(G)** Boltz-1 structures of selected CD20-bound designs that broadly outperformed the negative control. Significance stars relative to negative controls: *, p < 0.05; **, p < 0.01; ***, p < 0.001. Panel A was created using BioRender (https://biorender.com).

Flow cytometry was used to immunophenotype TCR and CAR expression (**Fig. SA3A, SA3B**). Seven of the designs (1506, 1892, 3494, 3718, 5300, 5624, 5981) proved to be CAR+ cells, with 10-50% of the population being CAR+ cells (**Fig. SA3D**). Design 2383 had <5% CAR+ cells, while 1304 and 3109 all but lacked expression (**Fig. SA3D**).

CAR-T cell cumulative expansion and cell division were evaluated five days after stimulation with CD20-positive target cells at varying effector-to-target ratios. Seven clones outperformed the CAR-negative untransfected control T cells in expansion and proliferation (**Fig. 3B, 3C, SA4, SA5, SA6**; clones 1506, 2383, 3494, 3718, 5300, 5624, 5981). The remaining three designs (1304, 1892, 3109) underperformed or were equal to the negative control.

Target-specific production of the cytokines IL-2 and IFNγ was assessed with flow cytometry, and the same seven designs outperformed the negative control when cultured with CD20+ targets (**Fig. 3D, SA7**). Clones 1506, 5300, and 5981 showed the most activity, generating similar, if not more, cytokines than the scFv control. The remaining three designs (1304, 1892, 3109) once again did not outperform the negative control.

Finally, specific lysis of CD20+ target cells over 48 hours was evaluated by co-incubating CAR-T cells with CD20-positive target cells (Raji-GFP) and CD20-negative controls (K562-GFP) at varying effector-to-target ratios. Although the same seven designs outperformed the negative control in terms of non-specific cytotoxicity, lysing both CD20+ and CD20-target cell lines at the measured E:T ratios, only four of these (1506, 2383, 3494, 3718) had significant, specific lysis of CD20+ cell lines above the negative control (**Fig. 3E, SA8**).

### Characterization of binding

Activation in the presence of CD20+ Raji cells implies that there were productive interactions with the designed binders, so we further assessed binding via a surface plasmon resonance assay in which detergent-solubilized human CD20 was flowed across the immobilized 80-amino-acid designs. The known scFv was used as a positive control and was found to have a *K_D_* of 1.75nM (**Fig. SA9**). Of the top ten binders, only three were found to have appreciable binding in this assay: 2383 had a *K_D_* of 643nM and designs 1506 and 5981 had detectable binding but weak signal, estimated by Anabel^43^ to be between 200-600nM (**Fig. 3F, Fig. SA9**). This is perhaps unsurprising, given that optimal CAR function often occurs within a moderate range of binding constants; if binding is too high, trogocytosis and T cell exhaustion can occur and many lower affinity CARs have been found to elicit good if not better tumor elimination ^44,45^.

### Competition winners and outcomes of methodological choices

Overall, design 2383 from Nucleate UK London bound strongly to CD20, had high specific lysis, moderate proliferation and expansion, low cytokine %, and low CAR+ %. Design 1506 from Perez Lab Gators had weaker binding but high specific lysis, cytokine %, proliferation, expansion, and CAR+%. Design 5981 from the Schoeder Lab also had weaker binding and high proliferation, expansion, and CAR+%, but had strong lysis with weak specificity. Designs 3494, 3718, 5300, and 5624 (from SNU LCDD, SNU LCDD, Amigo Acids, and Binding Illini, respectively) had no measured binding despite being broadly functional, with two of the four having target-specific lysis, and all four having overall lysis, suggesting that these designs may have largely relied on avidity effects or mechanisms other than CD20 recognition. Designs 1304 and 3109 did not express, while design 1892 expressed but failed all functional assays. One conclusion is thus that 7 of the top 10 designs were successful in functional assays, suggesting that a variety of generative design approaches (from teams Nucleate UK London, Perez Lab Gators, Amigo Acids, Schoeder Lab, SNU LCDD, and Binding Illini) can yield functional CAR-T cells.

The choices made by each team in the design process were numerous and provide ample ground for further methodological analysis. The best-performing generative methods were BAGEL^46^ and Chroma, followed by RFdiffusion, BindCraft, and team AIBI’s sequence-structure co-design model (**Fig. 4A**). Many designs were verified with structure prediction models, with ESMFold in first, then ColabFold and AlphaFold-like models (**Fig. 4B**). However, as ESMFold’s success is carried by its use in BAGEL, the true causal impact of the structure prediction models is unclear. That said, the effect of ProteinMPNN or SolubleMPNN usage is very strong, with the use of these models causing an increase in CD20-specific enrichment and simultaneous lack of detection in the proliferation assay, as further analyzed below (**Fig. 4C, 5B**). Conversely, ESM-2 usage resulted mostly in sequences that were recovered in the proliferation assay (**Fig. 4D**). Using molecular dynamics tools on the structure either before generating to obtain an optimal pose, or afterwards for filtering, correlated with a modest increase in successful designs, though this was not true for FastRelax or AmberRelax (**Fig. 4E-F**).

**Figure 4.**
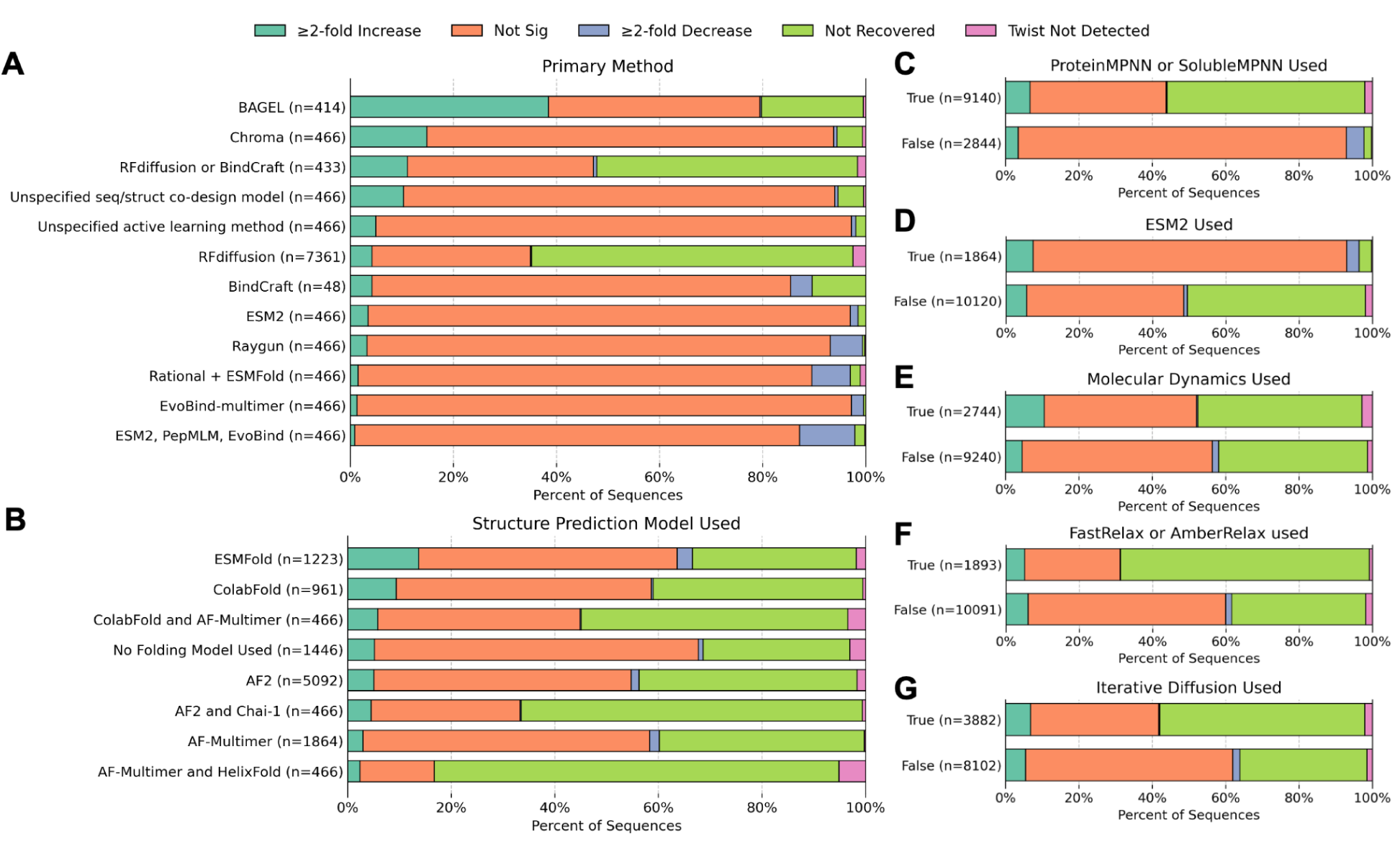
Experimental outcomes based on methodological choices. Methodological choices were categorized into: **(A)** choice of primary generative method, **(B)** choice of structure prediction model, **(C)** the use of ProteinMPNN or SolubleMPNN, **(D)** the use of ESM-2, **(E)** the use of molecular dynamics prior to or after generation, **(F)** the use of FastRelax or AmberRelax prior to or after generation, and **(G)** the use of diffusion models in an iterative manner. Although some trends are visible, the causality is unclear and may result from artifacts of this categorization failing to capture higher order methodological details.

**Figure 5.**
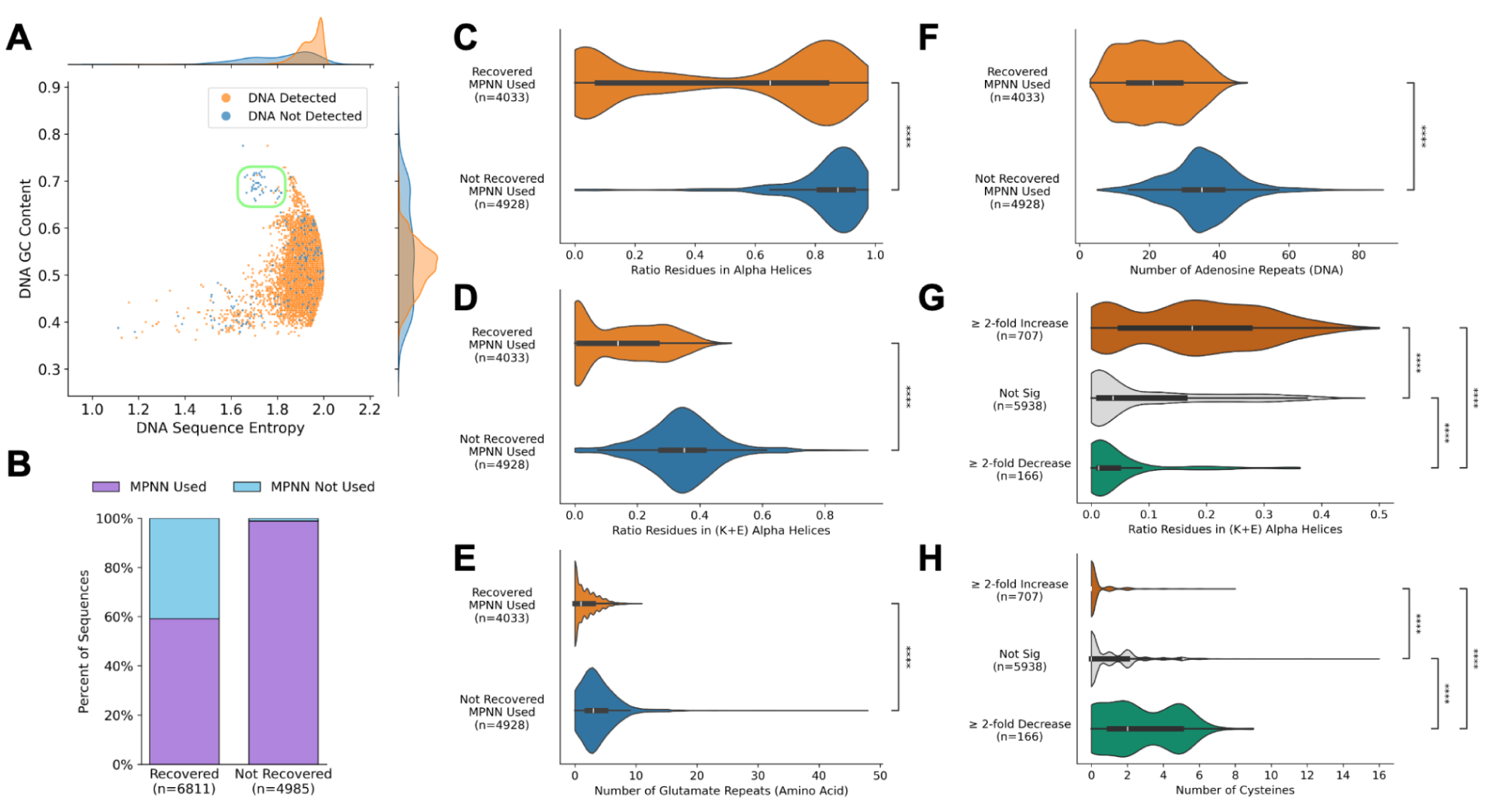
Computational analysis suggests mechanisms leading to success and failure. **(A)** The combination of DNA GC content and Shannon entropy separates out 55 designs that failed to synthesize as DNA. **(B)** Almost all of the designs that failed to be recovered were created with ProteinMPNN or SolubleMPNN. **(C-D)** The MPNN designs that failed to be recovered were enriched in alpha helices that contained mostly lysine and glutamate. **(E-F)** These designs had significant glutamate repeats (EE) in the amino acid sequence and adenosine repeats (AA) in the DNA sequence, both of which have been shown to cause ribosomal termination, suggesting a possible mechanism for the absence of these sequences. **(G)** The designs that caused a ≥2-fold increase in CD20-specific proliferation were enriched in (K+E) alpha helices, though to a lesser extent than those that were not recovered. **(H)** The designs that caused a ≥2-fold decrease in CD20-specific proliferation were enriched in cysteines predicted to form disulfide bonds within the binder chain (**Fig. SA11**). Significance from two-sided BH-corrected Mann-Whitney-Wilcoxon tests (**** for p ≤ 0.0001).

There were common methodologies shared between the top-performing teams (Perez Lab Gators, Amigo Acids, Schoeder Lab, Nucleate UK London, SNU LCDD, Binding Illini, and Furman Lab). First, all of these teams used a diffusion model, either RFdiffusion or Chroma. Second, all of these teams used ProteinMPNN or SolubleMPNN for some stage of the design. Third, many of these teams employed these models in an iterative or refinement strategy, although this was found to have minimal impact on success across all teams (**Fig. 4G**). The teams with the two best-performing binders (Perez Lab Gators, Amigo Acids, **Fig. SA10**) both had highly similar pipelines employing iterative RFdiffusion, ProteinMPNN and FastRelax. Notably, the team with the top-performing design (Perez Lab Gators, **Fig. SA10**) was the only team that reported folding their designs within the greater CAR sequence.

### Analysis of factors leading to design failure and success

The diversity of sequences generated during the competition by various methods are themselves a wealth of information on the success of AI-driven tools, as applied by many different users, for designing proteins to modify cellular functionality. The nucleotide and amino acid sequences of the 12,000 designed binders were used to compute over 400 metrics, ranging from simple readouts like sequence entropy to more involved ones such as the average energy of the binder-CD20 interface on a predicted structure (**Methods; Supplementary Information B**). The varied features were further analyzed for their ability to predict (a) detection in the DNA quality report, (b) whether or not the design was recovered by NGS after proliferation, and (c) the CD20-specific change in proliferation.

The Twist plasmid pool quality report indicated that 204 of the 12,000 designs were not detected. Almost all of the failed designs originated from 5 of the 28 teams (71%, **Fig. 2C**), suggesting that, despite codon optimization, particular design features led to synthesis failures. One common cause seemed to be high GC content and repetitive sequences; a cluster of failed sequences with high GC content and moderately low Shannon entropy can be observed (**Fig. 5A**). Applying a more stringent filter to remove sequences with <1.85 Shannon entropy and ≥0.65 GC content would have removed a quarter of the DNA failures (55 designs), while having no impact on functional sequences.

Of the synthesized designs, 6,811 were recovered by NGS after the proliferation assay (≥25 reads in all six replicates) and 4,985 were not recovered. As a functional CAR is necessary for sustained T cell fitness and survival, failure in this readout suggests failed protein expression, solubility, or folding. The recovered and non-recovered designs differ markedly in their amino acid compositions, with the non-recovered sequences being preferentially enriched in glutamate, lysine, and alanine (**Fig. SA11**).

Strikingly, 98.9% of the non-recovered sequences were designed with ProteinMPNN or SolubleMPNN (MPNN), whereas the recovered sequences came from both these models (59.2%) and other methods (40.8%) (**Fig. 5B**). To determine why some MPNN designs failed where others succeeded, the designs created without MPNN were removed, resulting in 4,033 recovered and 4,928 non-recovered sequences. Categorizing the secondary structure of these designs’ Boltz-1 predicted structures indicated that most of the non-recovered MPNN sequences were almost entirely alpha helices, with an average of 83% of the residues being part of a helix (**Fig. 5C**). Compared to the alpha helices from the recovered sequences, those from the non-recovered sequences were highly enriched in lysine (K) and glutamate (E) (**Fig. 5D**). In fact, the ratio of residues in K+E alpha helices was by far the strongest predictor of sequence recovery, averaging 35% of residues for non-recovered MPNN and 15% for recovered MPNN and achieving 0.91 ROC-AUC using a logistic regression model trained with stratified 5-fold cross validation (**Table SA1, Fig. SA12, Methods**).

We thus formulated hypotheses to explain the mechanism behind the importance of K+E alpha helices. Repeats of acidic amino acids are known to directly impact protein synthesis since “nascent chains enriched in aspartic acid (D) or glutamic acid (E) in their N-terminal regions alter canonical ribosome dynamics, stochastically aborting translation”^47^. There were 3.9 EE repeats in the non-recovered MPNN sequences on average versus 1.6 in the recovered, and this feature achieves 0.82 ROC-AUC (**Fig. 5E**). Similarly, adenosine repeats in the nucleotide sequence have been shown to be potent translational disrupters^48,49^, which may correlate with the use of lysine (AAA, AAG) and glutamate (GAA) codons. The non-recovered sequences were significantly enriched in adenosine repeats in the DNA sequence, with means of 35.4 adenosine dinucleotide repeats in the non-recovered versus 21.7 in the recovered (**Fig. 5F**). The number of AA repeats was one of the most predictive features for sequence recovery, achieving 0.88 ROC-AUC, and also plausibly explaining why DNA sequence entropy alone had high predictive power at 0.90 ROC-AUC (**Table SA1**).

The most likely explanation for these design biases is that ProteinMPNN and SolubleMPNN preferentially predict K and E^50^, a preference which may be further compounded by the fact that RFdiffusion frequently creates alpha helix-rich backbones^51^. That said, sequences that had ≥2-fold increase in proliferation still had a mean of 18% of residues in K+E alpha helices (relative to the 35% seen in non-recovered sequences and to the 4% seen in the designs without significant change in proliferation (**Fig. 5G**)).

The sequences that exhibited CD20-specific depletion contained a large number of cysteines that were predicted to form disulfides within the binder chain (**Figs. 5H, SA11**). Given that tonic signaling can be enhanced through disulfide bond formation between hinge domain sulfhydryls^52–54^, and that inefficient folding (or, more likely, misfolding) may expose the hinge sulfhydryls or those in the binding domain, weak tonic signals might be greatly enhanced upon CD20 contact and subsequent disulfide bond formation.

Beyond these individual correlations, global confidence statistics from structure prediction models, such as ipTM and pLDDT, did not predict experimental outcomes (**Table SA1**). Likewise, ipSAE^55^, LIS score^56^, SAP score^57^, Rosetta InterfaceAnalyzer metrics^58^, PDockQ^59,60^, ProteinMPNN likelihoods, and ESM-2 pseudo log likelihood (PLL) also lacked substantial predictive power (**Table SA1**). For the latter two metrics, artifacts in the data led to deviation from expected scoring behavior. Likelihoods obtained from ProteinMPNN and SolubleMPNN on the Boltz-1-predicted structures were generally biased in favor of designs created using these models (**Fig. SA15**). ESM-2 PLL correlated with, and slightly underperformed in predictive power (**Table SA1**), amino acid sequence entropy for most sequences (**Fig. SA16**). This trend can be explained by the masked language modeling task becoming easier the less entropic the sequence is, which is further exemplified by the two outlier clusters defying this trend which contained large repeated regions that plausibly make the task of single residue infilling trivially easy with an in-context lookup, in agreement with existing literature^61^.

Overall, these findings exemplify how simple metrics can be more predictive of function than complex deep learning-based scores. Filtering out designs with ≥60% GC content and <1.9 DNA sequence entropy, ≥45 AA repeats in the DNA, ≥8 glutamate repeats, and ≥30% of residues in a K+E alpha helix results removes 4,600 sequences and increases the percent recovered after proliferation from 56.8% to 81.2% while increasing the percent of those that had CD20-specific proliferation from 5.9% to 7.6%. Further removing any sequences with cysteines raises the CD20-specific proliferation rate to 10.6%, though it reduces the percent recovered slightly to 72.6%. Notably, this filter removes two of the designs from the top 10 that lacked function entirely (1304, 1892) while preserving the rest. In contrast, filtering out sequences with <0.5 Boltz-1 ipTM would have removed two of the top 10 designs that were broadly functional and had binding affinity (designs 1506 and 5981) while only increasing the recovery rate by 0.7% and the CD20-specific proliferation rate by 0.3%. The predictive features discovered herein may help constrain future CAR-T engineering, both by AI-driven methods and otherwise.

## Discussion

The *Bits to Binders* competition was hosted to evaluate AI-driven protein design tools via a discovery-to-therapeutic pipeline. We collected 12,000 binder designs from 28 teams around the world that were created with a wide variety of approaches. Only 204 variants could not be cloned, in part due to repetitive sequences with high GC content, and the rest were tested in LEAH Labs’ high-throughput screen to evaluate a variety of phenotypes, starting with CD20-specific CAR-T proliferation. 4,985 of the designs (42%) dropped out of the population, indicating that some sequences or proteins interfered with cell viability. To verify that CD20-specific proliferation in the pooled growth assay translated to overall T cell function, we performed a series of functional assays on the ten best-performing unique designs, measuring specific cytotoxicity, cytokine release, proliferation, and expansion. We found that the majority of binders (7/10) outperformed the negative control across CD20-specific cytokine release, proliferation, and expansion, and 4/10 outperformed it on CD20-specific cytotoxicity. Two of the otherwise unsuccessful designs exhibited high non-specific killing. Ultimately, all of the designs underperformed the scFv positive control in all functional readouts except for cytokine production, in which design 5300 achieved more than 50% higher intracellular accumulation when cultured with CD20+ targets. As a caveat, while we established that some of the designs have broad T cell function, this was only performed in a single clinical isolate and the efficacy of these CARs in different donor lines is currently unknown.

Almost all of the designs that failed to be detected in the proliferation screen (98.9%) were created with ProteinMPNN or SolubleMPNN. We found that most of the failed MPNN designs preferentially contained highly adenosine-rich, lysine- and glutamate-containing alpha-helices, encompassing an average of 35% of residues versus only 15% in the recovered designs. While unnatural structures may pose problems on their own, we believe that many of the MPNN designs failed because of translational interference by glutamate and/or lysine^47,48,49^. To the extent that this hypothesis is correct, it provides a cautionary note that beyond the structure-centric task of binding, optimizing a protein’s amino acid sequence, and corresponding nucleotide sequence, is important for success, especially designing specifically for cellular expression. That said, there was clearly a sweet spot for design, as lower abundance K+E alpha-helices (18%) were moderately predictive of CD20-specific proliferation, in contrast to only 4% K+E in designs that did not show significant changes in proliferation. Designs associated with CD20-specific depletion, potentially indicative of exhaustion^62^, were highly enriched in cysteines. This validates the cautionary choice many model developers have made in explicitly avoiding cysteines in generated proteins.

Analyzing the top teams, we found some correlations between methodological choices and experimental outcome. BAGEL, Chroma, Bindcraft, and RFdiffusion all performed moderately well at CAR binder design. However, most of the sequences in this competition were created using RFdiffusion and ProteinMPNN, and this monoculture eventually skewed failure modes (K+E failures). Annealing structures with molecular dynamics prior to or after generation seemed to increase design success, whereas FastRelax and AmberRelax had no effect. While the top-performing teams employed iterative diffusion, this did not increase success across the board.

Overall, the *Bits to Binders* competition demonstrates that generative protein design has advanced to the point that it can directly be assessed by functional benchmarking. In particular, the identification of potential failure modes (GC dropouts, K+E abundance, excessive cysteines) can further guide CAR-T design, with a set of relatively simple filters almost doubling the rate of CD20-specific proliferating designs. By designing for a functional outcome, the results also begin to piece apart the complex interplay between expression, display, background (tonic) signaling, ligand-dependence, and proliferation. In particular, these results provide a counterpoint to the common perception that robust scFVs are the only viable starting points for CAR-T design, and raise the interesting prospect of initially canting design towards lower affinity spaces. The datasets and principles emerging from efforts like *Bits to Binders* should continue to serve as milestones for standardizing evaluation practices, guiding the development of more biologically informed models, and accelerating the application of AI-driven protein design for functional and ultimately clinical applications.

## Online Methods

### Selecting designs to test in each assay

Teams were asked to submit 500 sequences in order of confidence of success. In total, 28 teams submitted designs. The first 466 sequences from each team were selected to accommodate 12,000 total designs, including 16 designs chosen by LEAH Labs. Some teams submitted less than 466 designs or submitted duplicates, resulting in fewer total sequences: Foldsmiths: 465, Furman lab: 433, Nucleate UK London: 414, Perforators: 343, Zist Rayanesh: 308, Amigo Acids: 187, and Virtue Therapeutics: 48. 32 of the submitted sequences contained “X” as one of the amino acids which was arbitrarily converted to “L” by Twist Bioscience.

The designs were sorted by -log10(FDR)*logFC, and the top ten non-redundant designs were chosen to proceed to the individual T cell functional assays. These designs came from the following teams: CAD Pro (1304), Perez Lab Gators (1506), LBM (1892), Nucleate UK London (2383), AIBI (3109), SNU LCDD (3494, 3718), Amigo Acids (5300), Binding Illini (5624), Schoeder Lab (5981). Sequence redundancy was determined by sharing at least 50% sequence identity (exact match).

For the binding affinity study, designs were pulled from the three CD20-specific proliferation categories: ≥2-fold increase, ≥2-fold decrease, and no significant change. Sequences from each category were clustered with MMseqs2. For the ≥2-fold decrease category, 10 sequences were chosen in order of ascending -log10(FDR)*logFC, skipping if a sequence from the same cluster was already chosen. For the no significant change category, a random sequence from 60 random clusters was chosen. For the ≥2-fold increase category, the 10 sequences that underwent individual functional assays were chosen, in addition to 20 sequences randomly chosen from the ≥2-fold increase category, ensuring that each sequence was chosen from a distinct cluster. In total, this amounted to 100 sequences along with the control scFv.

### Plasmid library construction

The CAR construct began with a CD8α leader sequence, “stuffer” sequence (deoxyribose-phosphate aldolase) in the place of a binder via HiFi Assembly (New England Biolabs), a CD28 hinge and transmembrane domain, a CD28 co-stimulatory domain, and a CD3ζ signaling domain (modified from^63^). The F31S dihydrofolate reductase (DHFR) mutant was included after a P2A cleavage site, 3’ to the CAR construct, for further enrichment of the edited T cell population via selection with methotrexate (MTX)^64^. The stuffer sequence was removed while the Whitlow 218 linker was introduced via HiFi Assembly (New England Biolabs). This “binderless” construct was onboarded for oligo pool cloning of the 12,000 designed 80-amino-acid binders between the CD8α leader sequence and the 218 linker by Twist Bioscience.

### Pooled CAR-T generation and proliferation screen

CAR-T cell libraries were generated with GeneWeld^38,39^, which enables precision integration of the CAR construct through homology mediated end joining by targeting the genome with CRISPR/Cas9 and the donor plasmid with a Universal gRNA to liberate the cargo and reveal 48 nucleotide homology arms. CAR variants were integrated near the N-terminus of the T cell receptor alpha chain, resulting in TRAC-negative, CAR-positive T cells.

Primary human T cells, previously isolated by BACS selection (Akadeum Life Science) from healthy donor leukopaks (Sanguine Biosciences) and batch cryopreserved in CryoStor CS10 (STEMCELL Technologies). For experiments, T cells were thawed and allowed to recover overnight in T cell media comprised of X-Vivo15 + 5% AB serum and, after a brief stimulation with DynaBeads at 3:1 bead to cell ratio, GeneWeld components were delivered via Nucleofection in P3 Buffer using the EO-115 program (Lonza). After a 72 hour recovery in T cell media containing 10 ng/mL each of IL-7 and IL-15, and CAR-T cells were transferred to GREX wells for an additional 7 days of expansion under methotrexate selection. After sampling gDNA for NGS of the “Input” sample, the cells were subject to repeated challenge with mitotically-inactivated (MI) - Raji-GFP target cells at an effector to target ratio of 1:1 in triplicate. Briefly, Raji-GFP cells were mitotically-inactivated with 10 ug/mL Mitomycin C (STEMCELL Technologies) with for 2 hours at a cell concentration of 1E6 cells/mL before washing three times and cryopreserving at 10-20E6 cells/mL in CryoStorCS10 for storage at −80°C. Running in parallel with MI-Raji-GFP co-cultures, the same CAR-T cell libraries were cultured alone in a GREX 6-well plate in triplicate to control for false-positives. After 2 weeks, a minimum of 12E6 cells were harvested per replicate for genomic DNA isolation, using Quick-DNA Miniprep Plus Kit from Zymo Research, and NGS analysis in triplicate for both conditions^65^. PCR was done with Herculase II Fusion DNA Polymerase from Agilent to add Nextera adaptors. PCR purification was done using the SPRIselect DNA Size Selection Reagent from Beckman Coulter. Samples were submitted to the University of Minnesota Genomics Center for further library preparation and sequencing using the Element AVITI24 sequencing platform.

Variants were counted, and those represented by fewer than 25 reads in any replicate were excluded from further analysis. For each construct, CD20-specific proliferation was defined as the ratio of normalized barcode abundance in the CD20-positive condition relative to the CD20-negative condition, CAR-T cells alone as a control for false-positive, non-specific expansion. Statistical significance was assessed using edgeR^66^, and false discovery rates (FDRs) were controlled at 5%.

### CAR-T cell immunophenotyping by flow cytometry

Following methotrexate selection, CAR-T cells were characterized by multi-parameter flow cytometry. 1E5 cells were harvested into a round-bottom 96-well plate and washed with ice-cold MACS Buffer (PBS supplemented with 0.5% BSA and 2 mM EDTA). Surface antigen staining was performed by incubating cells with a fluorophore-conjugated antibody cocktail targeting the CAR (anti-218L-AlexaFluor 488, Cell Signaling Technologies) and the TCR (anti-TCRa/b-APC, Miltenyi Biotec), each at a 1:50 dilution in MACS buffer for 10 minutes at 4°C in the dark. To assess viability and exclude non-viable cells from analysis, cells were washed with MACS buffer and stained with eBioscience Fixable Viability Dye eFluor 450 (Invitrogen) at a 1:500 dilution for 10 minutes at 4°C. After two final washes, cells were resuspended in MACS buffer and acquired on a MACSQuant Analyzer 16 flow cytometer (Miltenyi Biotec). Data were analyzed using FlowJo software (v10) with the following hierarchical gating strategy: 1) Identification of lymphocytes based on forward scatter (FSC-A) and side scatter (SSC-A) to exclude debris, followed by 2) doublet discrimination via FSC-A vs FSC-H to ensure single-cell analysis, and 3) gating on the viability dye-negative population to exclude dead cells, allowing 4) CAR and TCR subset and expression analysis within the live cell population. Controls included Mock-transfected T cells (TCR+CAR-) and unstained (FMO) samples that did not receive anti-CAR (218L) or anti-TCR antibodies.

### Specific lysis individual T cell functional assay

CD20+ target cells-specific cytotoxicity was evaluated after co-incubating 25,000 CAR-T cells per well with CD20-positive target cells (Raji-GFP) and CD20-negative controls (K562-GFP) at varying effector-to-target ratios for 48 hours. A MACSQuant Analyzer 16 flow cytometer (Miltenyi Biotec) was used to quantify specific lysis by counting the number of GFP+ target cells remaining per well when co-cultured with T cells, relative to wells containing targets cells alone as no lysis controls. Specific Lysis = (number of GFP+ targets remaining / target cells alone) x 100%. Multiple effector-to-target ratios allowed for AUC analysis to compare specific lysis of CD20+ targets (Raji^AUC^-K562^AUC^) between individual CAR-T clones of interest and control T cells.

### Cytokine generation individual T cell functional assay

T cells were cultured alone or co-cultured with Raji-GFP target cells at a 1:1 E:T ratio for 16 hours before adding Brefeldin A (BioTechne) for an additional 4-5 hours. After activation, cells were harvested, washed and stained with eBioscience Fixable Viability Dye eFluor 450 (Invitrogen) at a 1:500 dilution for 10 minutes at 4°C to exclude non-viable cells from analysis before fixation, permeabilization, and intracellular cytokine staining using CytoFix/CytoPerm kit (BioTechne). Specifically, anti-IL2-PE and anti-IFNg-VioR667 REA antibodies, along with fluorochrome conjugated REA isotype controls (1:50 dilution, Miltenyi Biotec), were utilized to enumerate cytokine producing T cells with a MACSQuant Analyzer 16 flow cytometer (Miltenyi Biotec). Fluorochrome-conjugated REA Intracellular isotype antibodies were used as controls to assess the percentages of T cells producing either IL2, IFNγ, or both were added together to reflect total frequency of cytokine producing T cells.

### Expansion and proliferation individual T cell functional assays

CAR-T cell division and cumulative expansion were tracked over 5 days after stimulation with CD20-positive targets at varying effector-to-target ratios. Expansion was quantified as total viable CAR-T yield 5 days after stimulation using volumetric sampling for accurate cell enumeration with the MACSQuant Analyzer 16 flow cytometer. T cell proliferation over 5 days was simultaneously inferred from cell division rates measured by the change in the mean fluorescence intensity (MFI) of Cell Trace Violet (Thermo Fisher Scientific) in T cells labeled according to the manufacturer’s instructions. Multiple effector-to-target (E:T) ratios along with no MI-Raji target controls allowed for AUC analysis to compare expansion and proliferation between individual CAR-T clones of interest and control T cells.

### Cell-free expression of binder variants for binding affinity

DNA constructs encoding the binder variants were designed by reverse-translating the protein sequences and optimizing them for manufacturability and yield using codon optimization algorithms. Each construct was fused to a C-terminal GFP11-TwinStrep (TST) tag to enable split-GFP-based expression quantification and affinity capture for kinetic characterization. Optimized variant sequences and the 3′ fragment encoding linker and affinity tags were ordered as Gene Fragments (Twist Bioscience). Constructs were assembled using the NEBuilder HiFi DNA Assembly Kit (NEB) in 2 μL reactions. Assembly quality was verified by capillary electrophoresis (Agilent ZAG DNA Analyzer), and DNA concentrations were quantified using the Qubit DNA Quantification Kit (Invitrogen).

Protein expression was performed in 8 μL reactions using an optimized prokaryotic cell-free protein synthesis (CFPS) system with 4 nM assembled DNA template. Reactions were incubated at 37 °C for 8–12 hours. Soluble expression was confirmed using a split-GFP complementation assay, and protein yields were normalized using an affinity-based quantification assay. Crude lysates were used directly for downstream kinetic measurements.

### Surface Plasmon Resonance (SPR) Affinity Characterization

Binding kinetics were characterized using SPR on a Carterra LSA XT high-throughput platform. A carboxymethylated chip surface was functionalized with Strep-Tactin XT (IBA Lifesciences) using EDC/NHS chemistry. TwinStrep-tagged binder variants (from crude CFPS lysates diluted 125-fold) were captured onto the surface using the 96-channel array head (750 s capture, 600 s baseline).

The target antigen, full-length human CD20 (Acro Biosystems, CD0-H52D5; Flag/His-tagged, detergent-solubilized), was reconstituted in running buffer (50 mM HEPES, 150 mM NaCl, DDM + CHS, pH 7.5) and injected in solution over the immobilized variants. CD20 was tested at five half-log serial dilutions starting from 1000 nM. Measurements were performed in triplicate using multi-cycle kinetics, consisting of 60 s baseline, 300 s association, and 600 s dissociation. Surfaces were regenerated with 10 mM glycine-HCl (pH 1.5) between full concentration series.

### Bio-Layer Interferometry (BLI)

A subset of variants was additionally tested by BLI on a Gator Pro system using the same CD20 preparation and detergent-containing buffer (50 mM HEPES, 100 mM NaCl, DDM + CHS). Binder variants were immobilized, and CD20 was run in solution as analyte. Assay steps consisted of 60 s baseline, 120 s loading, 120 s baseline, 60 s association, and 300 s dissociation.

### Data Processing and Kinetic Analysis

Sensorgrams were reference- and baseline-corrected and globally fitted to a 1:1 Langmuir binding model using Adaptyv Fitting software when signal quality permitted. Kinetic parameters (*KD, kon, koff*) were extracted from global fits across concentrations. When global fitting was not feasible, alternative models were applied and parameters were selected based on fit quality.

Variants were classified as binders based on quantifiable kinetic fits. In cases where fitting was not reliable but association-phase signal exceeded 300% of the negative control, binding was assigned based on signal magnitude and estimated using Anabel^43^.

### Computational feature generation and statistical modeling

For each submitted binder, we computed a panel of >400 features from the nucleotide and amino acid sequences (**Supplementary Information B**). Sequence-derived metrics included nucleotide and amino acid composition (for example, GC content), k-mer and repeat statistics (including homopolymers and dipeptide repeats), and Shannon entropy at the DNA and protein levels to quantify sequence complexity. Structure-informed features were derived from Boltz-1, ESMFold, and Chai-1 predictions of binder–CD20 complexes. These included model confidence statistics (e.g., pLDDT and ipTM), global and per-residue structural descriptors (e.g., secondary-structure composition and predicted disulfide content), and interface-focused metrics computed on the predicted complexes (e.g., contact counts and interface energy or interaction scores).

Features were analyzed against four outcomes: detection in the Twist plasmid pool quality report, recovery by NGS after the pooled proliferation assay, and CD20-specific change in proliferation. For binary outcomes, we trained logistic regression models using stratified 5-fold cross validation and evaluated performance with ROC-AUC. To mitigate method-specific confounding, analyses were repeated on restricted subsets when appropriate (e.g., only ProteinMPNN/SolubleMPNN-generated designs). Feature relevance was assessed by comparing cross-validated performance across individual features and by inspecting class-conditional feature distributions.

## Data Availability

All data associated with this study is publicly available on Zenodo at doi.org/10.5281/zenodo.18840101. Summarized data used for the analysis can also be found at github.com/kosonocky/bits-to-binders.

## Code Availability

Code used in the competition analysis is publicly available on Zenodo at doi.org/10.5281/zenodo.18840101 and at github.com/kosonocky/bits-to-binders. Code used by the competitors is listed in Supplementary Information E, when available. Inquiries on the competitors’ code should be directed to the contacts listed therein.

## Author Contributions

Conceptualization: C.W.K., A.M.A., A.L.F., D.R.B., S.C.E., A.D.E., W.A.W., E.M.M.; Organization and running of competition: C.W.K., A.L.F., P.R.W., D.R.B., S.C.E., W.A.W.; Design of *de novo* binders: *Bits to Binders* competitors; CAR-T experimental assays and data analysis: A.M.A., A.E.C.R., W.A.W.; Computational analysis of methods and designs: C.W.K., P.R.W., J.L., T.G.; Supervision: S.C.E., A.D.E., W.A.W., E.M.M.; Funding and sponsorship acquisition: C.W.K., A.L.F., D.R.B., S.C.E., A.D.E., W.A.W., E.M.M.; Writing — original draft and figures: C.W.K., A.M.A., A.E.C.R.; Writing — review and editing: All authors.

## Acknowledgements

We thank Claus Wilke for feedback on the manuscript and figure structures, Alia Clark-Elsayed for feedback on integrating competitor information into the manuscript, Daniel Gutierrez and Julian Englert for feedback on interpreting binding affinity data, Adam Salmi for creating the trophies, and Ron Green and Nathan Mandi for helping organize machine learning office hours for the competitors during the kickoff event.

The *Bits to Binders* competition was made possible by the generous support of academic and industrial sponsors. We thank LEAH Laboratories, Adaptyv Bio, Twist Bioscience, the University of Texas at Austin Center for Systems and Synthetic Biology, the Texas Advanced Computing Center (TACC), Modal, Lonza, ScaleReady, VWR, KUNGFU.AI, Maker Clinic, Synthia, the University of Texas at Austin Dell Medical School, Nucleate AI in Biotech, and the BioML Society.

We acknowledge additional funding from the Winkler Family Foundation (A.D.E.); the National Institutes of Health (R35GM122480 to E.M.M.); the Welch Foundation (F-1515 to E.M.M.; F-1654 to A.D.E. funding C.W.K.); BioMADE (T-OC-A-04-0087 to A.D.E. funding C.W.K.); DTRA-AI (HDTRA1201001 to A.D.E. funding C.W.K. and A.L.F.).

C.W.K. acknowledges funding from the University of Texas at Austin Graduate Continuing Research Fellowship and the Charles Smith, Jr. Graduate Fellowship in Molecular Biosciences. J.L. acknowledges the President’s PhD scholarship at Imperial College London for funding. D.R.B. was funded in-part by the Stengl-Wyer Endowment to The University of Texas at Austin’s College of Natural Sciences. The views and conclusions contained in this document are those of the authors and should not be interpreted as representing the official policies, either expressed or implied, of the U.S. Government nor other funding bodies. The U.S. Government is authorized to reproduce and distribute reprints for Government purposes notwithstanding any copyright notation herein.

Above all, we thank all participating teams and competitors whose designs were the foundation of this effort.

## List of *Bits to Binders* Competitors

Daniel Acosta, Stefano Angioletti-Uberti, Vinayak Annapure, Neil Anthony, Rana A. Barghout, Max Beining, Parth Bibekar, Dionessa Biton, Patrick Bryant, Anton Bushuiev, Cianna Calia, Ava Chan, Jie Chen, Akshay Chenna, Nicole Chiang, Vivian Chu, Joseph Clark, Alia Clark-ElSayed, Elliot Cole, Qian Cong, Shreyasi Das, Tanner Dean, Tyler Derr, Alejandro Diaz, Jesse Durham, Moritz Ertelt, Antonio Fonseca, Ora Furman, Tynan Gardner, Jokent Gaza, Nathaniel Greenwood, Rui Guo, Abel Gurung, Vivian Haas, Seyed Alireza Hashemi, Dieter Hoffmann, Sofiia Hoian, Corey Howe, Calvin XiaoYang Hu, Jinling Huang, Nader Ibrahim, Satoshi Ishida, Ansar Ahmad Javed, Aswini Javvadi, Roman Jerala, Daksh Joshi, Even Kiely, Haelyn Kim, Diego Kleiman, Sarah Knapp, Huan Koh, Petr Kouba, Yong Youn Kwon, Duško Lainšček, Jakub Lála, Hakyung Lee, Qiuzhen Li, Max Lingner, Yuxuan Liu, Yunchao Liu, Qianchen Liu, Ajasja Ljubetič, Marko Ludaic, Karl Lundquist, Ashish Makani, Javier Marchena-Hurtado, Freddie Martin, Jens Meiler, Fernando Meireles, David Miller, Nuria Mitjavila, Tjaša Mlakar, Rajarshi Mondal, Saeideh Moradvandi, Rocco Moretti, Sai Jaideep Reddy Mure, Pragati Naikare, Fatemeh Nasiri, Kaveh Nasrollahi, Yisel Martinez Noa, Joseph Openy, Aoi Otani, Vikrant Parmar, Jimin Pei, Alberto Perez, Rajat Punia, Francisco Requena, Dominic Rieger, Aparna Sahu, Elias Sanchez, Rohit Satija, Tadej Satler, Tom Schlegel, Yang Shen, Diwakar Shukla, Rangana De Silva, Benedikt Singer, Arjun Singh, Amardeep Singh, Bhumika Singh, Jinwoong Song, Harish Srinivasan, Hannah Stewart, Zhaoqian Su, Julia Varga, Xin Wang, Dion Whitehead, Matthew Williams, Max Witwer, Rujie Yin, Song Yin, Rongqing Yuan, Nikola Zadorozhny, Dishant Parag Zaveri, Genhui Zheng, Yizhen Zheng, Jian Zhou, Shaowen Zhu, Lin Zhu

## Supplemental Information

### A. Supplemental Figures and Tables

**Figure SA1.**
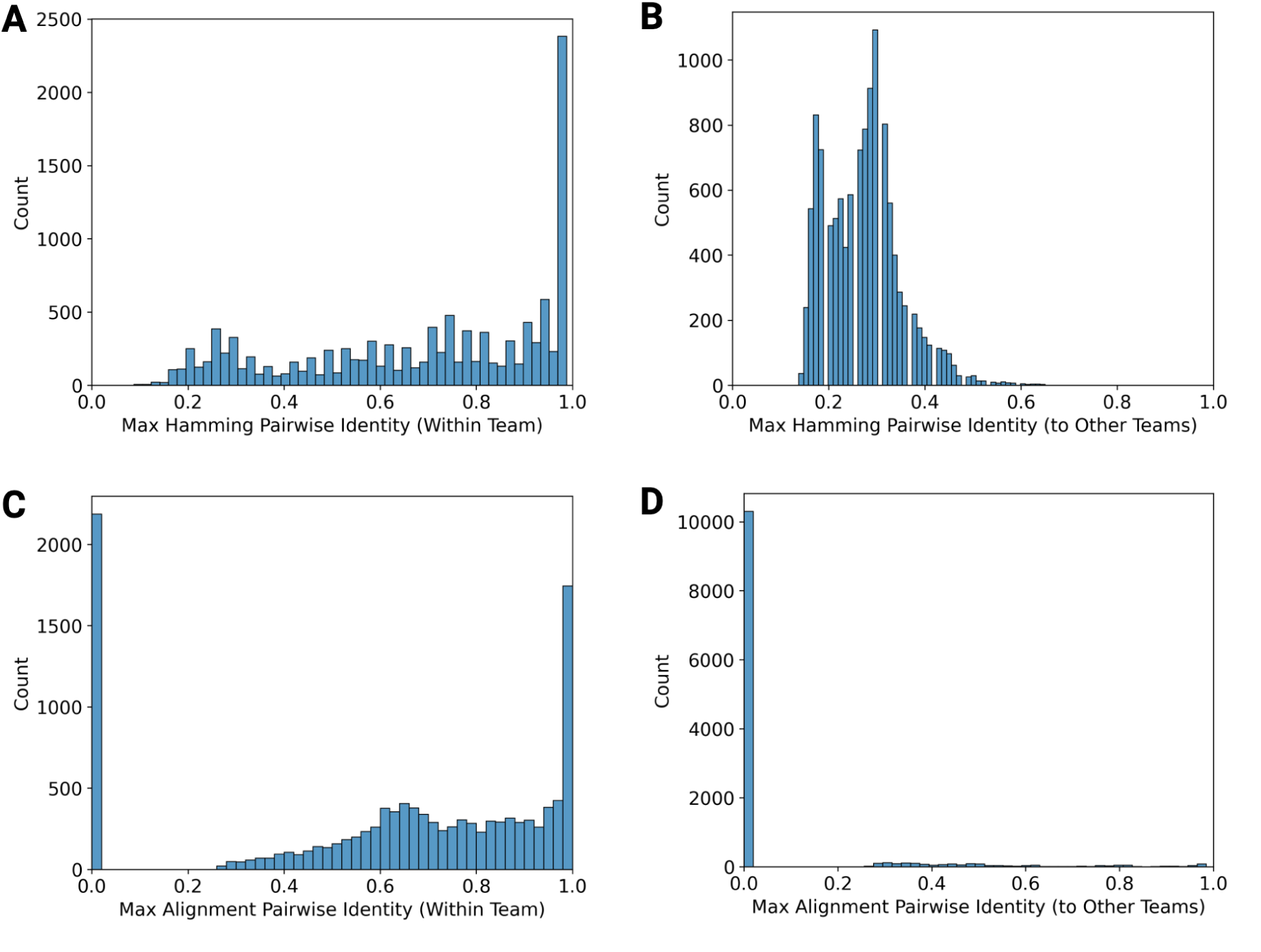
Maximum pair-wise sequence identities of the designed binders. The following sequence identity metrics were computed: **(A)** Hamming similarity (full sequence, position-by-position, ungapped) between each sequence and the other sequences from the same team, **(B)** Hamming similarity between each sequence and all sequences from the other teams, **(C)** Local alignment-based sequence identity (MMseqs2^37^, -s 7.5, --cov-mode 0, qcov & tcov ≥ 0.3) between each sequence and all sequences from the same team, **(D)** Local alignment-based sequence identity between each sequence and all sequences from the other teams. Insignificant alignments are set to zero pairwise identity. Although there was high within-team redundancy, the designs from a given team were largely unique relative to those from other teams. The exceptions to this were teams that included significant portions of known CD20 antibodies.

**Figure SA2.**
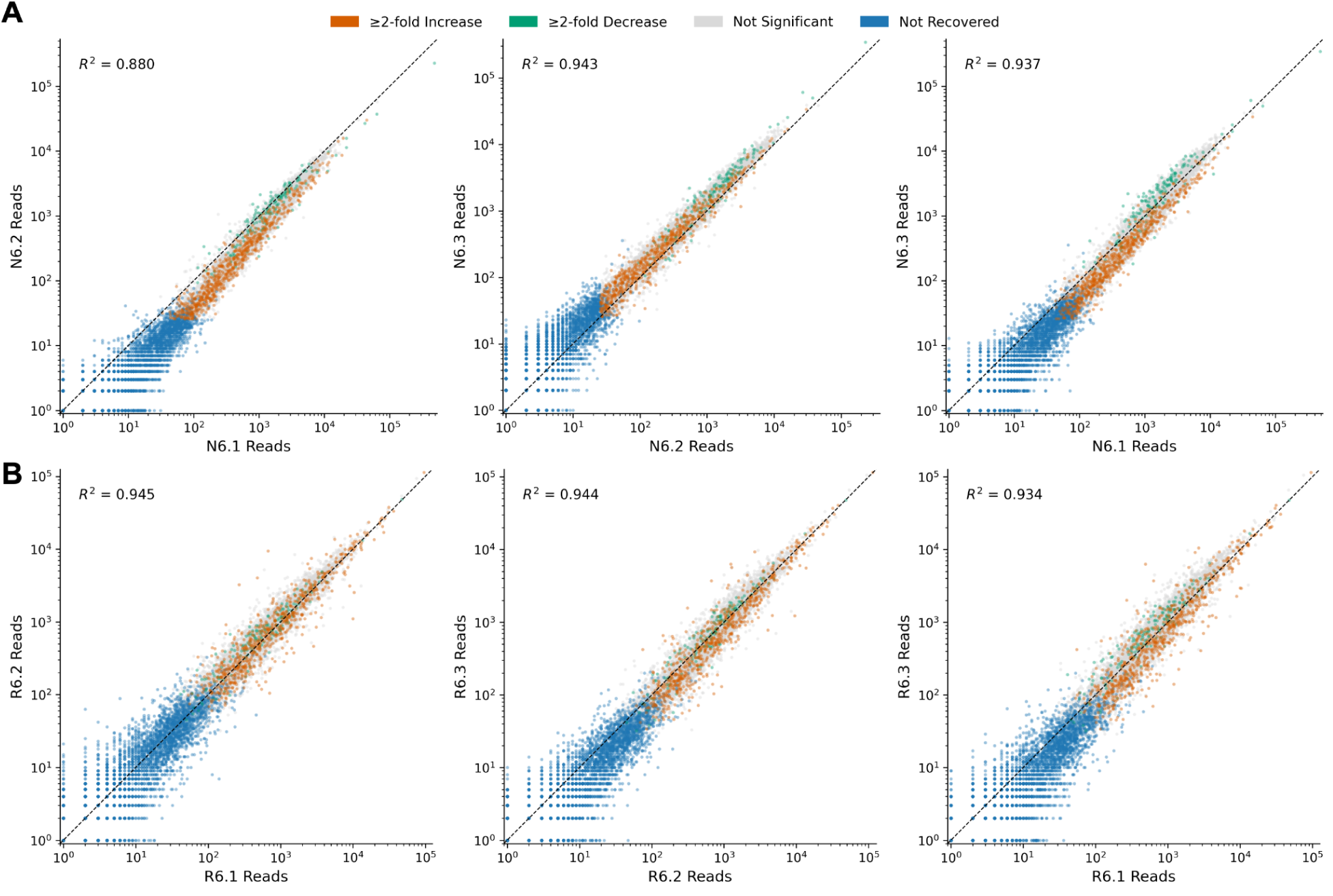
NGS reads demonstrate minimal jackpot effects between replicates in pooled proliferation assay. **(A)** Reads were generally correlated between the three non-CD20 replicates (N6) and the **(B)** three CD20+ co-culture replicates (R6), indicating that there was minimal jackpot effect occurring.

**Supplemental Figure SA3:**
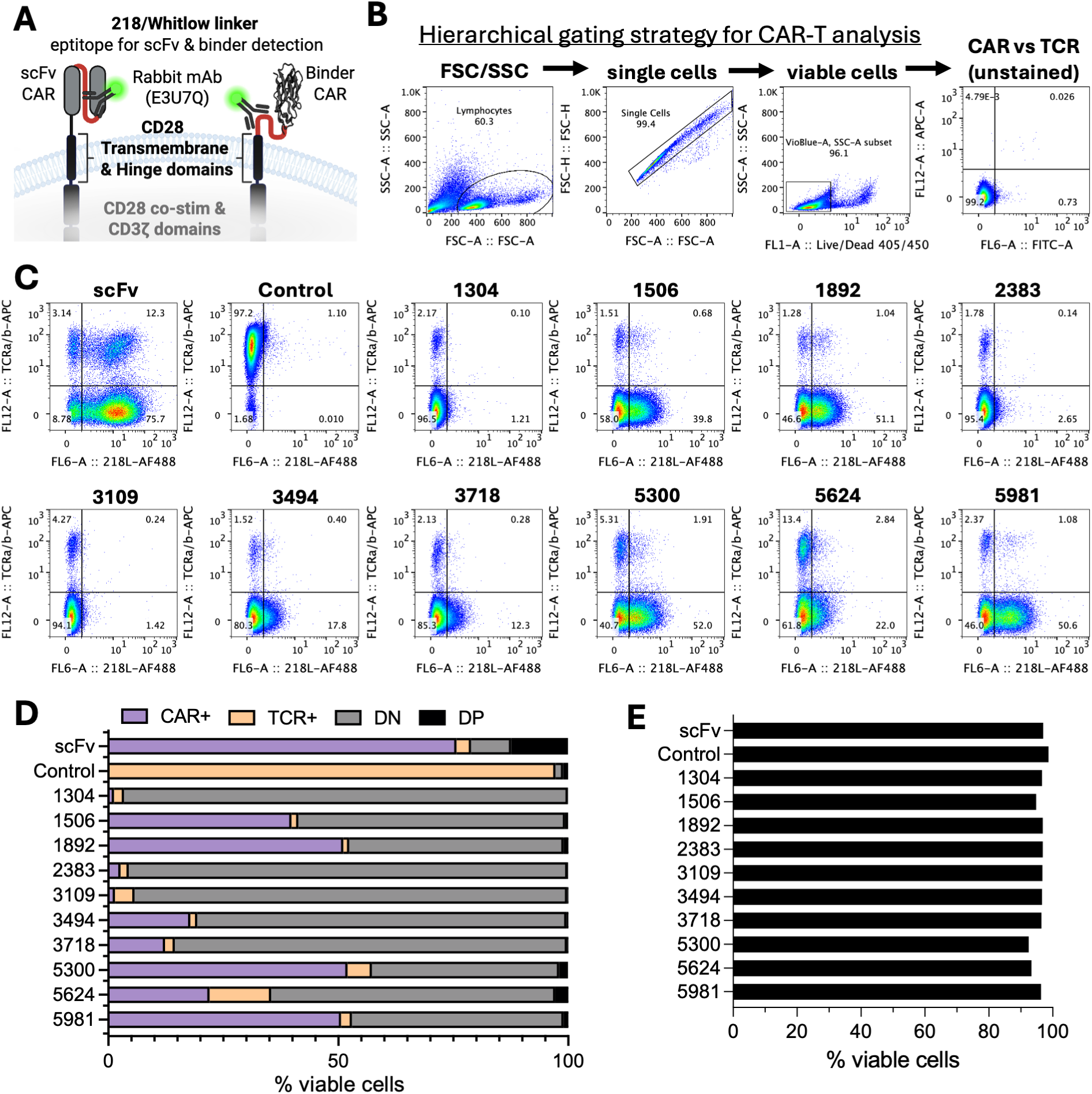
CAR and TCR expression determined by flow cytometry. **(A)** 218/Whitlow linker present in the scFv was designed into the binder construct as an antibody epitope. **(B)** Hierarchical gating strategy for viable CAR-T enumeration, as shown in **(C)** scatter plots depicting CAR and TCR surface expression for controls and binder variant CAR-T cells. **(D)** graphical representation of flow cytometry data showing T cell populations expressing CAR, TCR, neither, or both. **(E)** Viability assessment of controls and binder variant CAR-T cells.

**Figure SA4.**
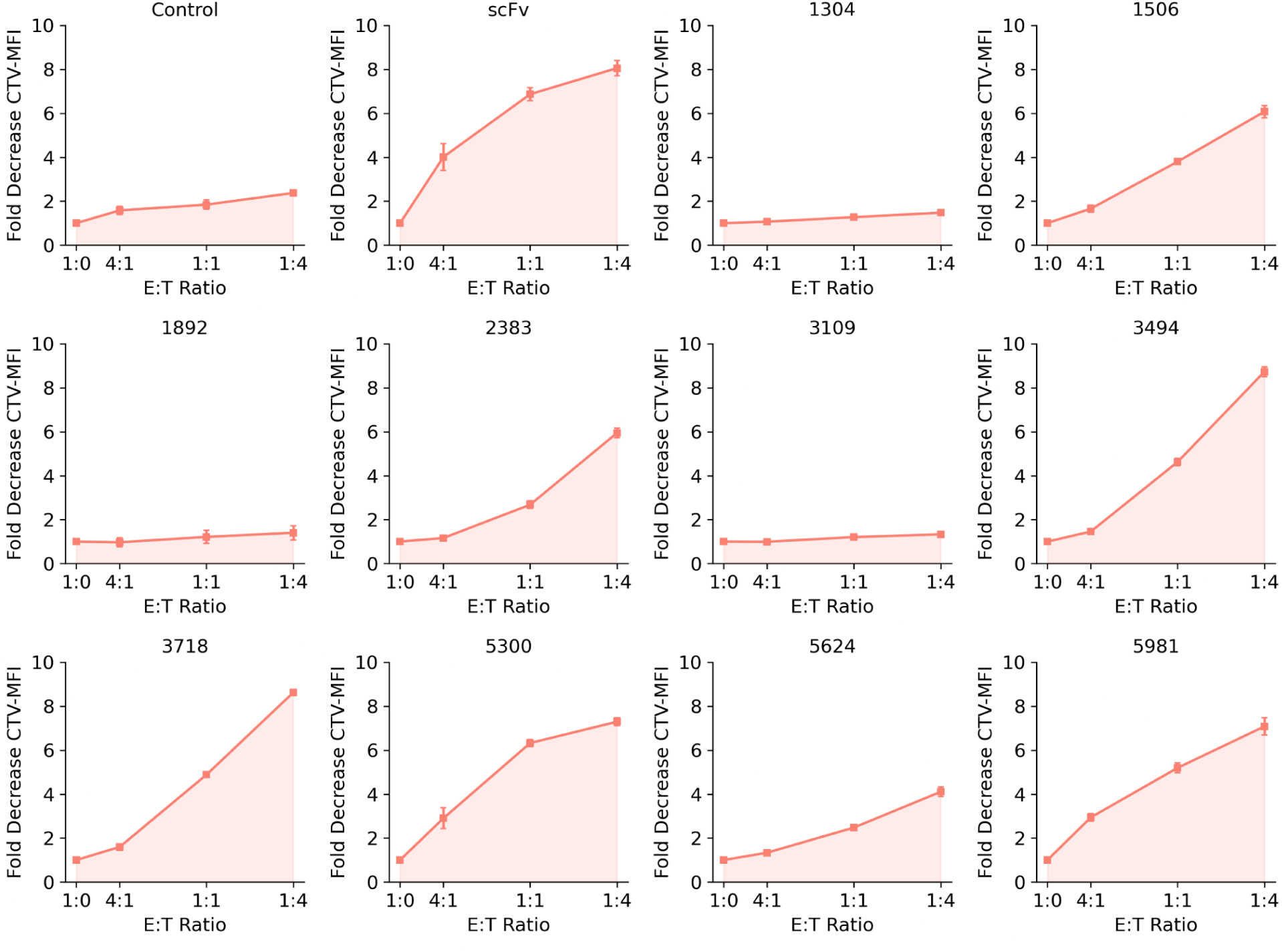
Proliferation via decrease in CTV-MFI. Proliferation was measured over 5 days following stimulation with Raji (CD20+) targets across multiple E:T ratios by decrease in median fluorescence intensity (MFI) of the Cell Trace Violet (CTV) signal.

**Figure SA5.**
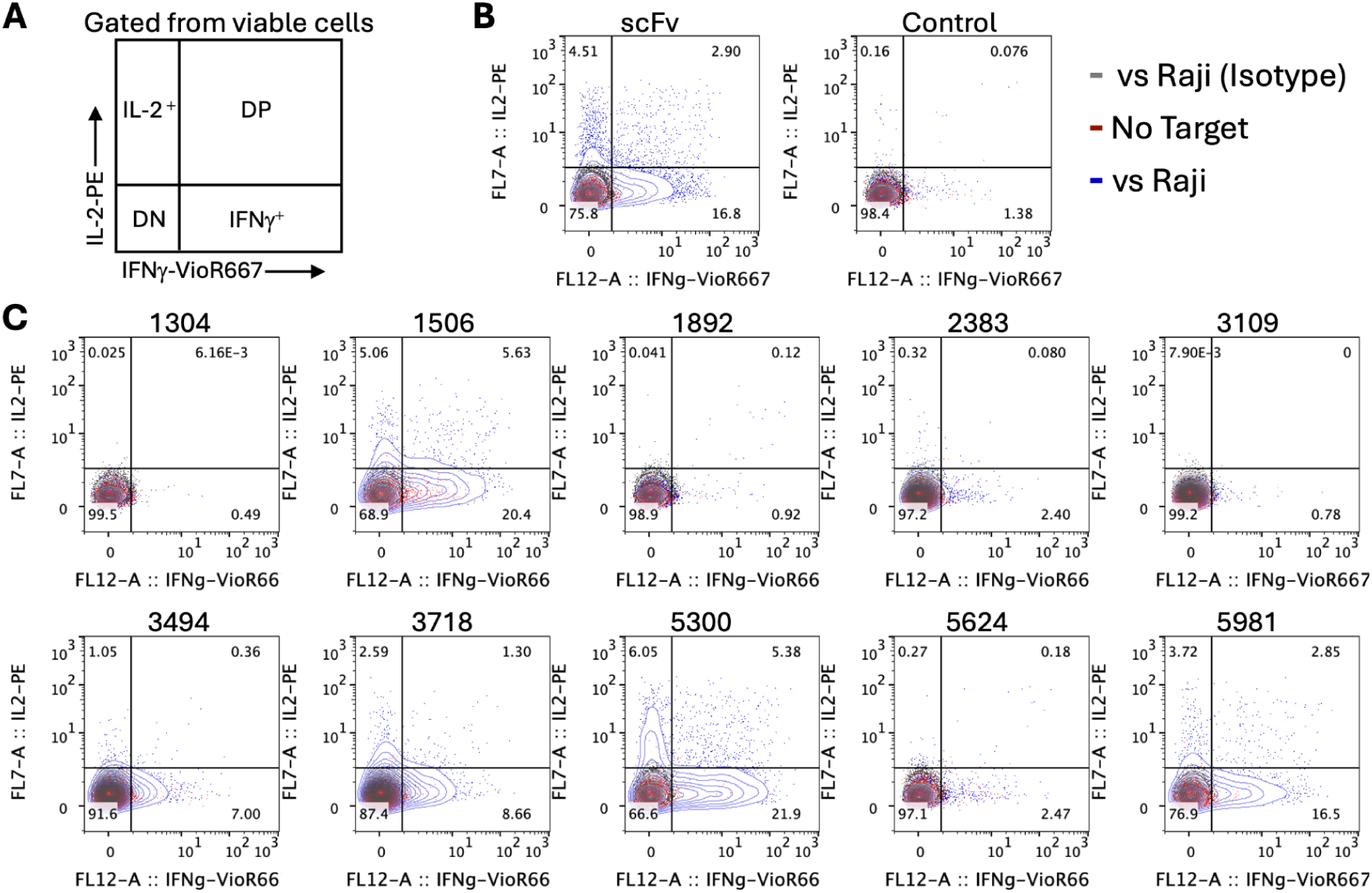
Flow cytometry data of T cell proliferation with CellTrace Violet (CTV) dilution. CTV (x-axis) peaks analyzed after 5 days in co-culture with MI-Raji target cells at various Effector-to-Target (E:T) ratios.

**Figure SA6.**
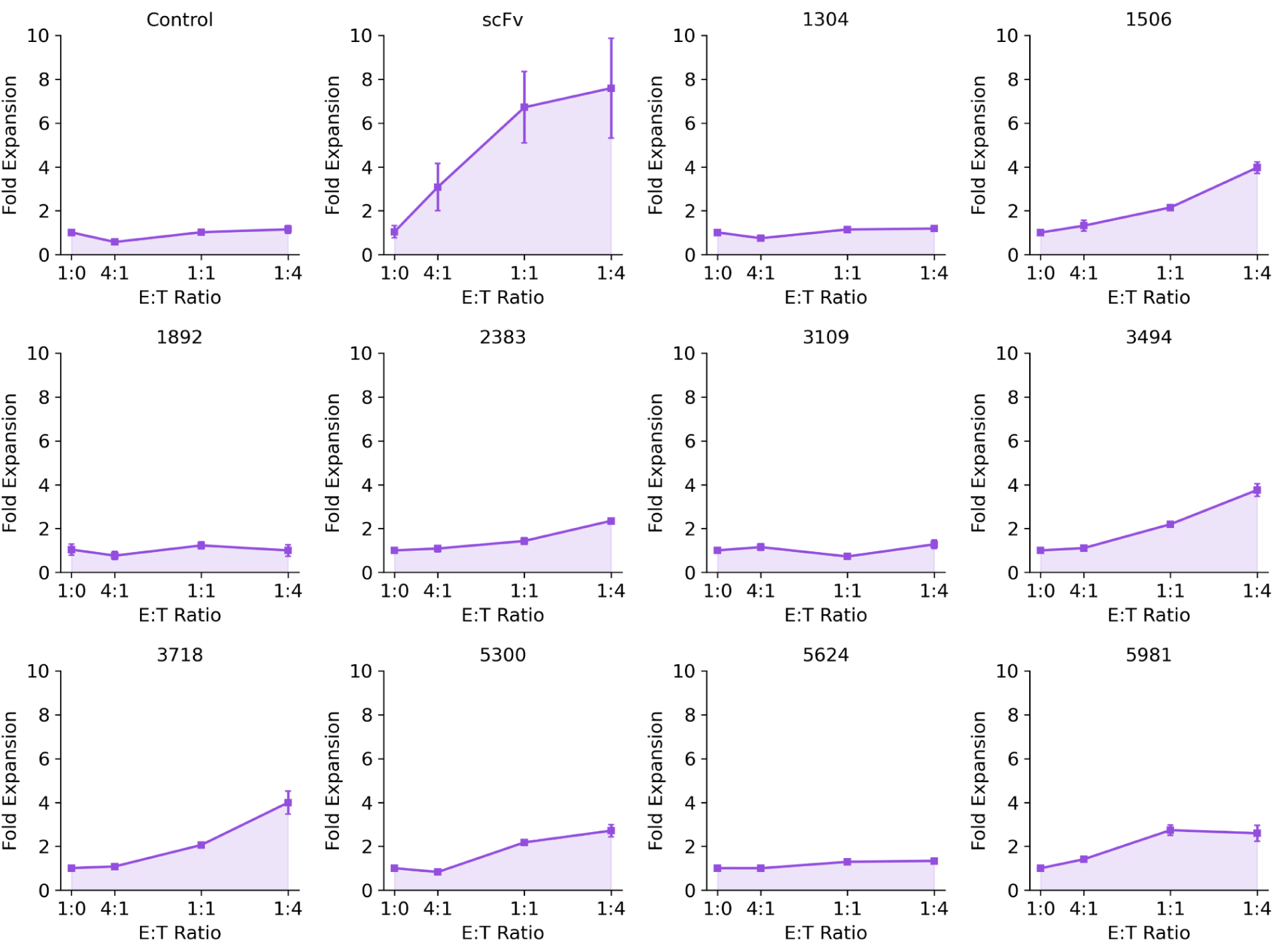
Fold expansion by cell counts. Total CAR-T expansion was measured over 5 days following stimulation with Raji (CD20+) targets across multiple E:T ratios by total viable CAR-T yield using volumetric flow cytometry.

**Fig SA7.**
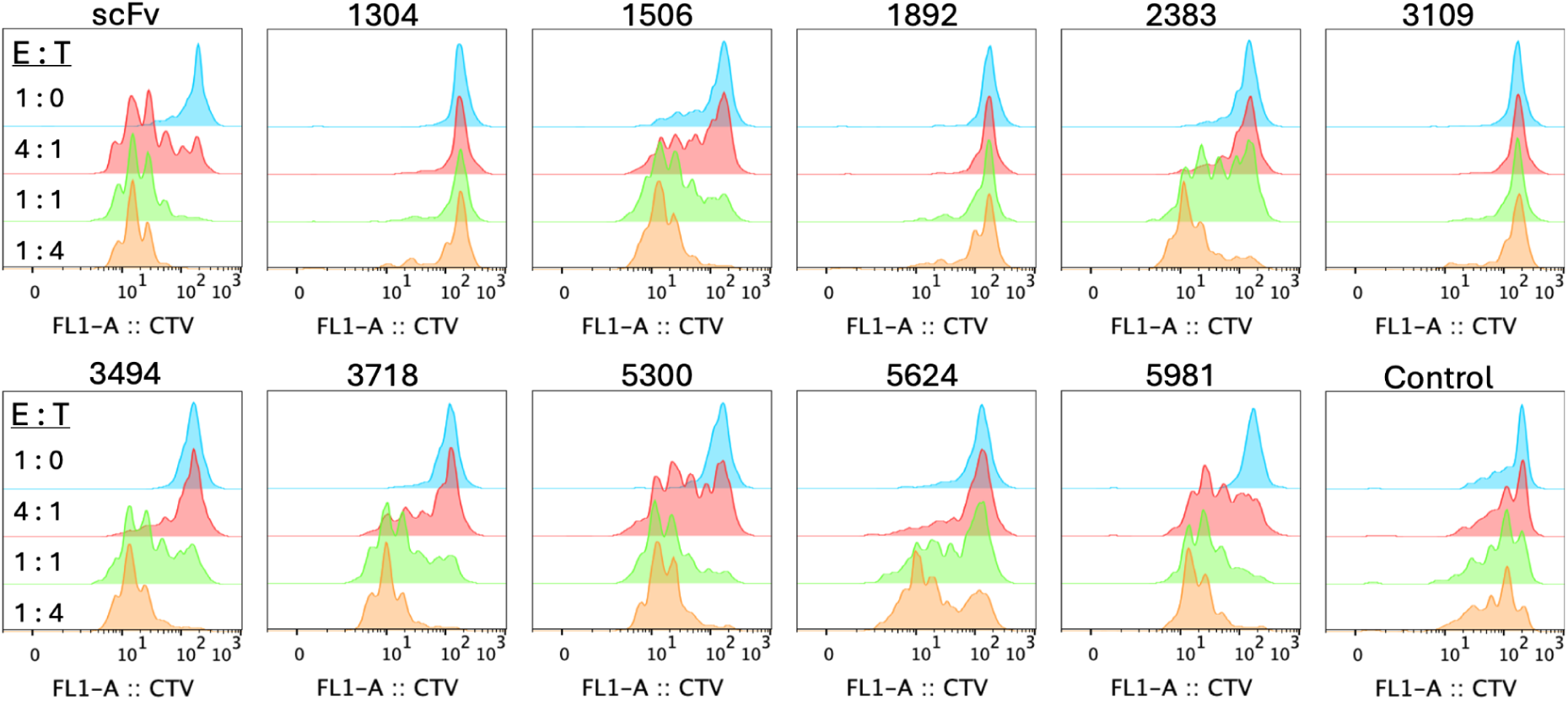
CAR-T cell cytokine generation induced by CD20+ Raji target cells. **(A)** Gating strategy illustrating cytokine-generating populations as analyzed by flow cytometry. **(B)** CD20 scFv CAR and Control (unedited, TCR+CAR-neg T cells), were stained with fluorochrome-conjugated antibodies after incubating alone (red) or in co-culture with Raji target cells. Specifically, anti-IFNg and anti-IL2 antibodies, or the requisite isotype antibody controls, were used to analyze both controls and **(C)** individual CD20 binders as CAR-T cells.

**Figure SA8.**
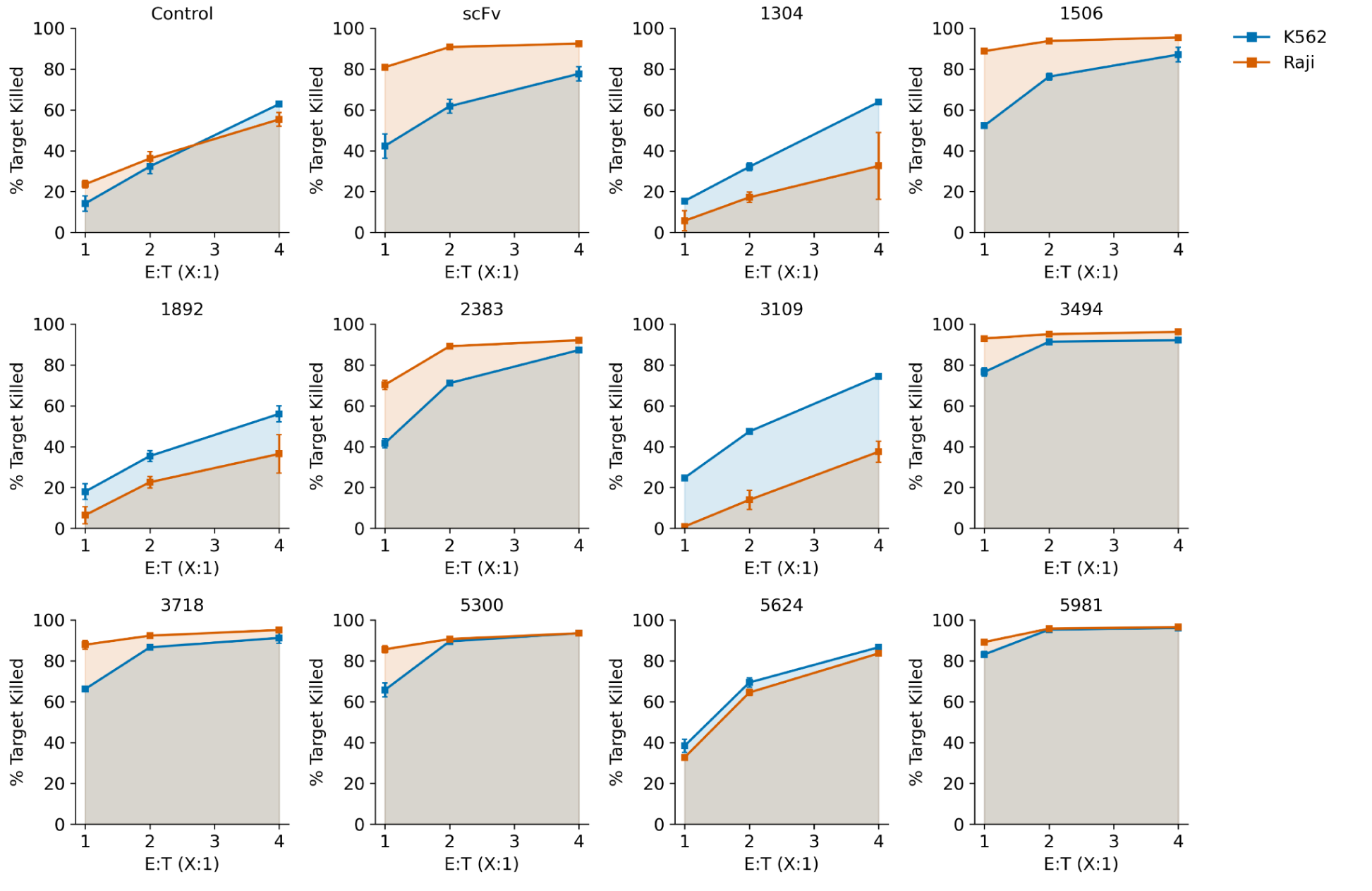
Raji-specific cytotoxicity vs K562 control. GFP assay readouts across multiple E:T ratios for Raji (CD20+) and K562 (CD20-) targets, overlaid for direct comparison.

**Figure SA9.**
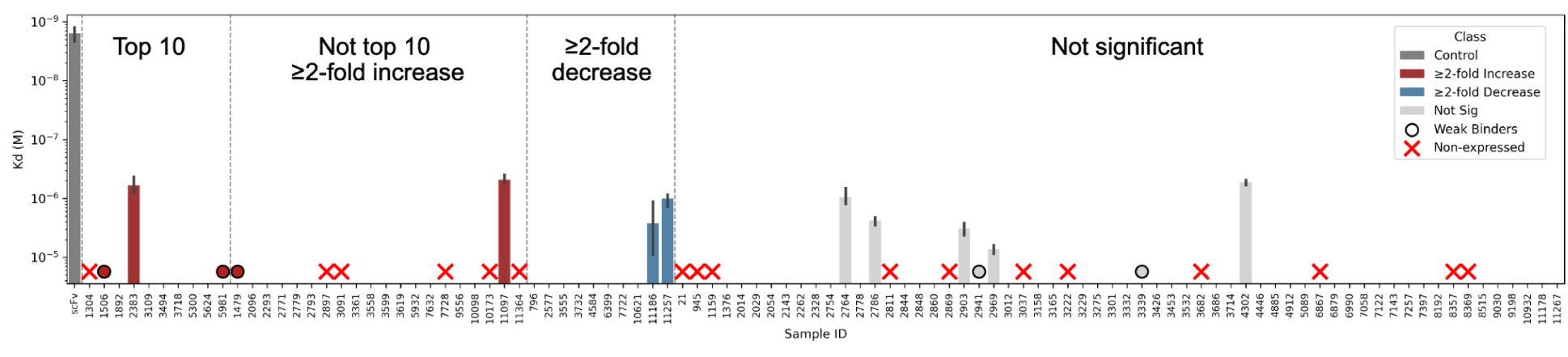
Binding affinities to detergent-solubilized CD20 for 100 binders expressed as 80-amino acid constructs by *in vitro* transcription/translation. Along with the positive control scFv, 100 binders were submitted for binding characterization via SPR and BLI spanning the three proliferation categories in triplicate. These include the top ten from the high-throughput proliferation screen, 20 additional designs showing CD20-specific ≥2-fold increase, 10 designs showing CD20-specific ≥2-fold decrease, and 60 that had no significant change. Those that demonstrated binding, but were not quantifiable in the experimentally tested concentrations, are denoted with a filled circle colored by its class. Those that were not able to be expressed as isolated 80-mer constructs by *in vitro* transcription/translation are marked with red X’s. Absence of a bar indicates successful expression but no observed binding.

**Figure SA10.**
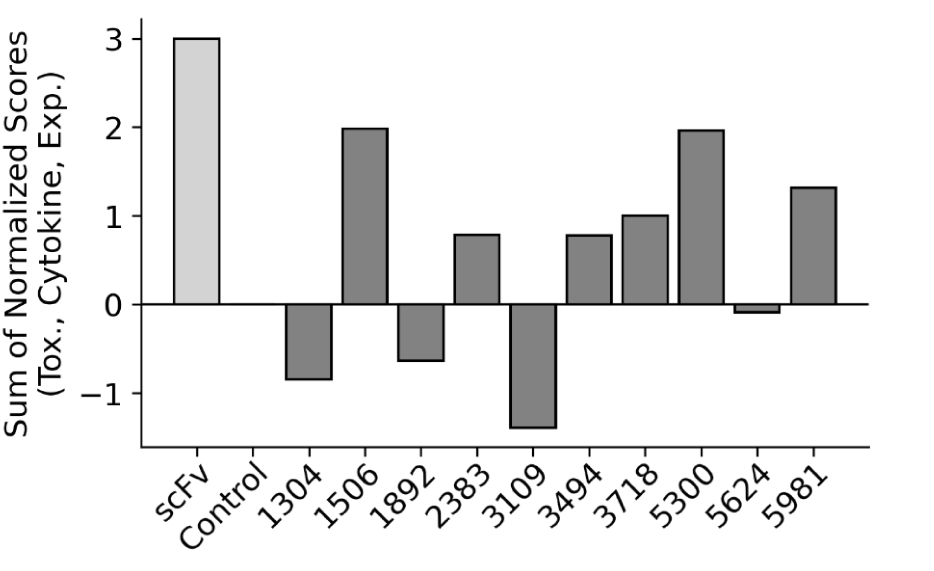
Rankings used to determine competition winners. To get an overall assessment on the functional performance of each selected design, the final rankings in the competition were determined by using cytotoxicity, cytokine, and expansion outcomes min/max normalized to the negative and positive control, respectively. These were chosen to reduce bias away from growth, which would have occurred had both proliferation and expansion been used in the final score.

**Figure SA11.**
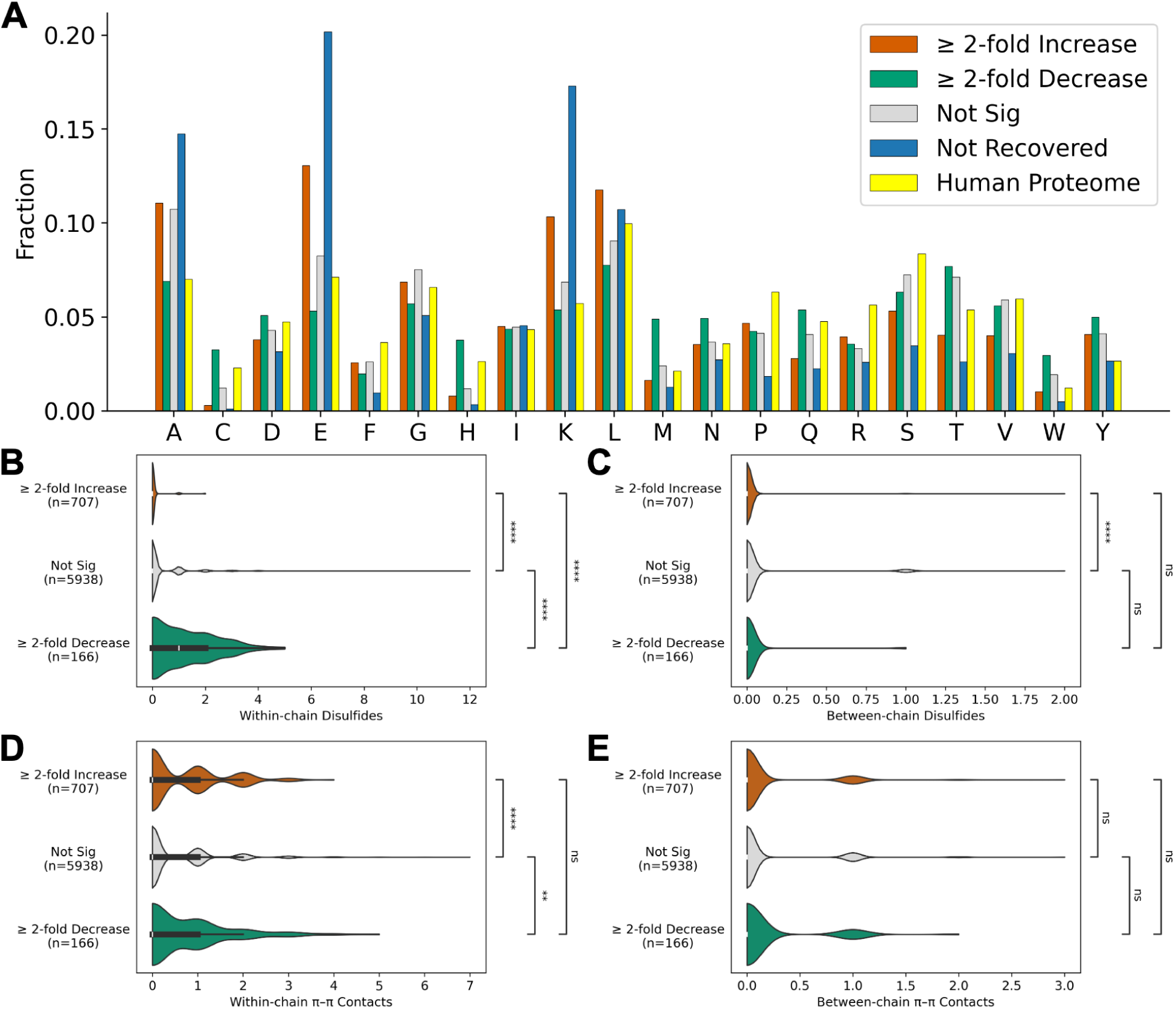
Amino acid compositional differences and predicted contacts. **(A)** Composition of the four categories obtained from the proliferation screen. Compared to the human proteome, the ≥2-fold increase category is enriched in A, E, K, and L and diminished in C and H, whereas the ≥2-fold decrease category was enriched in C, H, M, and W. **(B)** The ≥2-fold decrease category is enriched in predicted disulfide bonds within the binder chain on Boltz-1 predicted structures. **(C)** There is no trend for disulfides between the binder and CD20. **(D)** There is no trend for pi-pi contacts within the binder chain. **(E)** There is no trend for pi-pi contacts between the binder and CD20. Significance from two-sided BH-corrected Mann-Whitney-Wilcoxon tests; ns, not significant; *, p ≤ 0.05; **, p ≤ 0.01; ***, p ≤ 0.001; ****, p ≤ 0.0001.

**Figure SA12.**
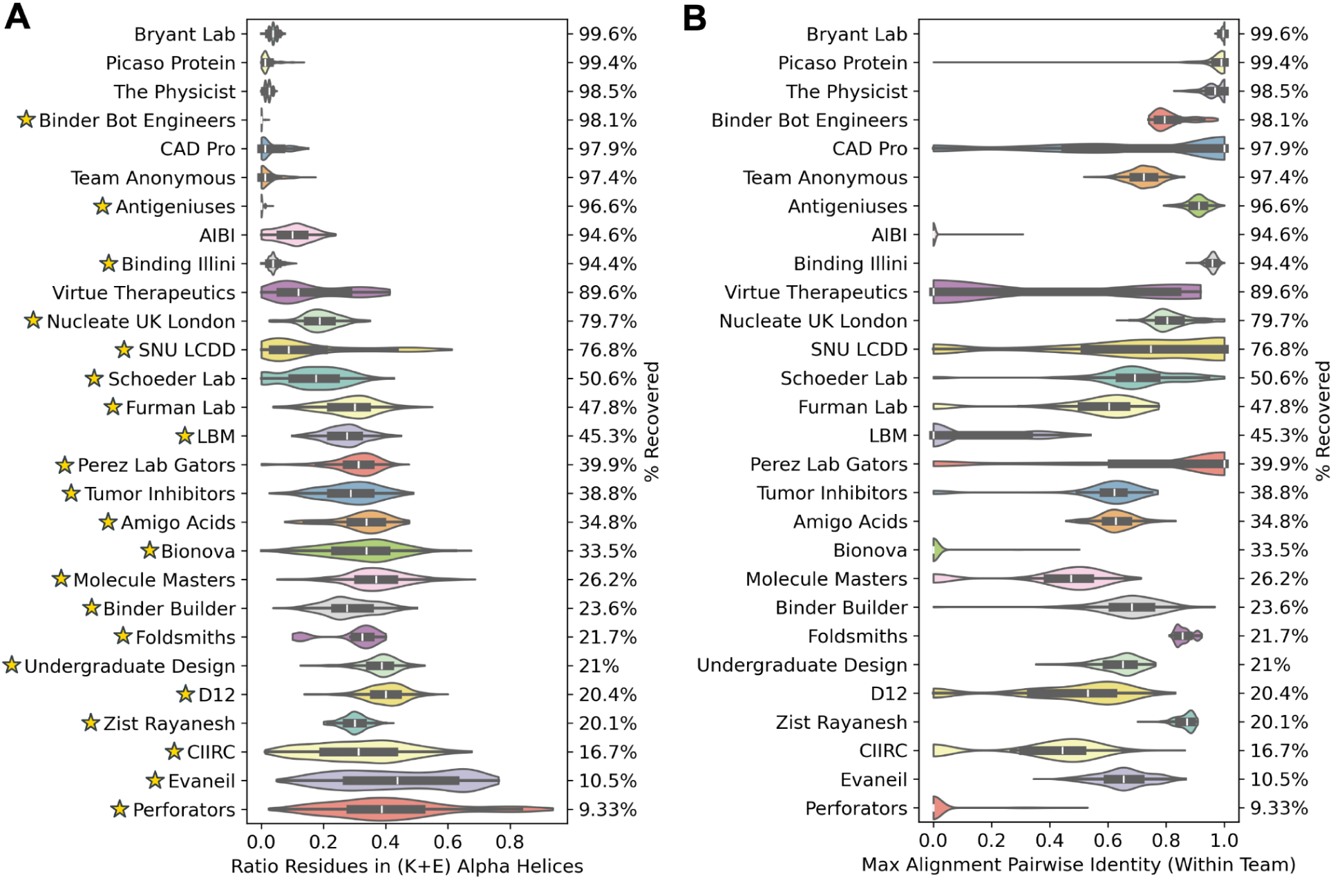
Team-wise breakdown of sequence recovery. **(A)** Sequence recovery by ratio of residues in K+E alpha helices. Teams that used ProteinMPNN or SolubleMPNN are denoted by stars. **(B)** Sequence recovery by sequence diversity calculated by the maximum MMseqs2 alignment pairwise similarity to a sequence from the same team (higher = more redundant; lower = more unique). Several of the teams with high sequence recoveries have redundant sequences, differing by only a few residues. However, several teams achieved high sequence diversity with substantial diversity between designs.

**Table SA1.** Best features for predicting experimental readouts. For each of four experimental outcomes, selected high-performing features are shown. Each feature was used to train a logistic regression model (max_iter=1000, 5-fold random split) to predict the given class. Similar to above, negative samples were filtered out at each step. Set sizes are as follows: DNA detected: 12K total; LEAH Recovered: 11796 total (removed DNA not detected); ≥2-fold increase and ≥2-fold decrease: 6811 (removed both DNA not detected and LEAH not recovered). Values are ROC-AUC scores across the 5 splits, sorted by ≥2-fold increase.

| Metric | DNA detected | LEAH recovered | $\geq 2$-fold increase | $\geq 2$-fold decrease |
| --- | --- | --- | --- | --- |
| ratio_KE | 0.617 | 0.896 | **0.725** | 0.620 |
| dssp_ke_alpha_ratio | 0.638 | **0.905** | 0.700 | 0.648 |
| num_AA_repeats_dna | 0.588 | 0.882 | 0.681 | 0.567 |
| num_EE_repeats_aa | 0.625 | 0.818 | 0.670 | 0.596 |
| num_cysteines | 0.561 | 0.683 | 0.663 | 0.738 |
| dna_sequence_entropy | 0.790 | 0.897 | 0.660 | 0.658 |
| aa_sequence_entropy | 0.795 | 0.833 | 0.626 | **0.757** |
| boltz_autoregwithseq_protein<br>mpnn | 0.536 | 0.665 | 0.630 | 0.729 |
| esm2_binder_LL_car_target | 0.763 | 0.834 | 0.597 | 0.710 |
| seq_kl_vs_human | **0.803** | 0.828 | 0.578 | 0.618 |
| boltz_A_BC_max_ipSAE | 0.555 | 0.517 | 0.519 | 0.489 |
| dna_gc_content | 0.568 | 0.625 | 0.519 | 0.645 |
| boltz_rosetta_complex_normalized | 0.493 | 0.570 | 0.499 | 0.463 |
| boltz_ipTM | 0.555 | 0.525 | 0.505 | 0.457 |
| boltz_ipLDDT | 0.570 | 0.662 | 0.570 | 0.516 |
| boltz_A_BC_max_lis_score | 0.533 | 0.532 | 0.515 | 0.503 |
| boltz_sap_score | 0.697 | 0.791 | 0.548 | 0.558 |
| boltz_A_BC_max_pDockQ | 0.546 | 0.636 | 0.553 | 0.629 |

**Figure SA13.**
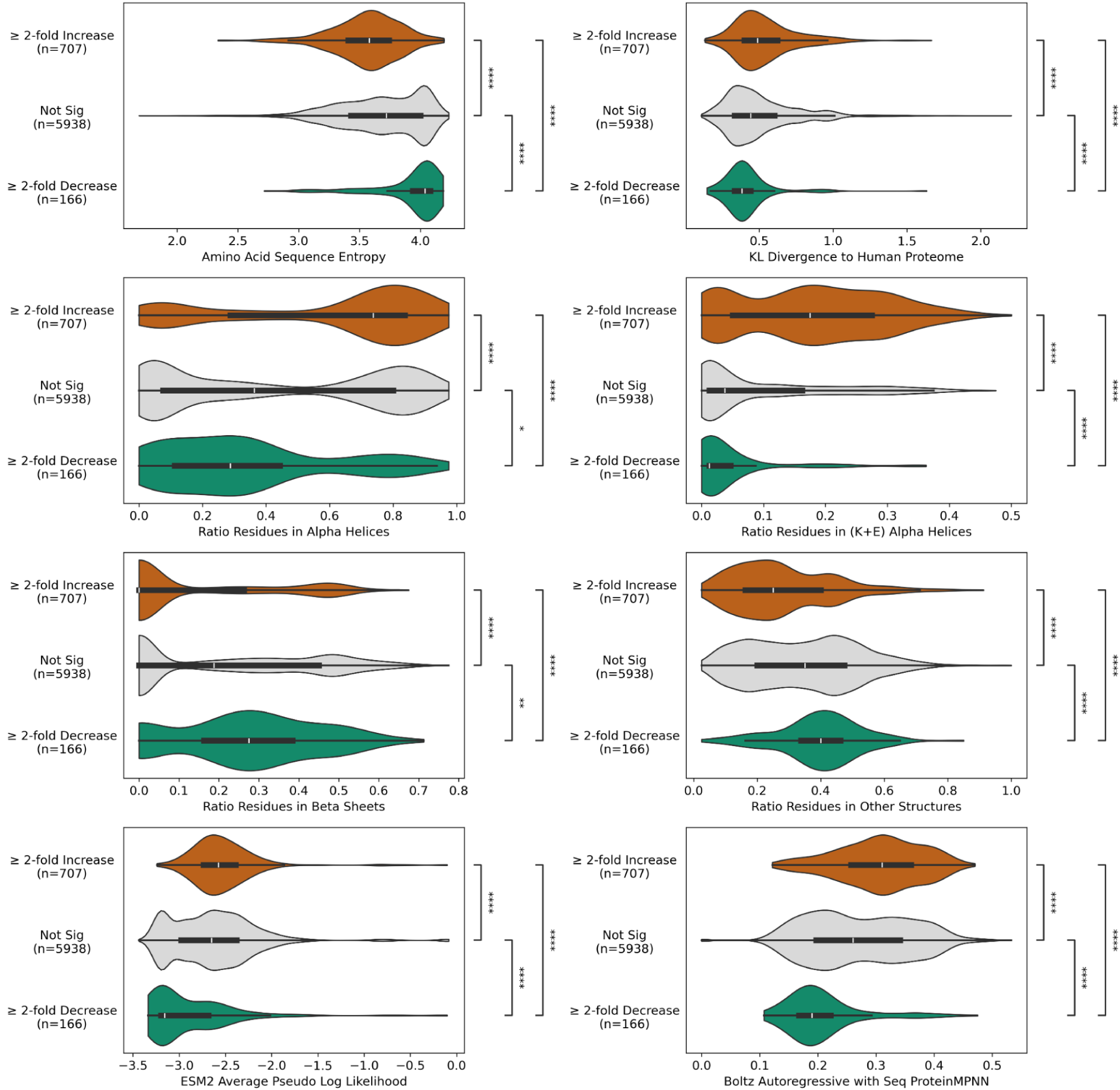
Additional differences between proliferation outcomes. Significance from two-sided BH-corrected Mann-Whitney-Wilcoxon tests; ns, not significant; *, p ≤ 0.05; **, p ≤ 0.01; ***, p ≤ 0.001; ****, p ≤ 0.0001.

**Figure SA14.**
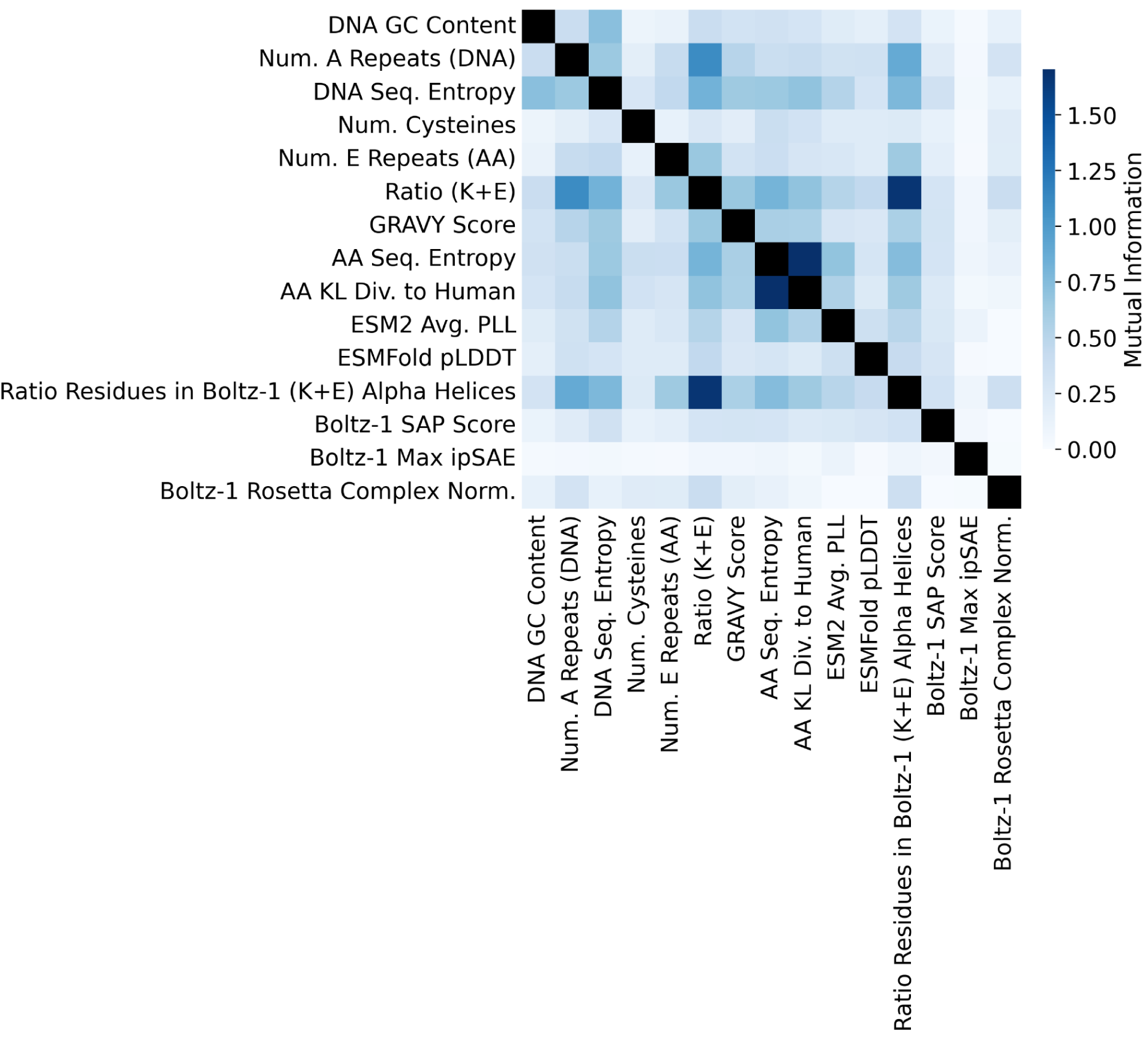
Mutual information between selected metrics. Mutual information was computed across metrics from all 12,000 sequences, demonstrating correlation between the different features.

**Figure SA15.**
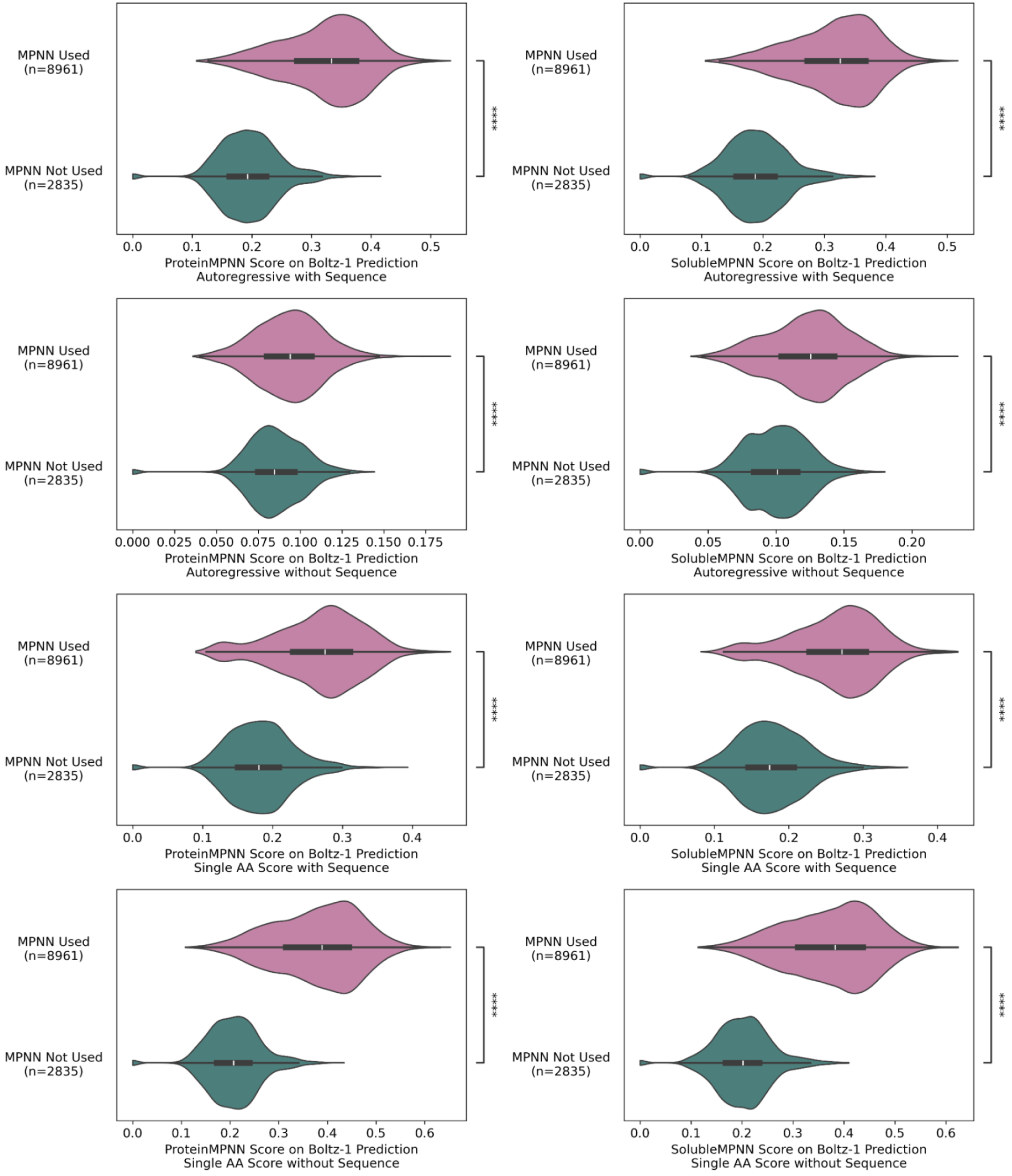
ProteinMPNN and SolubleMPNN scoring is generally biased toward sequences designed by said models. ProteinMPNN and SolubleMPNN scores were obtained from the predicted Boltz-1 binder:CD20 structures. The ProteinMPNN scoring setting “autoregressive without sequence” has the least sensitivity to the use of MPNN, suggesting that this should be used over the others when the scored proteins were designed with and without an MPNN model. Significance from two-sided BH-corrected Mann-Whitney-Wilcoxon tests; ns, not significant; *, p ≤ 0.05; **, p ≤ 0.01; ***, p ≤ 0.001; ****, p ≤ 0.0001.

**Figure SA16.**
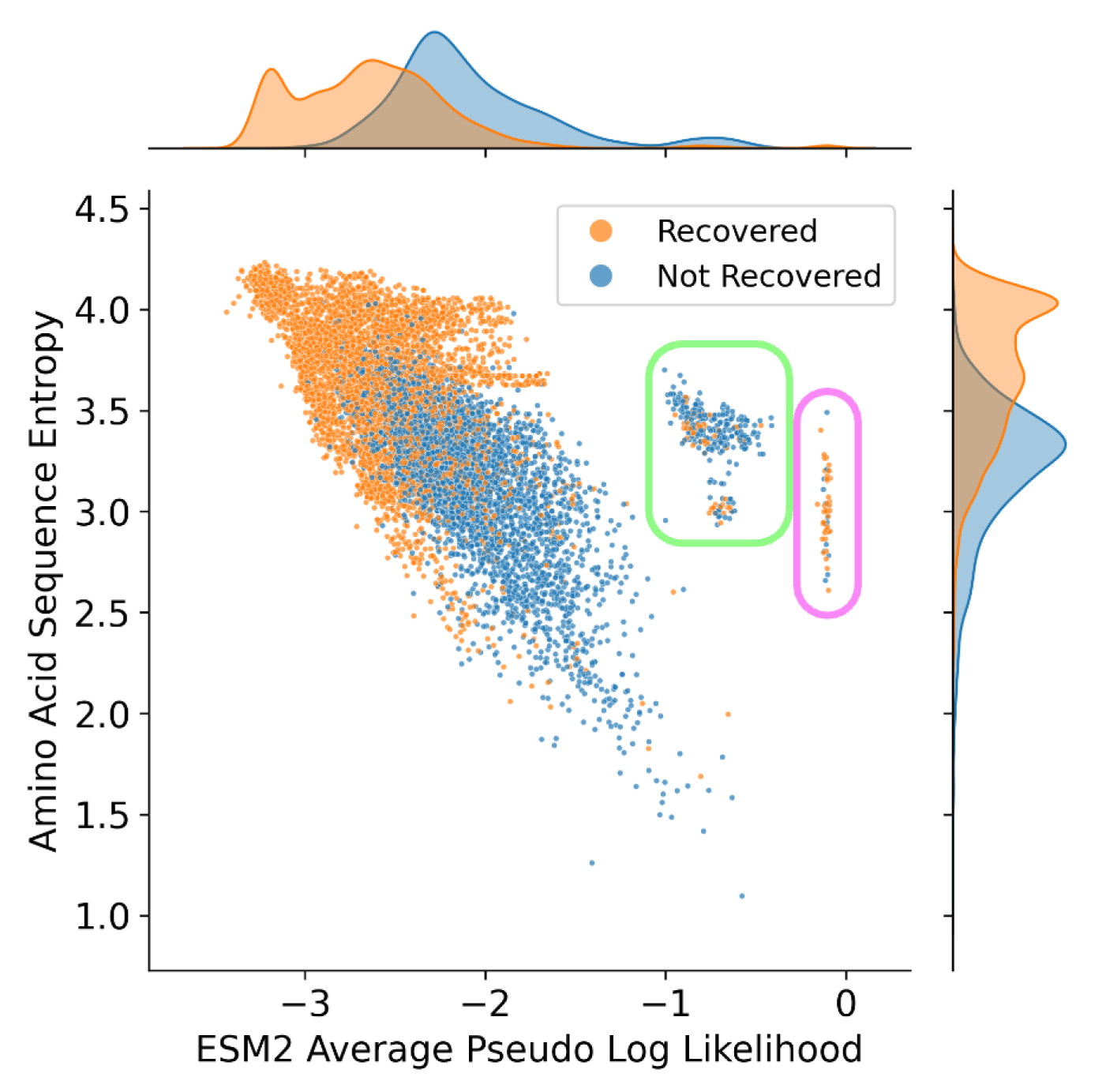
ESM-2 Pseudo Log Likelihood (PLL) artifacts and entropy correlation. ESM-2 PLL (0.85 ROC-AUC) generally correlates with the Shannon entropy of the amino acid sequence, increasing as sequence entropy decreases. However, two clusters defy this trend. The green region contains only sequences from a single team, in which all 268 sequences had duplicated segments of length 29 or more, with 58-70 residues being part of a repeated substring pair. The far right vertically linear cluster, within the pink region, contains only sequences from a single other team, in which all 60 sequences had 3-4 linkers, with 60-80 of the design’s residues being part of a repeated substring pair. Together, these explain the deviations from the ESM-2 PLL trend. This metric is computed by masking out one residue at a time, a task that becomes trivially easy when the answer is repeated elsewhere in the sequence. In agreement with existing literature^61^, caution should be exercised when using PLL from a protein language model to down-select sequences, as they may result from syntactic sequence artifacts rather than evolutionary likelihood.

### B. Computational Metrics

From the 12K binder sequences, a variety of *in silico* metrics were computed. Many metrics were computed using predicted structures from Boltz-1^67^, Chai-1^31^, and ESMfold^21^. To avoid modeling long disordered regions, a truncated CD20 sequence from PDB:6VJA was used, spanning residues 46 through 217.

**Table SB1.** Sequence-derived metrics.

| **Metric Name** | **Description** |
| --- | --- |
| DNA sequence entropy | Shannon entropy of the DNA sequences |
| DNA GC content | Ratio of guanine and cytosine in binder DNA sequence |
| Amino acid sequence entropy | Shannon entropy of the AA sequences |
| Number of cysteines | Number of cysteines in the binder sequence |
| Ratio of KE residues | Ratio of lysine and glutamate to total |
| Amino Acid composition KL divergence to human proteome | KL divergence between the binder's amino acid composition and that of the human proteome |
| Predicted charge | Computed with Bio.SeqUtils.IsoelectricPoint at pH 7 |
| Max percent identity within team (Hamming distance) | Highest percent identity, calculated using Hamming distance, between a sequence and all other sequences from the same team. Measure of sequence uniqueness |
| Max percent identity cross team (Hamming distance) | Highest percent identity, calculated using Hamming distance, between a sequence and all other sequences from other teams. Measure of sequence uniqueness |
| Max percent identity within team (MMseqs2 alignment) | Highest percent identity, calculated using MMseqs2, between a sequence and all other sequences from the same team. Measure of sequence uniqueness |
| Max percent identity cross team (MMseqs2 alignment) | Highest percent identity, calculated using MMseqs2, between a sequence and all other sequences from the same team. Measure of sequence uniqueness |
| MMseqs2 MSA depth | Number of sequence entries found through MMseqs2 |
| RNN solubility | Solubility prediction (0.0 to 1.0), using peptide.bio's RNN model |
| ESM-2 Full LL Alone | Log-likelihood for binder alone |
| ESM-2 Full LL CAR | Log-likelihood for CAR + binder |
| ESM-2 Binder LL CAR | Binder-specific log-likelihood with CAR context |
| ESM-2 Full LL CAR Target | Log-likelihood for CAR + binder + target |
| ESM-2 Binder LL CAR Target | Binder log-likelihood with CAR + target context |
| ESM-2 Full LL Target | Log-likelihood for binder + target |
| ESM-2 Binder LL Target | Binder log-likelihood with target context |

Multiple sequence alignments (MSAs) were computed across all binders using MMseqs2 to UniRef30 and ColabFoldEnv databases with default Colabfold MMSeqs server parameters. 10,329 of the 12,000 binders had zero MSA depth. Boltz-1 was used to generate structures, using the Boltz MSA server whenever MMseqs2 failed to return a valid output. Rosetta FastRelax was applied to each Boltz pose prior to Rosetta InterfaceAnalyzer.

**Table SB2.** Boltz-1 structure-derived metrics.

| Metric Name | Description |
| --- | --- |
| DSSP secondary structure metrics | Dictionary of Secondary Structure in Proteins (DSSP) was used to assign membership to alpha helices, beta sheets, and other secondary structure elements |
| Ratio of residues in K+E alpha helices | The ratio of residues that are lysine or glutamate and are found in an alpha helix. This was found to be highly correlated with NGS recovery after growth, and is also indicative of ProteinMPNN. |
| Hotspots engaged | Ratio of residues in large and small loops of CD20 that are engaged by the binder. |
| Hotspot PAE, iPLDDT, PDE | Values from Boltz matrix subsetting at the hotspot positions. |
| SAP Score | Spatial Aggregation Propensity, computed from Rosetta. |
| Rosetta InterfaceAnalyzer | All metrics from Rosetta InterfaceAnalyzer were computed: https://docs.rosettacommons.org/docs/latest/application_documentation/analysis/interface-analyzer |
| Angle of binder N/C termini relative to central axis of CD20 | This was computed to check whether geometric constraints determine binder success. |
| Binder location relative to membrane | Binder's COM displacement from origin. Positive means outer membrane. |
| ipSAE<sup>55</sup> | https://github.com/DunbrackLab/IPSAE |
| LIS score<sup>56</sup> | https://github.com/flyark/AFM-LIS |
| pDockQ<sup>59</sup> | https://www.nature.com/articles/s41467-022-28865-w |
| pDockQ2<sup>60</sup> | https://academic.oup.com/bioinformatics/article/39/7/btad424/7219714 |
| Number of predicted disulfide bonds (within chain) | Predicted disulfides within the binder chain only. Disulfides were found by collecting all CYS residues with an SG atom, checking every pair's SG-SG distance, and recording pairs within 3.0 Å |
| Number of predicted disulfide bonds (between chains) | Predicted disulfides between the binder and CD20. Disulfides were found by collecting all CYS residues with an SG atom, checking every pair's SG-SG distance, and recording pairs within 3.0 Å |
| Number of predicted pi-pi interactions (within chain) | Predicted pi-pi interactions within the binder chain only. Pi-pi was found by collecting aromatic residues (PHE/TYR/TRP/HIS), computing each ring's centroid and plane normal from selected ring atoms, and recording residue pairs whose centroids are within 7.0 Å and whose ring-plane normals are within 30° |
| Number of predicted pi-pi interactions (between chains) | Predicted pi-pi interactions between the binder and CD20. Pi-pi was found by collecting aromatic residues (PHE/TYR/TRP/HIS), computing each ring's centroid and plane normal from selected ring atoms, and recording residue pairs whose centroids are within 7.0 Å and whose ring-plane normals are within 30° |
| ProteinMPNN autoregressive with sequence | Likelihood derived from ProteinMPNN using:<br>-autoregressive 1<br>-use_sequence 1 |
| ProteinMPNN autoregressive without sequence | Likelihood derived from ProteinMPNN using:<br>-autoregressive 1<br>-use_sequence 0 |
| ProteinMPNN single AA with sequence | Likelihood derived from ProteinMPNN using:<br>-single_aa_score 1<br>-use_sequence 1 |
| ProteinMPNN single AA without sequence | Likelihood derived from ProteinMPNN using:<br>-single_aa_score 1<br>-use_sequence 0 |
| SolubleMPNN autoregressive with sequence | Likelihood derived from SolubleMPNN using:<br>-autoregressive 1<br>-use_sequence 1 |
| SolubleMPNN autoregressive | Likelihood derived from SolubleMPNN using: |
| without sequence | –autoregressive 1<br>–use_sequence 0 |
| SolubleMPNN single AA with sequence | Likelihood derived from SolubleMPNN using:<br>–single_aa_score 1<br>–use_sequence 1 |
| SolubleMPNN single AA without sequence | Likelihood derived from SolubleMPNN using:<br>–single_aa_score 1<br>–use_sequence 0 |

Given that ESMFold does not natively multimer input, we introduced a shift in the positional encoding and/or a glycine linker. We tested different chain permutations depending on the position of the binder (start, middle, end); as well as with or without the glycine linker (mode_glycine_linker). The following scores were computed on these structures.

**Table SB3.** ESMFold structure-derived metrics.

| Metric Name | Description |
| --- | --- |
| pTM, iPAE, iPTM, pLDDT | Common folding metrics |
| Binder location relative to membrane | Binder's COM displacement from origin. Positive means outer membrane. |
| ipSAE | https://github.com/DunbrackLab/IPSAE |
| LIS score | https://github.com/flyark/AFM-LIS |
| pDockQ | https://www.nature.com/articles/s41467-022-28865-w |
| pDockQ2 | https://academic.oup.com/bioinformatics/article/39/7/btad424/7219714 |

**Table SB4.** Chai-1 structure-derived metrics.

| Metric Name | Description |
| --- | --- |
| pTM, iPAE, iPTM, pLDDT | Common folding metrics |
| Binder location relative to membrane | Binder's COM displacement from origin. Positive means outer membrane. |
| ipSAE | https://github.com/DunbrackLab/IPSAE |
| LIS score | https://github.com/flyark/AFM-LIS |
| pDockQ | https://www.nature.com/articles/s41467-022-28865-w |
| pDockQ2 | https://academic.oup.com/bioinformatics/article/39/7/btad424/7219714 |

**Table SB5.**
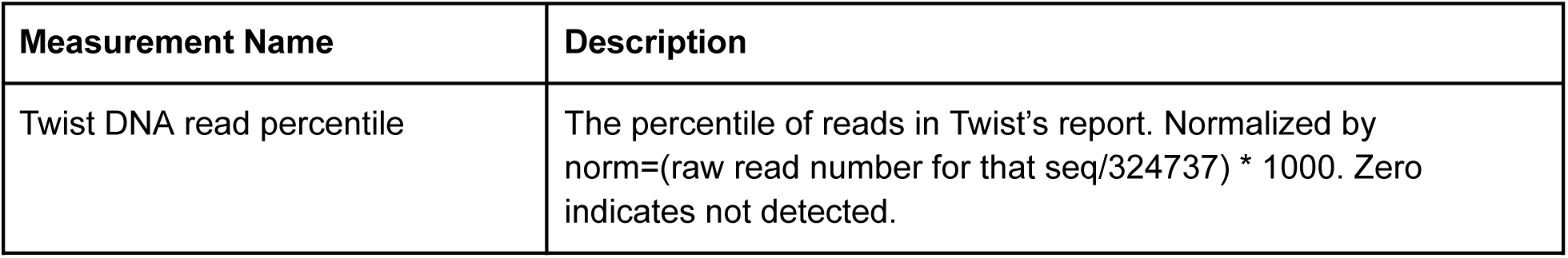
Twist measured values.

**Table SB6.** LEAH 12K proliferation values.

| Measurement Name | Description |
| --- | --- |
| K: CD20- NGS reads (x3) | NGS counts after proliferation without presence of CD20+ cells |
| R: CD20+ NGS reads (x3) | NGS counts after proliferation in presence of CD20+ cells |
| logFC | $\log_2(R/K)$ |
| logCPM | log counts per million |
| FDR | False discovery rate |
| Recovered | <25 reads in any of the six replicates |
| Fourclass | Up: $\geq 2$-fold logFC and significant based on FDR<br>Down: $\leq 2$-fold logFC and significant based on FDR<br>Not Sig: Not significant based on FDR<br>Not Recovered: <25 reads in any of the six replicates |
| Final score | $-\log_{10}(FDR) * \log FC$ |

**Table SB7.**
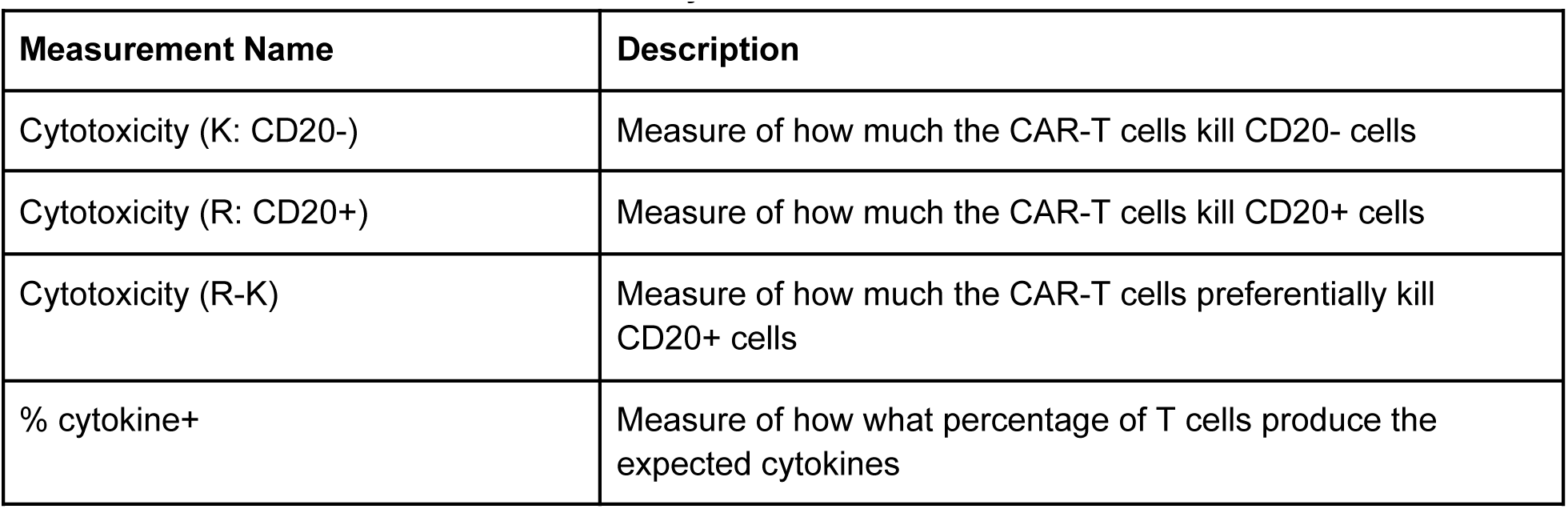
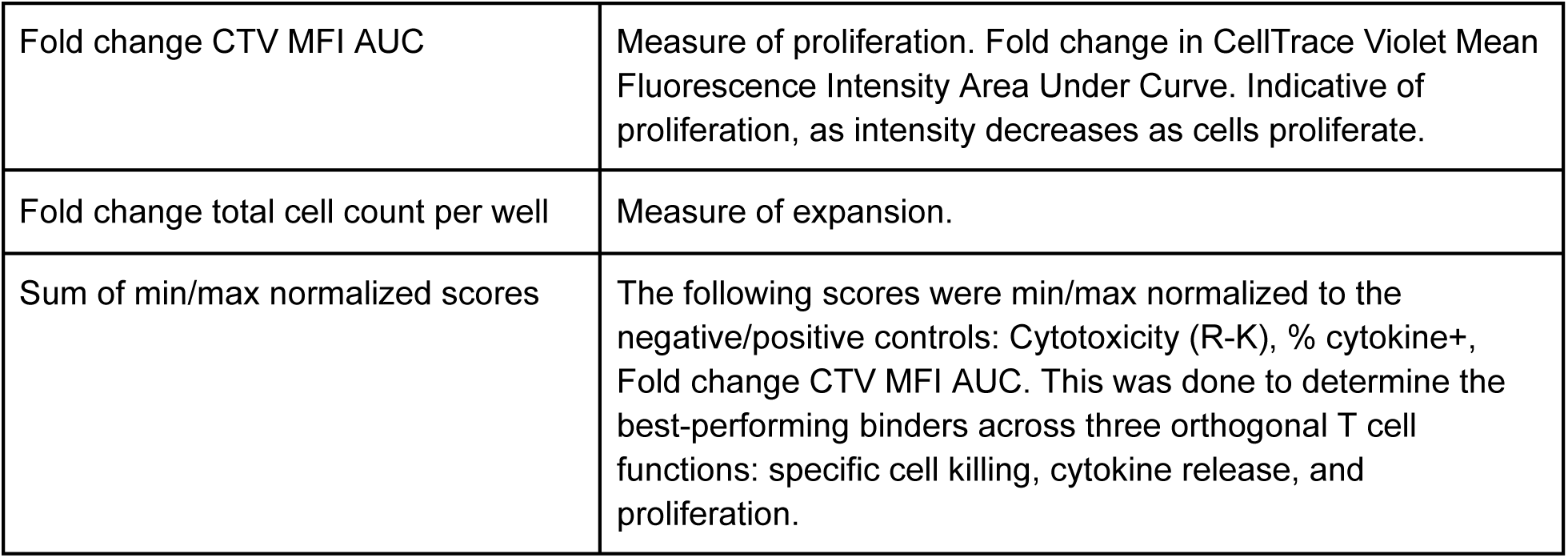
LEAH individual functional assays.

### C. Competition Information, Discussion, and Lessons Abbreviated timeline

May 2024: Initial planning begins

June 2024: Sponsor acquisition

July 16th, 2024: Public announcement

August 24th, 2024: Kickoff event

September 31st, 2024: Submission deadline (original)

October 7th, 2024: Submission deadline (post-extension)

October 2024: Sequences ordered

Jan 30th, 2025: DNA arrived from Twist to UT Austin. Shipped to LEAH Labs

May 2025: 12K proliferation results announcement

August 2025: Functional assays for top 10 completed

September 2nd, 2025: Winning teams announced

November 2025: Adaptyv begins binding affinity study

December 2025: Adaptyv completes binding affinity study

## Entrant statistics

**Figure SC1.**
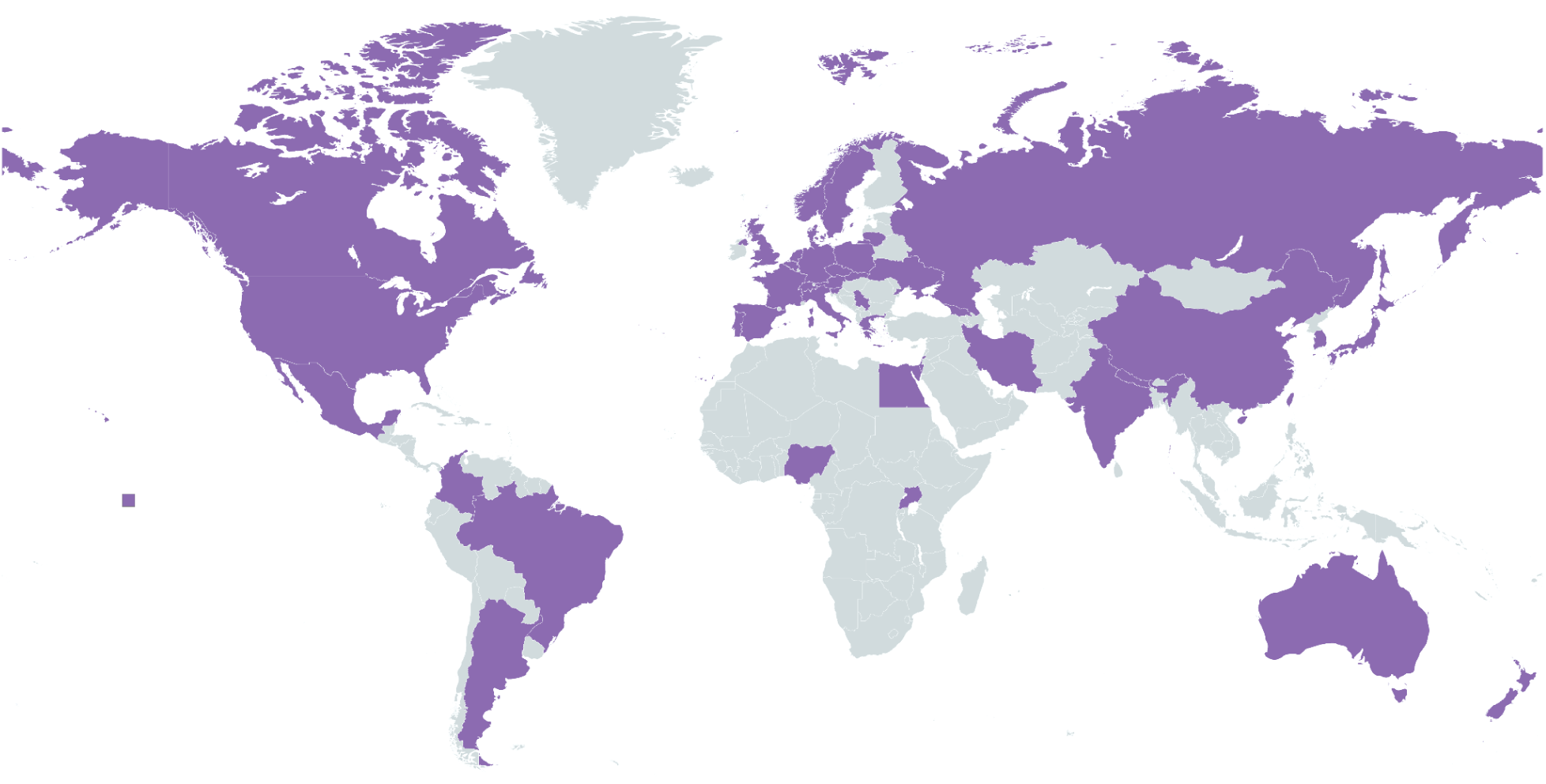
Countries of citizenship reported by competition entrants. Entrants span 42 countries across 6/7 continents. Data collected from the 169 competition RSVPs and the team formation responses. Map created with mapchart.net.

**Figure SC2.**
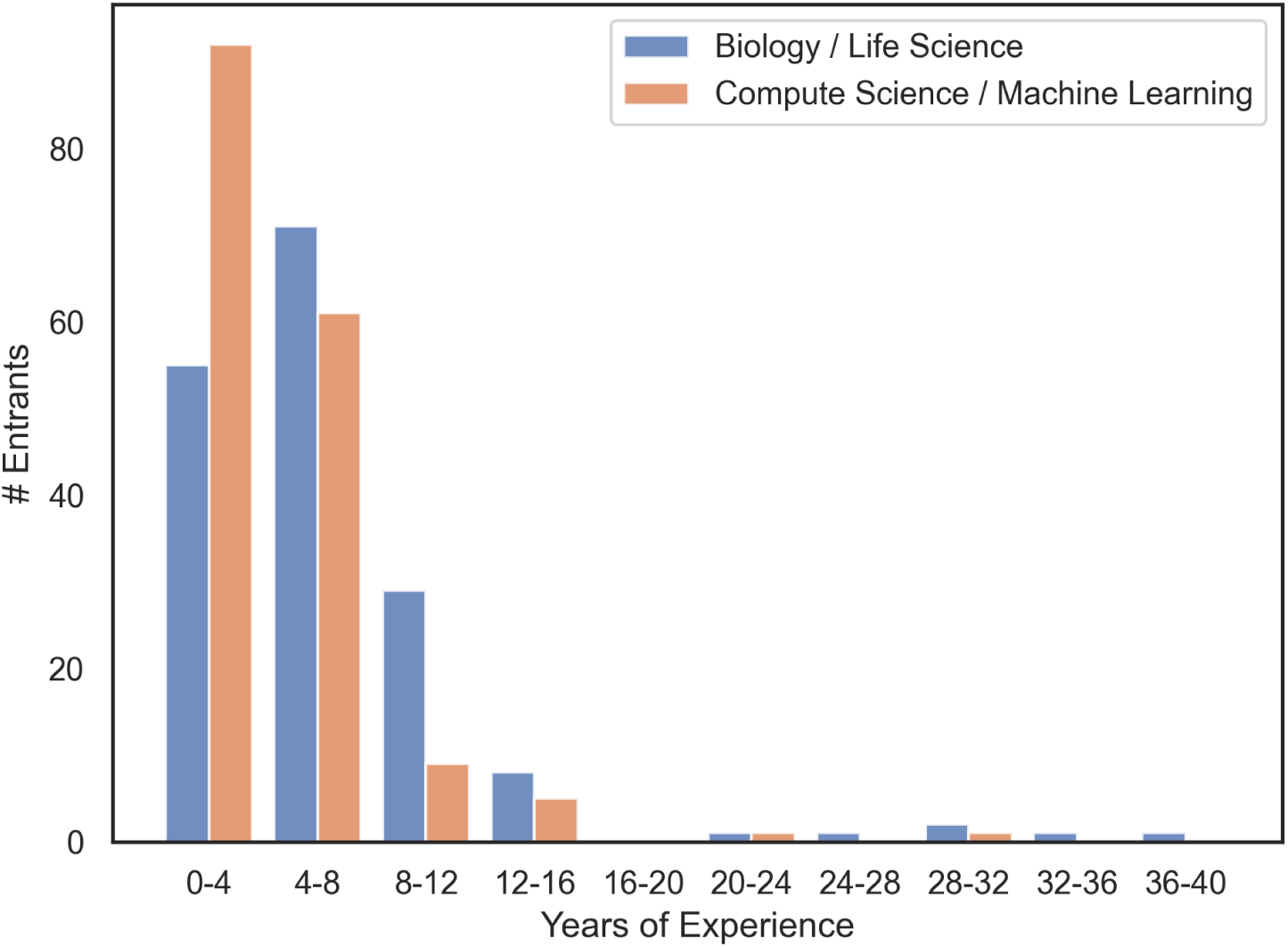
Competition entrant years of experience. Data collected from the 169 competition RSVPs and the team formation responses.

### Intellectual property considerations and avoiding conflicts of interest

Given the fact that competitors from academic and commercial entities were designing potential therapeutics validated by commercial entities, we required a very clear stance on how intellectual property would be handled. After considering the various options, it was deemed that the best option was for everyone to release all claims on intellectual property, and that LEAH Labs would not commercialize any of the submitted sequences. This was further complicated by some models having licenses that forbid commercial entities from testing the created proteins. To avoid any conflicts, competitors were required to acknowledge and agree to a participant agreement that explicitly addressed both intellectual property rights and the use of commercial software. This agreement was as follows:

Participant Agreement:

1. Intellectual Property: No intellectual property (IP) rights will be claimed on the designed sequences. All designed sequences must be released to the public domain. The organizers and sponsors will not be claiming intellectual property rights to the designed sequences nor will they be commercializing them.
2. Open Source and Public Availability Requirement:

a. All datasets, models, methods, and code developed or used during the competition should be made open source whenever possible.
b. We discourage teams from using closed-source and license-restrictive models such as AlphaFold3 and ESM-3. However, we cannot police this given the number of submissions we are dealing with, so by submitting your designs you agree that the burden of responsibility for all software used rests upon you, the submitting party, and not the organizers or sponsors.

Further, the participant represents that they have the right to use any software, models, tools or data that they have used or will use to produce any entry in the competition. Participant also represents that Participant (i) has not filed a patent application or submitted an invention disclosure to Participant’s employer or institution for the purpose of a patent application being filed, and (ii) is not aware of any invention disclosure or patent application, that attempts to protect or cover the sequence, sequences, molecule or molecules, or any method of using them, that Participant has submitted or will submit as an entry in the competition. Participant further represents that Participant will not in the future file such an application or submit such an invention disclosure to an employer or institution.

### Kickoff event

A 2-hour remote kickoff event was held on Saturday August 24th, 2024. The purpose of this event was to formally reveal the design target, explain the logistics in more detail, and more. The event proceeded with the following schedule:

● Motivation for the competition
● Sponsor talks: Lonza Group, ScaleReady, Twist Bioscience, LEAH Labs
● Problem statement, target reveal, CAR-T background information
● Submission requirements, IP considerations
● Experimental validation overview and winners selection process
● Example binder design workflow

● Compute options
● Closing remarks
● Office hours

Slides: https://github.com/kosonocky/bits-to-binders/blob/main/misc/kickoff_slides.pdf Recording: https://www.youtube.com/watch?v=fSoEX0c-Gko

### Resources provided to competitors

A compilation of resources was provided to the competitors after the kickoff event. This included papers, code, and web-servers for: protein structure prediction, inverse folding, generative models, language models, and other relevant tools. Additional papers and lectures were provided on computational protein design, along with several reviews on CAR-T therapies and CD20. Links to guides on using TACC, Synthia, and additional tools suggested by Adaptyv Bio were additionally provided. These resources can be found at: https://github.com/kosonocky/bits-to-binders-resources.

### Compute support

Competitors were given access to compute at the Texas Advanced Computing Center (TACC) Vista GH200 supercluster. Many competitors did not use this resource as many models and open source software were not yet compatible and compiled for Vista’s ARM operating system. This highlights a need for computational resources to be aligned with the user expertise for usability.

### Miscellaneous logistics

Advertisement for the competition went out on the organizer’s social media platforms (LinkedIn and X), university communications, and communications from the competition’s sponsors. An RSVP form was sent out with the announcement. This form was used to collect competitor demographics and was used to add people to a Slack channel for further announcements. In this channel, people without teams were encouraged to reach out to one another and form teams on their own. Throughout the event, announcements were primarily made through this Slack channel, though occasional communications went out over email. The main disadvantage of using the free version of Slack for an event like this is that messages disappear after 90 days, meaning that past communications become inaccessible.

The kickoff event was hosted virtually, with organizers present in-person in Austin, Texas to ensure a smooth rollout. The event was on Saturday at the end of the summer prior to the beginning of the fall semester to improve attendance from both industry and academic participants.

### Determination of winning teams

From the 12K proliferation screen, ten of the top-performing non-redundant designs were chosen to proceed to a series of individual CAR-T functional screens. From these top ten, data on proliferation, expansion, cytotoxicity, and cytokine production were obtained. A final score was created from this data to best reflect overall performance without being biased to a given behavior. Given their probable correlation, proliferation was withheld from the final score and just expansion was retained. Values for cytotoxicity, cytokine, and expansion were min/max normalized to the negative/positive controls, and summed for the “final score”. Designs were then ranked by this final score, and the top three designs were deemed 1st, 2nd, and 3rd place. A fourth winner was also announced to be the team with the highest percentage of successful sequences in the 12K proliferation screen.

The top three teams by 12K screen hit rate were as follows:

1. *Nucleate UK London (38.4%)
2. Binding Illini (14.8%)
3. Furman Lab (11.1%)

The top three finalist teams by the min/maxed “final score” were as follows:

1. *Perez Lab Gators (1.99/3.00)
2. *Amigo Acids (1.95/3.00)
3. *Schoeder Lab (1.34/3.00)

Each winning team, designated with an asterisk, was gifted a 3D-printed trophy of their successful binder. For Nucleate UK London, a random successful binder was used.

**Figure SC3.**
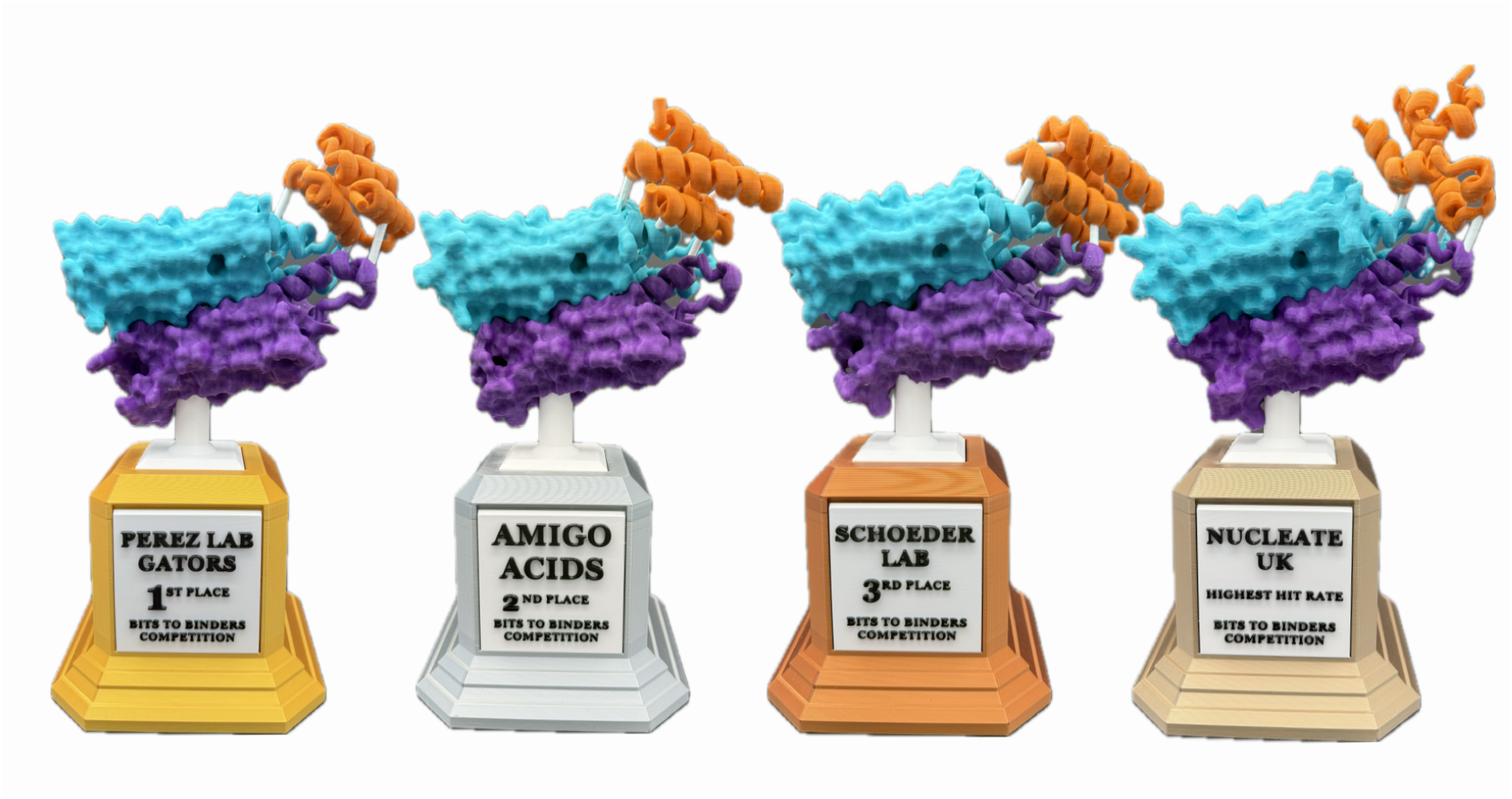
3D-printed trophies for the winning teams. Trophies were given to the three teams with the best individual sequences and to the team with the highest hit rate in the 12K proliferation screen.

### Feedback to organizers

Feedback was obtained from 14 teams that entered the competition. All of these teams except one submitted sequences, with the remaining one not submitting due to scheduling conflicts.

**Figure SC4.**
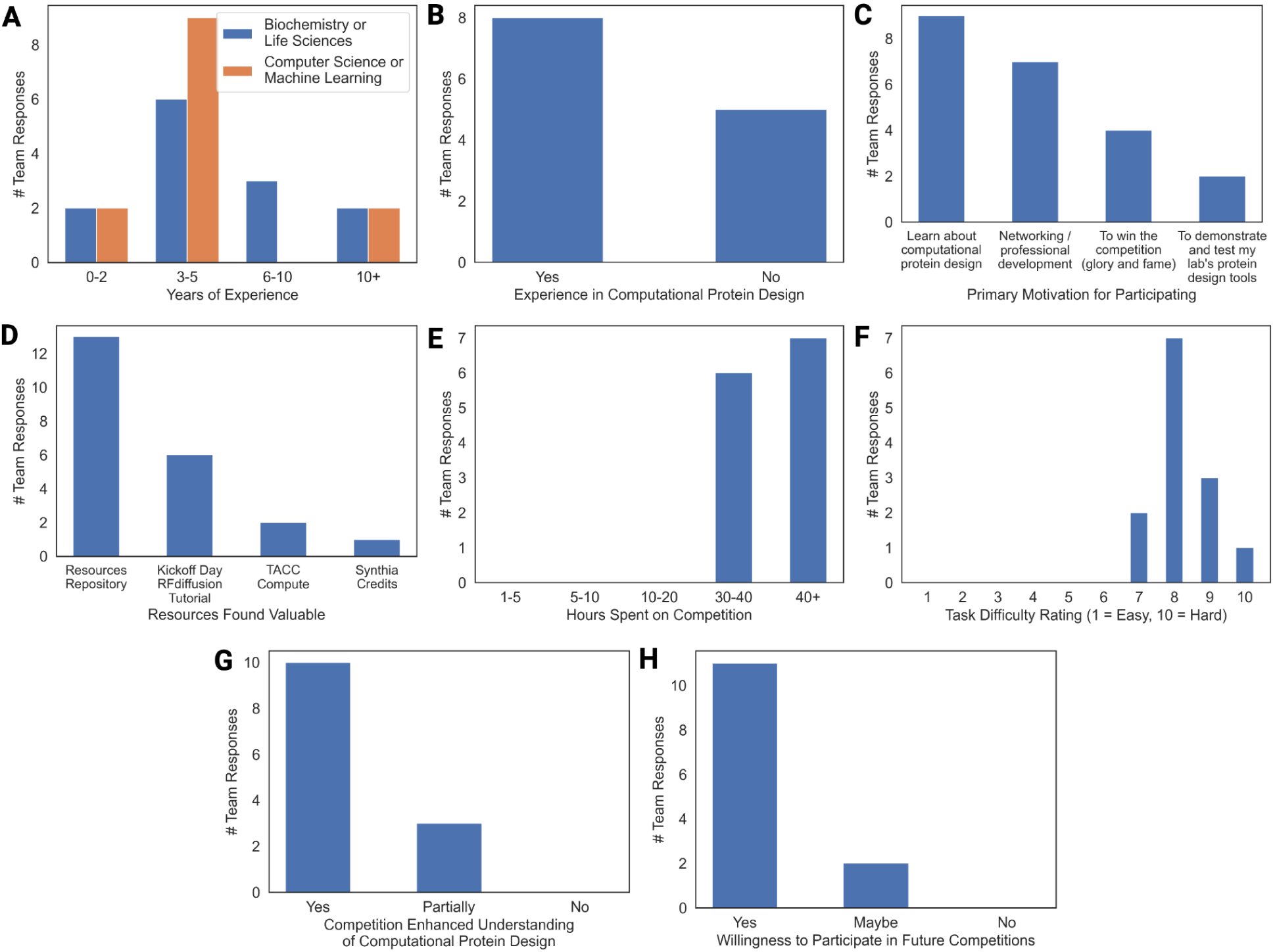
Survey response from 13 teams that submitted designs. **(A)** Years of experience in biochemistry, life sciences, computer science, and machine learning. **(B)** Previous experience in computational protein design. **(C)** Primary reason for entering the competition. Survey response from 13 teams that submitted designs. One other reason for entering was “I met all of the people on my team at a summer internship at the Baker lab. We wanted to stay in touch!!”. **(D)** Provided resources that were deemed valuable. **(E)** Hours spent on the competition. **(F)** Protein design task perceived difficulty. **(G)** Enhancement of participant understanding in computational protein design. **(H)** Participant interest in future competitions.

Additional Question 1. If [you didn’t find any of the provided resources useful], why? Which additional resources, if any, would have been useful?

“It was very difficult to coordinate and manage compute resources and the data between them”

“Easier setup process to access TACC computer resources, I was using custom scripts so didn’t need synthia credits, and I have used rfdiffusion before”

“AWS or GCP credits”

Additional Question 2. Did you find participating in the competition worthwhile for other reasons? Why or why not?

“I feel like it was extremely fun and challenging. I learned a lot about state of the art techniques for binding to hydrophilic surface proteins, something that I didn’t know a lot about before the competition.”

“It was great to have a fully remote networking opportunity, and to work on my teamwork skills.”

“Interesting to work on a target that one would usually avoid for binder design (small extracellular domain, no known native interaction)”

“Yes, collaboration with different individuals”

“Absolutely. I learned a lot of domain-specific knowledge that I wasn’t able to get.”

“I was testing my scientific collaboration platform to address the difficulties in scientific collaboration”

Additional Question 3. What aspects of the competition worked well?

“I think the overall competition and the instructions were very clear”

“Resources and kickoff”

“It’s awesome that you test so many of the designs directly in CARs”

“Discussion session and help session”

“timeline”

“Github repos, tutorials”

Additional Question 4. What changes or improvements would you like to see in future competitions?

“The process of finding a team in Slack was a little awkward. Maybe some clear guidance at the beginning about how to find teammates would have made this part a little smoother.”

“Better documentation for connecting to and using TACC resources with common models.”

“Easier to access resources (more user friendly), and more closing communication / followup”

“More office hours in regards to infrastructure”

“Sharing workflows between teams”

Additional Question 5. What topic would you like to see a competition focused on in the future?

“Enzymes!!!”

“Designing proteins/peptides for selective purification of lanthanide metals”

“Enzyme design”

“Enzyme activity, peptide binders”

“Spatial Transcriptomics that can be verified in the lab”

“Drug interaction, kinetic improvement of enzymes”

Additional Question 6. Do you have any additional feedback, comments, or suggestions?

“Do an enzyme competition. I think at this time next year the tech will be available and now that AF3 is out this would be such a cool contest!”

“Many thanks to the organizers for this great opportunity!”

“This was an awesome competition. I learned a lot!”

“A great competition idea, thank you for making it possible.”

### D. Sponsor Contributions

The *Bits to Binders* competition benefitted considerably from a number of academic and industrial sponsors. This competition would not have been possible without their generous support.

**LEAH Laboratories:**

Website:

https://www.leahlabs.com/

**Contribution:**

LEAH Labs helped plan and run the competition. Utilizing their patented precision integration technology developed in primary human T cells, LEAH Labs conducted pooled library integration and high-throughput screening as well as performed all experimental validation on T cells, and analyzed the experimental results.

**Adaptyv Bio:**

Website:

https://www.adaptyvbio.com/

Contribution:

Adaptyv Bio provided triplicate binding affinity measurements for 100 binders to CD20 free-of-charge.

**Twist Bioscience:**

Website:

https://www.twistbioscience.com/

Contribution:

Twist provided a 50% discount on the plasmid pools.

**The University of Texas at Austin Center for Systems and Synthetic Biology:**

Website: https://www.facebook.com/CSSBUT2015/

Contribution:

The UT Austin Center for Systems and Synthetic Biology provided funding to purchase the DNA and the trophies.

**Texas Advanced Computing Center:**

Website:

https://tacc.utexas.edu/

Contribution:

TACC provided free compute to competitors on their Vista GH200 superchip cluster.

**Modal:**

Website:

https://modal.com/

Contribution:

Modal provided $5K in compute credits that allowed J.L. to compute *in silico* metrics for the competition measurement data.

**Lonza:**

Website:

https://www.lonza.com/

Contribution:

Lonza provided reagents for CAR-T cell experiments.

**ScaleReady:**

Website: https://www.scaleready.com/

Contribution:

ScaleReady provided reagents for CAR-T cell experiments.

**VWR:**

Website:

https://www.vwr.com/

Contribution:

VWR provided reagents for CAR-T cell experiments.

**KUNGFU.AI:**

Website:

https://www.kungfu.ai/

Contribution:

KUNGFU.AI provided AI consulting to the competitors during the kickoff event.

**Maker Clinic:**

Website:

linkedin.com/in/adamsalmi

Contribution:

Maker Clinic provided custom 3D-printed trophies for the winning binders at a discounted rate.

**Synthia:**

Website:

https://www.synthialabs.com/

Contribution:

Synthia provided credits to competitors to use their protein design LLM.

**The University of Texas at Austin; Dell Medical School:**

Website:

https://dellmed.utexas.edu/

Contribution:

The UT Austin Dell Medical School helped advertise the competition and facilitate some of the connections to sponsors.

**Nucleate AI in Biotech:**

Website:

https://nucleate.org/

Contribution:

Nucleate AIxBio helped advertise the competition.

**The BioML Society:**

Website:

https://www.biomlsociety.org/

Contribution:

The BioML Society formulated, organized, and ran the competition.

1. E. *Bits to Binders* Teams, Competitors, and Design Methods

At the time of design submission, teams were asked to provide an abstract on the methods used as well as steps to reproduce the designs. Email addresses are listed for the designated primary team contacts. Questions regarding a team’s methods should be directed toward these individuals.

## 1. AIBI

Team Members:

Yang Shen, Shaowen Zhu, Yuxuan Liu, Rujie Yin, Dishant Parag Zaveri, Sai Jaideep Reddy Mure, Pragati Naikare

Abstract:

We mainly used a joint sequence-structure co-design model, which is a diffusion model conditioned on target CD20 structure & epitope, without or with the fixed context’s sequence and structure of the designed protein. We made multimer inference using three diffusion model variants and monomer inference with linkers across chains using the last variant. And we combined the four types of model inferences with five types of epitopes (three for known CD20 antibodies, the union of the three, and the entire extracellular domain) and generated 180 designs for each of the 20 combinations. The resulting 3600 designs are geometrically ranked based on contact consistency with intended epitopes. We also used a pretrained sequence-alone ESM2^21^ to impute the 80 amino acids from the context sequence and picked the top 50 from the 200 designs based on PIPR-predicted^68^ affinity to CD20.

Reproducibility:

Method A: Main method, no citation yet, designs 1-270 and 301-480.

A1. Joint sequence-structure generation using target (and scaffold) conditioned diffusion models

A2. Model selection using curated cocrystal structures between CD20/CD19 and antibodies

A3. Filtering and ranking designed structures based on consistency with desired contacts

Method B: ESM2 (650M parameters), designs 271-300 and 481-500.

B1. Sequence imputation from the rest of CAR sequences by iterative sampling of a pretrained ESM2, which is not specific to CD20.

B2. Filtering and ranking designed sequences based on sequence-predicted binding affinity to CD20

## 2. Amigo Acids

Team Members:

Antonio Fonseca, Joseph Openy, Ava Chan, Freddie Martin, Ansar Ahmad Javed, Dionessa Biton, Shreyasi Das, Francisco Requena

Abstract:

We used a combination of free diffusion with RFdiffusion^16^ constrained by length and using hotspots located on the extracellular portion of the CD20. With the designs that showed reasonable binding affinity, we performed partial diffusion followed by FastRelax to obtain new proteins that could bind more effectively. The ranking of the design was done with AF2^29^.

Reproducibility:

- Free diffusion with RFdiffusion (80 AAs) using hotspots
- ProteinMPNN^23^ to obtain sequences
- AFPulldown^69^ for structure assessment
- Sort designs by ipae and iptm
- Perform partial diffusion with RFdiffusion
- Predict sequences using ProteinMPNN+FastRelax
- Predict structure using ColabFold^30^

## 3. Antigeniuses

Team Members:

Alia Clark-ElSayed, Satoshi Ishida, Daniel Acosta, Tynan Gardner

Abstract:

We generated a list of protein sequences that are known to bind to CD20. Due to the fact that this was over 500 sequences, we wanted to down-select to the top ∼10 binders. To do this we used AlphaFold-Multimer^70^ to co-fold the binder with CD20. Based on the model scores, we selected the top 10 sequences to use for further design and diversification. We diversified these structures with RFdiffusion and then generated the sequences using ProteinMPNN.

Reproducibility:

- Generated list of CD20 binders from databases
- AlphaFold Multimer to assess binding (https://www.biorxiv.org/content/10.1101/2021.10.04.463034v2)
- RFDiffusion (https://www.nature.com/articles/s41586-023-06415-8)
- ProteinMPNN (https://www.science.org/doi/10.1126/science.add2187)

## 4. Binder Bot Engineers

Team Members:

Genhui Zheng, Rajarshi Mondal, Calvin XiaoYang Hu, Aoi Otani, Aparna Sahu

Abstract:

In this binder design experiment, we employ two approaches—active learning (AL)-based methods and traditional strategies—for antibody sequence generation. First, these methods are used to generate initial template sequences, which are then filtered using binding affinity prediction tools to select promising candidates.

Next, from the filtered templates, we create a diverse mutational library by introducing amino acid mutations, insertions, and deletions. This library is further evaluated using stability and other protein property prediction tools to ensure viability and functionality.

Finally, the sequences are ranked based on their predicted binding affinity, stability, and additional relevant properties. The top 500 sequences are selected for submission.

Reproducibility:

Sequence Generation:

Utilize both active learning (AL)-based methods and traditional methods to generate initial antibody template sequences [1-2]^23,71^.

Binding Affinity Filtering:

Apply binding affinity prediction tools to filter and retain only the most promising sequences[3]^29^.

Library Creation:

Generate a diverse sequence library by performing mutations, insertions, and deletions on the filtered templates to explore a wide range of amino acid variants.

Property Evaluation:

Use tools to evaluate stability and other key protein properties (e.g., solubility, aggregation, immunogenicity) for the entire sequence library [4]^72^.

Ranking and Selection:

Rank sequences based on binding affinity, stability, and additional protein properties. Select the top 500 sequences for submission.

## 5. Binder Builder

Team Members:

Corey Howe

Abstract:

I used the standard binder design pipeline of RFDiffusion for backbone generation, Soluble MPNN for sequence design, and AF2 for validation. With some promising binders in hand, I diversified and further optimized them with sequence redesign of the non-interacting residues as well as in silico directed evolution with AF2.

Reproducibility:

-Trim down the dimer CD20 structure to remove most of the transmembrane domain

-Run RFDiffusion with default settings at different binder sizes to find good designs

-Run Soluble MPNN to design sequences to the RFD diffusion backbones

-Validate the sequence designs with AlphaFold2 multimer

-Optimize the top designs by sequence redesign of non-interacting residues with soluble MPNN

-Optimize the top designs by in silico directed evolution with AlphaFold2 (code is personal fork of ColabDesign)

-Rank by plddt, i_pae, i_ptm

## 6. Binding Illini

Team Members:

Diwakar Shukla, Diego Kleiman, Tanner Dean, Song Yin, Lin Zhu, Joseph Clark

Abstract:

After extracting interface residues from Rituximab bound to CD20 (PDB ID: 6VJA)^73^, we utilized Chroma^17^ and ESM to generate sequences preserving those interface amino acids. Then, we predicted the structures of all generated sequences using Colabfold and selected the most promising models based on pLDDT and ipTM. We carried out a design loop similar to EvoBind2^74^, where new sequences where generated using ProteinMPNN and structures were predicted using Colabfold. The structure prediction that minimized a loss function based on pLDDT, ipTM, iPAE, and radius of gyration was then used as input for the next round.

Reproducibility:

- Extract interface residues from Rituximab (<5 A away from CD20)
- Generate sequences of desired length by stitching together the interface residues using Chroma (conditioned on interface structure and sequence) and ESM (conditioned on interface sequence)
- Predicted structures of CD20-peptide complex using Colabfold
- Carried out iterative design loop, sequences were generated from ProteinMPNN and structures were predicted using Colabfold. Loss function was based on pLDDT, ipTM, iPAE, and radius of gyration.

## 7. Bionova

Team Members:

Xin Wang, Yunchao Liu, Zhaoqian Su, Tyler Derr, Rocco Moretti, Jens Meiler

Abstract:

We implemented a standard pipeline encompassing backbone generation, inverse folding, and filtering to tackle the challenge. For binder design with the RFdiffusion model, our focus was on generating binder backbones that specifically target epitope 3 of CD20, located in the extracellular region. A diverse subset of these backbones was selected for inverse folding using ProteinMPNN, producing multiple sequence designs for each backbone. AlphaFold2 was then used to validate the 3D structures of the sequence designs by comparing the RMSD score to the original backbone. Finally, the Chai-1 model predicted the complex formed between CD20 and the filtered binder sequences. After excluding complex predictions that failed to meet the correct binding site and extracellular constraints, we ranked the binder sequences based on their iptm scores output by Chai-1.

Reproducibility:

1. Backbone Generation via RFDiffusion [1]
2. Inverse Folding via ProteinMPNN [2]
3. Relaxed Filtering via AlphaFold2 Initial Guess [2]^75^
4. Final Validation of Chai [4]

## 8. Bryant Lab

Team Members:

Patrick Bryant, Qiuzhen Li

Abstract:

This study employed EvoBind-multimer, a novel in silico-directed evolution framework based on AlphaFold2, to design linear peptides targeting CD20. The peptides, ranging from 5 to 20 amino acids in length, were designed to function as molecular glues, bridging interactions between two proteins of interest. The CD20 sequence was extracted from a known protein complex structure, and the algorithm autonomously identified optimal binding sites for the designed peptides.

Reproducibility:

Obtain the CD20 sequence from the Structure of CD20 in complex with rituximab Fab (PDB: https://www.rcsb.org/structure/6vja).

1. 2. Utilize EvoBind-multimer, an in-house tool, for cyclic peptide design.
2. Set peptide length ranges from 5 to 20 amino acids.
3. Configure EvoBind-multimer^74^ to autonomously select optimal binding sites on CD20.
4. Run EvoBind-multimer for 1000 iterations per peptide design.
5. Execute 8 cycles per iteration.
6. Analyze resulting cyclic peptide designs for potential molecular glue function between ligase domains and CD20

## 9. CAD Pro

Team Members:

Aswini Javvadi, Daksh Joshi

Abstract:

In this study, we aimed to design novel peptide binders targeting the CD20 antigen by employing a multi-step workflow. Initially, sequences of 80 amino acids in length were generated using PEPMLM and ESM models with both unconditional and conditional generation methods. These sequences were subsequently filtered based on AlphaFold2 (AF2) metrics, including inter-residue predicted aligned error (iPAE), predicted local distance difference test (pLDDT), and Local Interaction Score (LIS) models, to assess structural quality. Additionally, binding affinity predictions were employed to further refine the selection process.

We then performed structure prediction and molecular docking to evaluate the interaction potential of these sequences with CD20. The best candidates were selected by focusing on those with strong interactions specifically at the epitope 3 region of CD20. Sequences showing the highest number of interactions were ranked as top candidates for further analysis.

The Final Data was further augmented with sequences generated using EvoBind2, going through the same filtration steps as above.

Reproducibility:

•Generated 80 length sequences of target based using PEP-MLM^20^

Chen T, Dumas M, Watson R, Vincoff S, Peng C, Zhao L, Hong L, Pertsemlidis S, Shaepers-Cheu M, Wang TZ, Srijay D, Monticello C, Vure P, Pulugurta R, Kholina K, Goel S, DeLisa MP, Truant R, Aguilar HC, Chatterjee P. PepMLM: Target Sequence-Conditioned Generation of Therapeutic Peptide Binders via Span Masked Language Modeling. ArXiv [Preprint]. 2024 Aug 11:arXiv:2310.03842v3. PMID: 37873004; PMCID: PMC10593082.

•Generated sequences using ESM2 https://huggingface.co/facebook/esm2_t36_3B_UR50D

•Used iPAE, PLDDT, IPTM as scoring functions to filter sequences along with binding affinity

Bennett NR, Coventry B, Goreshnik I, Huang B, Allen A, Vafeados D, Peng YP, Dauparas J, Baek M, Stewart L, DiMaio F, De Munck S, Savvides SN, Baker D. Improving de novo protein binder design with deep learning. Nat Commun. 2023 May 6;14(1):2625. doi: 10.1038/s41467-023-38328-5. PMID: 37149653; PMCID: PMC10163288.

•Structure prediction using alphafold2

•Hdock for predicted structures and interactions screening using foldX^35^

Yan, Y., Tao, H., He, J. et al. The HDOCK server for integrated protein–protein docking. Nat Protoc 15, 1829–1852 (2020). https://doi.org/10.1038/s41596-020-0312-x

•Evobind for augmenting with more sequences, going through the same steps as above for filtration. Li, Qiuzhen, Efstathios Nikolaos Vlachos, and Patrick Bryant. “Design of linear and cyclic peptide binders of different lengths only from a protein target sequence.” bioRxiv (2024): 2024-06.

## 10. CIIRC

Team Members:

Nikola Zadorozhny, Petr Kouba, Anton Bushuiev

Abstract:

We utilized RFDiffusion, an open-source state-of-the-art method, to generate protein structures targeting epitopes associated with current antibiotics. By employing the “Complex Model” highlighted in the publication, as well as beta model and guided diffusion via potentials, we aimed to design effective binders. Sequences were obtained using ProteinMPNN. Due to challenges in filtering and validation with standard pipeline (AF2 initial guess), we explored alternative tools like HelixFold3^32^ and AlphaFold-Multimer to assess some of our designs. Additionally, we generated designs using BindCraft^18^.

Reproducibility:

- Utilized RFDiffusion for protein binder generation, experimenting with beta models and guided diffusion using potentials [1].
- Focused on hot-spot selection from antibiotic-targeted epitopes.
- Designed sequences using ProteinMPNN [2].
- Encountered validation issues with standard pipelines like AlphaFold2 Initial Guess [3].
- Used HelixFold3 [4] and AlphaFold-Multimer [5] for alternative validation on some designs.
- Made additional designs using BindCraft [6].

## 11. D12

Team Members:

Tadej Satler, Roman Jerala, Tjaša Mlakar, Duško Lainšček, Ajasja Ljubetič

Abstract:

Our approach to protein binder design involved a scaffold-guided workflow combining RFdiffusion, ProteinMPNN, and AlphaFold2, with final filtering based on PyRosetta metrics. We generated binder blueprints, creating adjacency matrices for binders up to 80 amino acids long. RFdiffusion was then used to target specific regions of the hCD20 antigen, exploiting both symmetric and non-symmetric designs. Symmetric diffusion allowed us to take advantage of the target’s symmetry for more efficient binder generation. Binders were validated through sequence design with ProteinMPNN and structural verification via AlphaFold2, followed by optional redesign and refinement through partial diffusion.

Reproducibility:

https://github.com/kosonocky/bits-to-binders/blob/main/misc/d12_reproducibility.md

## 12. Evaneil

Team Members:

Neil Anthony, Even Kiely

Abstract:

The design of novel protein binders targeting CD20 involved a multi-step computational approach. Starting with the crystal structure of CD20 from PDB entry 6VJA, we extracted the extra cellular regions of CD20 only. Key hotspot residues were manually identified to guide the design of backbone conformations that could promote binding interactions in targeted regions. Using RFdiffusion, we generated diverse backbone conformations compatible with these hotspot constraints. These backbones were then used as scaffolds for sequence design using ProteinMPNN, which predicted optimal amino acid sequences to stabilize each backbone conformation while maintaining the desired binding properties. Predicted binder sequences were scored and ranked based on their predicted stability and binding potential.

Reproducibility:

- Extract CD20 from 6VJA, and crop ECM residues -> cd20ecm
- Manually select hotspot residues to promote backbone binder in different regions
- Run RFdiffusion
- Use ProteinMPNN to generate residue on each backbone

https://github.com/RosettaCommons/RFdiffusion

https://github.com/dauparas/ProteinMPNN

## 13. Foldsmiths

Team Members:

Harish Srinivasan, Rongqing Yuan, Rui Guo, Jesse Durham, Jimin Pei, Qian Cong, Jian Zhou

Abstract:

We used relatively modern tools to create binders using rfdiffusion/custom model ->proteinmpnn -> af initial guess and template targeting. For hotspots we used the given interface in the pdb, hydrophobic patches and also from mmgbsa. We optimized the sequence with in silico saturated mutagenesis on alphafold metrics. We found mmgbsa proposed the best hotspots and proteinmpnn struggled to generate good interface residues based on ipae.

Reproducibility:

0)hotspots based on hydrophobic patches and mmgbsa

1)Rfdiffusion (https://www.nature.com/articles/s41586-023-06415-8)

2) proteinmpnn (https://www.science.org/doi/10.1126/science.add2187)

3) alphafold with initial guess and template targeting (https://www.nature.com/articles/s41467-023-38328-5)

## 14. Furman Lab

Team Members:

Ora Furman, Julia Varga, Sarah Knapp

Abstract:

Our main design strategy was centered on the definition of promising starting fragments that form critical interactions with the receptor, which were then extended to form a stable minibinder. The starting fragments were determined using an adapted version of our PatchMAN^76^ peptide docking protocol, which searches for protein structures that contain structural motifs that are similar to patches on the receptor surface. Fragments that complement these structural motifs in these proteins are then extracted as candidate seeds, if they bind to the extracellular region.

Starting from these fragments, we used to RFdiffusion to build a stable minibinder scaffold around them (both with and without specifying hotspots based on available crystal structures, ProteinMPNN to design a matching sequence, and finally Alphafold2 to test refolding of the design into the target structure. For promising designs, we intensified the search using partial RFdiffusion. We also experimented with the newly published BindCraft protocol.

Promising designs were filtered by iPAE and rmsd, and then minimized with Rosetta^33^, scored with InterfaceAnalyzer and contact molecular surface area. Then, we ranked the designs according to ipTM, ipAE, contact molecular surface area and dGcross/dSASA*100 (measures that we and others found to correlate well with successful structure predictions and designs).

Reproducibility:

1. RFDiffusion with motif-scaffolding, starting from fragments defined using PatchMAN
2. Extract starting fragments: We ran PatchMAN[1] to find proteins with structural match to surface patches. From these matches we extracted complementing fragments that could be used as starting points for binder design. We masked the non-extracellular regions, as we did not want the binders to touch the membrane-embedded regions.
3. Build full binder starting from these fragments: We ran RFDiffusion[2] with motif-scaffolding and sequence inpainting of the extracted template in two different modes:
4. General, without specified hotspots
5. Targeted: with specified hotspots, defined using Rosetta alanine scanning[3]^77^ of crystal structures (Y161 and N171 of CD20)
6. Filter promising backbone candidates: From the backbone outputs, we removed those with high radius of gyration (>16A), and/or with significant contact of the membrane regions (>15% of binder residues), and with low contact to the extracellular regions (<20% of binder residues)
7. Design sequences for backbones: We ran ProteinMPNN using the ColabDesign[4] implementation (excluding cysteine, temperature 0.15), which also provided an interface to refold with AlphaFold2[5]. For refolding, we used templates for CD20, initial guess (as implemented in Benett et al.[6]), AlphaFold2-multimer-v3 model 1 and recycles of 3.
8. Select promising candidates: We removed designs with low predicted alignment errors (ipae > 10A), minimized them with Rosetta[7], and calculated Rosetta interface scores. We later further removed designs with rmsd > 3 between the template and the refolded structure.
9. Partial diffusion on accepted designs
10. To increase the diversity of the designed backbones, we performed additional partial diffusion using RFDiffusion. partial_T was set to 10, and 10 new backbones were generated from original designs that were selected after manual inspection of the best refolding designs. The same ProteinMPNN (but with temperature of 0.3) and AlphaFold2 refolding protocol was run, and the same cutoffs applied as before.
11. A second round of partial diffusion was conducted on the top 15 performing designs from the initial partial diffusion, following the same methodology as in part A.
12. BindCraft

BindCraft[8] was run until reaching 10 accepted designs with default parameters, using the provided Colab notebook. 2 designs with iptm values of equal or better of the methods described above were included in the final set.

1. Ranking candidates:

For ranking the sequences, we calculated the rank of each design by four metrics: ipTM, ipAE, contact molecular surface area and dGcross/dSASAx100. We summed up these ranks for each designed sequence, and sorted the designs according to them.

## 15. LBM

Team Members:

Fernando Meireles, Parth Bibekar, Benedikt Singer

Abstract:

Knowing that CD20 is hard target to design binders against, we decided to perform some Molecular Dynamics on the target to get a wider variety of starting structures. We also assumed that flexible chains comprising the epitope might have different conformations one of them maybe being a bit wider allowing for a bigger binding site. After clustering the MD trajectories a total of 29 cluster remained from which 8 were selected based on a more open confirmation. A subsequent RFDiffusion run was started to generate a total of 5,000 structures where either randomly the true target structure or an MD-derived structure was used. Additionally from a set of hotspots, two hotspots were randomly sampled.

Inverse folding was performed using either ProteinMPNN with or CARBonAra^25^ with their respective default parameters. For each RfDiffusion-generated structure 5 sequences were generated.

After this localcolabold’s version of AlphaFold2 was applied followed by Alphafold-multimer where the true structure of the target was provided. The thus produced metrics (pAE, pLDDT) were used for the submission.

(A short remark about the delayed hand-in: After the deadline extension we had some internal misunderstandings of the timezones. We hope that our submission will still be considered)

Reproducibility:

- Target structures: 6VJA (https://www.science.org/doi/10.1126/science.aaz9356)
- MD-parametrisation: GROMACS 2024.02 (https://www.sciencedirect.com/science/article/pii/S2352711015000059)
- Structure-binder generation: RFDiffusion (https://www.nature.com/articles/s41586-023-06415-8)
- Inverse folding:

- Protein-MPNN (https://doi.org/10.1126%2Fscience.add2187)
- CARBonAra (https://www.biorxiv.org/content/10.1101/2023.06.19.545381v1.full.pdf)
- Validation:

- Colabfold (https://www.nature.com/articles/s41592-022-01488-1)

- Localcolabfold (https://github.com/YoshitakaMo/localcolabfold)
- AlphaFold2 (https://www.nature.com/articles/s41586-021-03819-2)
- AlphaFold-Multimer (https://www.biorxiv.org/content/10.1101/2021.10.04.463034v1)

## 16. Molecule Masters

Team Members:

Karl Lundquist, Dion Whitehead, Arjun Singh, Abel Gurung, Amardeep Singh

Abstract:

We developed a computational pipeline to design protein binders targeting CD20. Starting with RFDiffusion, we generated initial binder structures and optimized their sequences using ProteinMPNN alongside Rosetta FastRelax to optimize for folding into the designed structure as well as complementarity with the extracellular region of CD20. To do this we carried out a series of rounds in which we balanced exploration and exploitation by carrying the top performers into subsequent rounds and creating new binders in each round as well. In each round, we predicted the 3D structures using AlphaFold2. We removed binders that were poorly predicted by AF2 and also those that would protrude into the membrane after proper alignment. In order to accurately assess the binding stability, we used an energy minimization and molecular dynamics simulation procedure to relax the side-chains. Using this equilibrated structure we evaluated binding stability by calculating the total energy and the non-bonded interaction energy between the binder and CD20. We also estimated binding free energies (ΔG) using Prodigy. We also calculated binder radius of gyration in order to preference compact designs. Our scoring function combined binder radius of gyration, interaction energy, and ΔG, adjusted by total energy, to effectively rank the binders. Designs meeting our criteria for energy thresholds and RMSD advanced through the rounds, improving their stability and affinity. Finally, we consolidated and re-ranked the top binders from all rounds. By integrating RFDiffusion for structure generation and ProteinMPNN with FastRelax for sequence design, along with careful filtering steps, our pipeline efficiently produces high-affinity binders specific to the extracellular region of CD20.

Reproducibility:

- RFDiffusion to generate initial protein binder structures

- Watson, J.L., Juergens, D., Bennett, N.R. et al. De novo design of protein structure and function with RFdiffusion. Nature 620, 1089–1100 (2023). https://doi.org/10.1038/s41586-023-06415-8

- ProteinMPNN, FastRelax, AF2 procedure for binder design

- Bennett, N.R., Coventry, B., Goreshnik, I. et al. Improving de novo protein binder design with deep learning. Nat Commun 14, 2625 (2023). https://doi.org/10.1038/s41467-023-38328-5

- Dauparas, J., Anishchenko, I., Bennett, N., et al. (2022). Robust deep learning–based protein sequence design using ProteinMPNN. Science, 378(6615), 49–56. https://doi.org/10.1126/science.add2187

- Jumper, J., Evans, R., Pritzel, A., et al. (2021). Highly accurate protein structure prediction with AlphaFold. Nature, 596(7873), 583–589. https://doi.org/10.1038/s41586-021-03819-2

- Molecular Dynamics for side chain relaxation^28^

- Eastman, P., Swails, J., Chodera, J.D., et al. (2017). OpenMM 7: Rapid development of high performance algorithms for molecular dynamics. PLOS Computational Biology, 13(7), e1005659. https://doi.org/10.1371/journal.pcbi.1005659

- Prodigy for binding free energy calculation^36,78^

- Vangone, A., & Bonvin, A.M.J.J. (2015). Contacts-based prediction of binding affinity in protein–protein complexes. eLife, 4, e07454. https://doi.org/10.7554/eLife.07454

- Xue, L.C., Rodrigues, J.P., Kastritis, P.L., Bonvin, A.M.J.J., & Vangone, A. (2016). PRODIGY: a web server for predicting the binding affinity of protein–protein complexes. Bioinformatics, 32(23), 3676–3678. https://doi.org/10.1093/bioinformatics/btw514

## 17. N6 / Team Anonymous

Team Members:

Nicole Chiang

Abstract:

We used an approach combing domain knowledge and machine learning. We generate a library of peptide sequences incorporating known binding motifs, emphasizing the integration of an aromatic cradle composed of tryptophan and tyrosine residues, along with hydrogen bond donors like asparagine or serine. Each sequence is evaluated using ESMFold to determine its pLDDT score, and sequences with low scores are filtered out to retain those likely to exhibit better structural integrity. We hope this simple but combined approach may address the challenges of stability and folding in peptide binders while ensuring that specific structural features enhance binding affinity.

Reproducibility:

1. Sequence Generation:

* Generate peptide sequences incorporating known binding motifs based on scaffolds that are known to interact with proline rich domains (which is sort of present in the extracellular loops of CD20), such as SH3 and WW domain^79^.

* Design the sequences to include an “aromatic cradle” with tryptophan and tyrosine residues for enhanced recognition of proline residues.

1. Incorporation of Structural Features:

*Integrate additional hydrogen bond donors, such as asparagine or serine, to interact with backbone carbonyl groups, improving stability.

1. Charged Residue Complementation:

*Introduce complementary charged residues around the binding motifs to enhance binding specificity.

1. Model Selection:

* Utilize ESMFold, a protein structure prediction model, to evaluate the generated peptide sequences.

1. pLDDT Score Calculation:

* Input each generated sequence into ESMFold to obtain pLDDT scores, which indicate structural integrity.

1. Filtering Based on pLDDT Scores:

*Filter the generated sequences by retaining only those with higher pLDDT scores, ensuring a focus on sequences likely to exhibit better structural stability^80^.

## 18. Nucleate UK London

Team Members:

Jakub Lála, David Miller, Stefano Angioletti-Uberti

Abstract:

We employed the high-level programming language from (Hie et al. 2022)^81^, later implemented as a part of the BAGEL package^46^. Two design strategies were employed, detailed below.

First, full 80-mer sequences were designed against the CD20 dimer. After the Monte Carlo (MC) optimization, SolubleMPNN^24^ was employed to generate alternative sequences from the same backbones. Specifically, we retrieved the top-100 designs from the MC optimization based on the final value of the objective/energy function. These final sequences were not de-duplicated, so the total number of unique designs from this part is not 100. We then further repainted each backbone many times with SolubleMPNN, and took the top-100 based on the confidence score in SolubleMPNN. In total, this gave 200 sequence submissions from the first design strategy.

Second, 20-mer segments were designed using the MC optimization against the CD20 monomer epitope alone. RFDiffusion^16^ was then used to design the 40-mer linker between the two monomer binders, where we alternated various monomer binders with one another (e.g., a monomer binder A and B would be combined as AA, AB, BB). The linkers alone were then inpainted with SolubleMPNN to get the full 80-mer sequence. The monomer binder segments were thus kept conserved from the ESMFold-driven MC optimization. As before, we took the top-300 designs based on the confidence score from SolubleMPNN.

We also validated some of our best designs by visually inspecting 10 ns molecular dynamics, but due to a lack of time did not use this extensively to do any ranking or filtering.

Reproducibility:

1. ESMFold optimization design^81^
2. Earlier version of BAGEL^46^
3. SolubleMPNN alternative sequence design^24^
4. MD Validation with OpenMM^28^

## 19. Perez Lab Gators

Team Members:

Jokent Gaza, Alberto Perez, Bhumika Singh, Yisel Martinez Noa, Qianchen Liu, Rangana De Silva

Abstract:

Binders for the CD20 extracellular domain were generated using the RFDiffusion to ProteinMPNN-FastRelax pipeline. As an orthogonal assay, the generated sequences, along with the CD20 dimer, were folded using AlphaFold2 Initial Guess. Our initial plan was to use the binding epitopes of three antibodies (rituximab, ofatumumab, and obinutuzumab) as initial motifs for RFDiffusion. Inpainting would then generate a scaffold that preserves the binding epitope and/or interacts with other residues in the CD20 binding domain. However, although the pipeline produced stable monomers, the predictions for the complex were suboptimal. An initial cut-off of pAE < 15 was set, and none of the 18000 designs (6000 per antibody) met this threshold.

For plan B, we decided to start from a completely de novo design strategy. Out of the 3000 designs, only 5 passed the pAE threshold. Of these, only two (called DN1 and DN2) had all the residues in the extracellular region of CD20. For both DN1 and DN2, we extracted the residues that directly interact with CD20. We then performed another round of RFDiffusion to generate 1000 designs that have a scaffold for these residues. For DN1, we obtained only one good design, while DN2 generated 65. We then extended the number of designs for DN2 to 8000, resulting in 211 designs. These involved optimizing either the entire sequence or just the non-binding residues using ProteinMPNN. We also optimized the non-binding residues of DN1,giving a final list of 223 sequences. A quick sanity test was then performed to test if these are stable proteins. Specifically, we performed AF2 predictions for both the monomer and the full CAR construct. We surmised that good binders should retain their structure both as a monomer and in the full CAR (RMSD < 2 Å) and should be compact (RoG < 14 Å). From these, we were able to further reduce the number of designs to 193 (12 from DN1 and 181 from DN2).

To reach the 500-sequence target, we tried optimizing the final 181 sequences from DN2 using proteinMPNN. We performed three runs, for a total of new 543 sequences. Of these sequences, 166 passed the pAE filter. We also revisited the initial predictions with antibody epitopes and de novo designs. We took all the sequences that have pLDDT > 85 for the binder, and performed the same sanity test. We then ranked the designs based on their pLDDT score as a monomer to decide which of them will be included in the submission. To introduce diversity in our submission, we decided to prioritize the results from the antibody epitopes, as they have more varied topology. In total, the final 500 sequences included 359 good binders, 60 stable monomers from antibody epitopes, and 81 stable monomers from de novo predictions.

Our github page can be found in: https://github.com/PDNALab/BioML

Reproducibility:

1. Extracted the binding epitopes of the following antibodies – rituximab (6VJA), ofatumumab (6Y92), and obinutuzumab (6Y97).
2. Generated 6000 designs for each antibody—3000 using the Complex_base model of RFDiffusion and another 3000 using the Complex_beta model.^82^ (1-5)
3. Filtered the designs using a pAE threshold (pAE < 15), but no designs met the criteria.
4. Generated 3000 de novo designs for the CD20 structure from 6VJA using the Complex_beta model with reduced noise (i.e., with denoiser.noise_scale_ca and denoiser.noise_scale_frame set to zero) during sampling. Only five designs passed the filter, and only two (called DN1 and DN2) were considered reasonable binders.
5. Extracted the binding epitope of DN1 and generated 1000 additional designs. Only one design passed the threshold.
6. Tried to optimize the non-binding residues of DN1 using proteinMPNN. The final list of designs from DN1 is 12.
7. Extracted the binding epitope of DN2 and generated 1000 additional designs, resulting in 65 designs. Extended this to 8000 designs, to get a list of 211 DN2 designs.
8. The combined total of designs from DN1 and DN2 was 223. A sanity test was conducted by comparing the binder as a monomer and in the full CAR, using RMSD < 2 Å and a radius of gyration < 14 Å as criteria. The final list of DN1 and DN2 designs was reduced to 193.
9. Optimized the 181 sequences from DN2, resulting in an additional 166 sequences.
10. Performed the same sanity test on all designs (both antibody-based and de novo) with pLDDT > 85. For the final sequence list, we prioritized designs with antibody epitopes and pLDDT > 90.
11. The final list of sequences included 359 good binders, 60 stable monomers from antibody epitopes, and 81 stable monomers from de novo predictions.

## 20. Perforators

Team Members:

Vinayak Annapure, Vikrant Parmar, Huan Koh, Saeideh Moradvandi, Yizhen Zheng

Abstract:

We analyzed the known binders against CD3 and used those residues as a constraint for diffusing potential binders for CD20. We further used ProteinMPNN to solve the reverse folding problem for the designed binders and used ESMFolds confidence score to select the potential binders

Reproducibility:

N/A

## 24. The Physicist

Team Members:

Rohit Satija

Abstract:

Solved structures of CD20 in complex with antibodies are available on the PDB: [6VJA](https://www.rcsb.org/structure/6vja), [6Y92](https://www.rcsb.org/structure/6Y92), and [6Y97](https://www.rcsb.org/structure/6Y97). In all three cases, the hotspot region of CD20 present inside the binding region is the short macrocyclic polypeptide connected by a disulfide bridge ‘NCEPANPSEKNSPSTQYC’ (166-184, chain D, 6vja). For this reason, we decided to focus on this epitope as our binder target. We chose to take a rational design approach to pick fragments of antibodies closest to the target epitope in the PDB structures using a nearest neighbor search algorithm. These sequences were ranked according to a perplexity score computed from embeddings calculated using an open source 33 layer ESM-2 model with 650M params, originally trained on UniRef50. Top 500 sequences were submitted for experimental validation. This work resulted in a number of [jupyter notebooks](https://github.com/rohitium/bioml-challenge-2024/tree/main/notebooks) and a [rational protein design package](https://github.com/rohitium/rational_protein_design) that could be useful for future protein design studies.

Reproducibility:

To replicate the final result, i.e.

[submission.fasta](https://github.com/rohitium/bioml-challenge-2024/blob/main/results/submission.fasta), follow these steps:

1. **Binder Design:** This step is accomplished using the [rational_design.ipynb](https://github.com/rohitium/bioml-challenge-2024/blob/main/note books/rational_design.ipynb) notebook. Briefly, the python package [rational_protein_design](https://pypi.org/project/rational-protein-design/) contains a class called ‘BinderDesigner’ that takes in the following attributes:

* ‘pdb_filè: Path to an existing PDB structure of the bound complex (Str)

* ‘chain_id’: Chain ID of the target (Str)

* ‘target_residues_rangè: Range of residues on Target chain to design binders for (Tuple(Int))

* ‘neighbor_chain_id’: Chain ID of the candidate bound molecule (Str)

* ‘N’: Length of binder (Int)

* ‘num_seq’: Number of binders to design (Int)

Provided these attributes to an object of the ‘BinderDesigner’ class, the ‘design_binder()’ method runs all the functions necessary to design binders for the target sequence and store them in a provided fasta file path.

1. **Rank Binders:** This step is accomplished using the [compute_embeddings.ipynb](https://github.com/rohitium/bioml-challenge-2024/blob/ma in/notebooks/compute_embeddings.ipynb) notebook. Briefly, we create embeddings for all candidate binder sequences produced in the previous step using Facebook’s ‘esm2_t33_650M_UR50D’ model (https://github.com/facebookresearch/esm?tab=readme-ov-file#main-models-you-should-use-). Next, we compute perplexity for each individual sequence (https://en.wikipedia.org/wiki/Perplexity) that is a metric describing how well the model predicts the existence of the sequence. We rank order the binder sequences using this score and save the top 500 sequences to file.

## 22. Picaso Protein

Team Members:

Elias Sanchez, Nuria Mitjavila, Sofiia Hoian, Alejandro Diaz, Marko Ludaic, Jinling Huang, Nader Ibrahim, Yong Youn Kwon

Abstract:

We aimed to design high-affinity binders targeting Rituximab’s epitope (C171-174) using three computational strategies. First, we used nanobodies and language models with Raygun to generate optimized binder sequences of 80 aa tailored to Rituximab’s epitope. Second, we explored novel sequence spaces through de novo design without existing templates. Third, we generated cyclic peptides and replaced Rituximab’s CDR2 and CDR3 regions with random peptide pairs, refining these using custom metrics. A rules-based mutation strategy was applied, mutating residues based on interaction interface and structural metrics like pLDDT and iPAE. AlphaFold2 Multimer was used to model and select optimal candidates after each mutation round.

Reproducibility:

-We aimed to design high-affinity binders targeting Rituximab’s epitope (C171-174) using three distinct computational strategies. [1]^83^

-First, we utilized existing nanobodies and language models with Raygun to generate optimal binder sequences and reduce the size to 80 aa, specifically tailored to Rituximab’s epitope on CD20. [2-4]^22,84,85^

-Second, we performed de novo design of binders to explore novel sequence spaces without relying on existing templates.

-Third, we generated cyclic peptides with high predicted Local Distance Difference Test (pLDDT) scores and replaced the CDR2 and CDR3 regions of Rituximab’s heavy chain with randomly selected peptide pairs, followed by sequence refinement using custom scripts and metrics to select optimal candidates. [6]

-To optimize these initial sequences, we applied a rules-based mutation strategy: residues at the interaction interface were mutated at a low rate favoring substitutions with positive BLOSUM scores; neighboring residues were subjected to higher mutation rates based on their pLDDT and inter-residue Predicted Aligned Error (iPAE) values; non-interacting and non-neighboring residues experienced even higher mutation rates aimed at increasing their pLDDT values. Neighboring and interaction residues where determined with getcontacts.[5]

-After each round of in silico mutations, subsets of offspring sequences were generated and modeled using AlphaFold2 Multimer to predict their three-dimensional structures. This iterative process applied evolutionary pressure, selecting sequences with high confidence according to both pLDDT and iPAE metrics, ultimately refining candidate binders toward optimal interaction with Rituximab’s epitope.

Get_contacts: https://github.com/getcontacts/getcontacts/tree/master

## 23. Schoeder Lab

Team Members:

Moritz Ertelt, Max Beining, Dominic Rieger, Dieter Hoffmann, Tom Schlegel, Max Lingner, Vivian Haas

Abstract:

The workflow initiated with the crystal structure (PDB: 6VJA) and molecular dynamics (MD) simulations to model the outer loops. Principal component analysis (PCA) identified three predominant loop configurations, and both, the original crystal structure and a representative structure of each cluster, were selected for subsequent steps. Using RFdiffusion, binders were generated by targeting trimmed extracellular domains in dimeric configurations. Hotspot regions, including crucial interaction residues from CD20 and Rituximab, guiding peptide generation and de-novo binder design. Filtering criteria such as contact order, shape complementarity, and secondary structure were applied to identify potential high-quality backbones suitable for sequence design. Sequence design was performed using ProteinMPNN across different temperature levels, with AlphaFold-based target-binder-complex predictions supporting scoring and selection of the designs. Final binders were iteratively refined using metrics like pae_interaction, pLDDT, Rosetta’s dG_separated, and additional LIA and LIS filters. In parallel, an AlphaFold hallucination approach was employed to generate peptides of varying lengths. These peptides were scaffolded using RFdiffusion and optimized iteratively for binding against CD20. In addition, peptide repeats with linker regions were introduced to enhance avidity-like binding properties.

Reproducibility:

Crystal structure (PDB: 6VJA) and an MD simulation for the outer loops as a starting point, MD + PCA results in three main loop configurations (use only one cluster representative for further steps)

Using RFdiffusion [1] in various approaches, only working on trimmed input structures (extracellular part, dimer configuration)

Generating binders by providing different hotspot regions (e.g., the original key interaction residues from CD20 with Rituximab [3], unpublished surface clustering algorithm)

Peptide generation + scaffolding De-novo binder design

Perform filtering by pyrosetta and specific cutoffs for contact order, shape complementarity, secondary structure etc.

ProteinMPNN [4] for sequence generation using different temperatures and Alphafold initial guess for target-binder prediction and scoring based on the pipeline from [5]

Final designs were filtered using pae_interaction, plddt and Rosetta [6]^34^ dG_separated In addition to that, some designs were filtered using the metrics suggested by [7], namely LIA and LIS

Binders were optimized iteratively with ProteinMPNN and Alphafold initial guess by using good scoring models as inputs or partially diffused variations of them

Using an inversion of the AlphaFold structure prediction network [2] and their protocol to generate peptides of different lengths generated peptides were scaffolded using RFdiffusion

Peptide repeats with linker regions were designed to achieve avidity-like binding

## 24. SNU LCDD

Team Members:

Jinwoong Song, Hakyung Lee, Haelyn Kim

Abstract:

We applied multi-track for successful binder design.

For backbone design, we applied 1) de novo, 2) grafting antibody CDR loops 3) grafting CDRH3 only.

We generated the protein sequences with proteinMPNN and filtered out the sequences with Rosetta modeling and scoring.

The remained sequences were modeled the complex structure with AlphaFold2.

Lastly, We run short Molecular Dynamics simulation for the complex structure in order to sort the designs in binding affinity to CD20.

Reproducibility:

RFdiffusion

ProteinMPNN

Rosetta

AlphaFold2

Molecular Dynamics

## 25. Tumor Inhibitors

Team Members:

Ashish Makani, Cianna Calia, Rana A. Barghout, Matthew Williams, Javier Marchena-Hurtado, Vivian Chu

Abstract:

Expanding the set of known protein binders to human CD20 holds promise for medical applications such as CAR T-cell therapies for B-cell malignancies. However, CD20 presents a challenging target for de novo binder design for multiple reasons: The extracellular loops (ECL1/ECL2) protruding out of the cell membrane provide relatively small surfaces for binding and contain many hydrophilic residues [1], and judging by the slightly lower pLDDT in these loops in CD20’s AlphaFoldDB structure [2, 3]^87,88^, the exposed regions may also have some degree of flexibility. We therefore constructed a multi-tiered approach to attempt to maximize the success rate of CD20 binders we generated with state-of-the-art deep learning tools. First, we used RFdiffusion’s [4] motif scaffolding functionality to create a minimal target structure consisting of segments of both CD20 monomers from PDB 6VJA [1], united into a single asymmetrical chain by two diffused glycine linkers. With this structure as the provided target, we used RFdiffusion’s “beta” complex parameters to generate over 25,000 80-residue binder backbones. To allow us to generate many sequences per backbone without wasting time on low-quality designs, we filtered our backbones using Biopython [5] to ensure contact with Tyr161 in CD20’s ECL2, to prohibit contact with the nearest diffused linker, and to avoid binders with excessively long helices and/or too few target contacts. We also eliminated any backbones incompatible with the cell membrane by aligning each target-binder complex to a membrane-embedded structure of CD20 from MemProtMD [6] and discarding any backbones with more than 10 heavy-atom clashes between the binder and the lipids. For all backbones passing this filtering, we used the ColabDesign framework [7] to run ProteinMPNN [8] to generate 64 sequences each with a sampling temperature of 0.000001, followed by screening with AlphaFold2 (using model_1_ptm with initial guess, target templating, three recycles, and no MSAs) [9, 10]. Because we found binders with pae_interaction < 10 to be rare, we prioritized further exploration of the sequence and structure space surrounding “near hits” that had pae_interaction < 22 and RMSD < 8 Å with respect to the designed structure; for all backbones giving one or more near-hit sequences we generated and screened 64 additional sequences with a sampling temperature of 0.1, and for some promising candidates we generated still more sequences and in several cases also diversified the binder backbone with partial diffusion. With these methods we compiled 1,323 binder sequences with pae_interaction < 22 and RMSD < 8 Å, which we ranked by pae_interaction (and re-ranked by RMSD within pae_interaction windows of 1.0). After discarding those with pLDDT < 80, those with long alanine stretches, and those within 20 amino acids of higher ranking candidates, 868 sequences remained. Our submission contains the highest ranking 500 of these remaining sequences.

Reproducibility:

- RFdiffusion [4] to create minimal target structure from PDB 6VJA [1]

- RFdiffusion [4] to generate binder backbones

- Backbone filtering with Biopython [5]^89^ and check for membrane clashes using structure from MemProtMD [6]

- ProteinMPNN [8] to generate sequences (via ColabDesign [7])

- AlphaFold2 [9] with initial guess [10] to screen designs (via ColabDesign [7])

- Additional sequence generation / partial diffusion for best designs

- Rank best sequences by pae_interaction and RMSD

## 26. Undergraduate Design

Team Members:

Max Witwer, Nathaniel Greenwood, Jie Chen, Hannah Stewart, Elliot Cole

Abstract:

This study aims to develop high-affinity mini-binders for the extracellular domain of CD20. CD20, a transmembrane protein overexpressed in many B cell malignancies, is a prime target for novel therapeutics. CAR T-cell therapy, an immunotherapy technique, involves engineering a patient’s T-cells to express chimeric antigen receptors (CARs) that recognize antigens like CD20, enabling them to destroy malignant B cells. Computational protein design has enabled the creation of novel protein backbones that can interact with a range of structures, facilitating the development of mini-binders to enhance targeting in CAR T-cell therapy. However, current computational limitations and CD20’s unique structure make it a challenging binding problem due to the hydrophilic nature of its extracellular domain. Current design methods rely on hydrophobic residues for energetically favorable binders, leading to two challenges: first, unconstrained design generates proteins that bind mainly to the hydrophobic transmembrane domain; second, binders to the extracellular region have low predicted binding scores by tools like AlphaFold2. To address these issues, a design campaign used RFdiffusion, ProteinMPNN, and AlphaFold2. CD20’s structure (PDB entry 6VJA) was downloaded, and the rituximab Fab was deleted using PyMOL. Initially, 80-residue “seed structures” were created in reasonable locations, approximating previously designed Fabs like rituximab (6VJA) and obinutuzumab (6Y9A).

To alleviate these issues, the structure was mutated to drive the AI models to produce better results. First, phospholipid-facing residues in the transmembrane domain were mutated to aspartate, producing a pseudo-solubilized protein to discourage AI models from designing transmembrane region binders. Next, six key residues (GLU 174, SER 177, SER 179 on each CD20 dimer) were mutated to LEU 174, VAL 177, VAL 179. High-temperature partial diffusion (60-100 noise steps) guided the model to bind to the desired domain, producing high-affinity CD20 binders, AlphaFold2 interface predicted aligned error (PAE). To create mini-binders for the native target, an evolutionary algorithm was used to slowly mutate the 80 residue structure toward more favorable results. The structure of CD20 between generations was slowly mutated back to its native hydrophilic form. Designed backbones were mutated using low-temperature partial diffusion (15-18 noise steps) and sequences were mutated with ProteinMPNN; the produced designs were selected for interface root mean squared deviation (RMSD) and interface PAE. This was repeated across nine generations. A final refinement step was performed using predictions from an extended version of AlphaFold2 dubbed AlphaFold multimer, which is designed to specifically predict protein-protein interactions. The final tenth round of recycling was performed directly on multimer predictions and the resulting sequences were ranked based on AlphaFold multimer interface PAE; additionally, cutoffs were imposed for the interface RMSD, binder predicted local distance difference test (pLDDT) and interface predicted template modeling (TM) score. The final ranked pool of results contains 500 predicted sequences.

Reproducibility:

Steps:

1. Use crystal structure 6VJA, remove rituximab and portions of the transmembrane domain (TD) that are far from the extracellular domain. Mutate any remaining outward facing TD residues to aspartate. Mutate hydrophilic residues in the extracellular binding pocket to hydrophobic residues with similar geometries.
2. Use PyMol builder to build four helices in the extracellular binding pocket
3. Use partial RFdiffusion[1] to make these helices into biochemically reasonable backbones. Use high partial T to sample diverse backbones that are not simply a four helical bundle. Use ProteinMPNN[2] and AlphaFold2 initial guess[3].
4. Choose best designs[4]^90^ based on pae_interaction, mutate 1-2 residues back, recycle those designs, and repeat until the native structure is recovered.
5. Run AlphaFold multimer[5]^91^ on the predicted structures and perform a final round of partial diffusion, rank the final designs based on AlphaFold multimer interface PAE and RMSD with cutoffs for desired metrics.

## 27. Virtue Therapeutics

Team Members:

Rajat Punia, Akshay Chenna

Abstract:

Used BindCraft model for binder designs with default filter parameters.

Reproducibility:

1. Receptor residues that make H-bond with the antibody in PDB 6Y92 are taken as hotspot residues.
2. PDB 6Y92 is chosen to decide as hotspot to design binders that can bind to either chain of the receptor and not the interface. This results in small reduction in entropic penalty of binding.
3. Binders are designed with default settings using BindCraft. None of the design was able to pass the default filters. Best designs from the trajectory are used for submission.

## 28. Zist Rayanesh

Team Members:

Seyed Alireza Hashemi, Kaveh Nasrollahi, Fatemeh Nasiri

Abstract:

Design begins by analyzing the target molecule to identify binding hotspots. At this initial stage, hotspots are selected based on hydrophobic patches and known antibody epitope sites involved in binding. For each of the six identified hotspots, 100 binders, ranging from 12 to 18 residues, are designed using RFdiffusion. This is followed by generating 8 sequences per binder with MPNN+Fastrelax and then using AlphaFold-Multimer for structure prediction. Validation of binders is based on metrics like binding energy and iPAE, and the best hotspot is chosen for further optimization. This step was constrained by time and computational resources.

After selecting a pool of promising mini-binders, the next phase involves designing scaffolds for them using RFdiffusion. Subsequently, the binders are dimerized to target both accessible sides of the molecule, provided it has a symmetrical structure.

The final designs must conform to a standard format with 80 residues. Binders requiring fewer than six residues to bridge gaps are connected using a Gly-Gly-Ser linker. For larger gaps, the RFdiffusion pipeline is run again to optimize the design.

308 sequences were submitted due to the limitation of computational unit (i3 12100f + RTX 3060 + 16 GB of RAM)

Reproducibility:

- Finding Hotspots (https://www.acrobiosystems.com/L-965-CD20.html?gad_source=1)
- RFdiffusion for binder design (https://github.com/RosettaCommons/RFdiffusion)
- MPNN for designing sequences (https://github.com/dauparas/ProteinMPNN)
- ColabFold (Local + MMseq server) (https://github.com/sokrypton/ColabFold)
- Pyrosetta as a filtering step (https://docs.rosettacommons.org/docs/latest/application_documentation/analysis/interf ace-analyzer)

